# Multi-scale place fields provide a predictive basis for hierarchical control

**DOI:** 10.64898/2026.10.28.762461

**Authors:** Changmin Yu, Shir R. Maimon, P. Dylan Rich, Tamir Eliav, Liora Las, Albert K. Lee, Nachum Ulanovsky, Máté Lengyel

## Abstract

Hippocampal place cells typically exhibit unimodal spatial selectivity in small enclosures, but express multiple, irregularly spaced fields in environments of naturalistic scale. However, existing predictive-map theories of the hippocampus do not scale well to large environments and do not account for these heterogeneous place field characteristics. Here, we introduce the factorised hierarchical successor representation (fHSR), a predictive map that supports hierarchical control using subgoal-directed subroutines. We show that the fHSR allows efficient generalisation across tasks, scaling favourably with environmental size, and it reproduces key statistical and topological characteristics of multi-scale CA1 place cell activity in rats and bats. Analyses of experimental data further confirmed a novel prediction of the fHSR: population firing rates exhibit spatial periodicity with a characteristic wavelength independent of environmental size. These results identify the integration of hierarchical control and predictive representations as a normative principle that unifies efficient navigation with the spatial organisation of hippocampal activity.

---

The hippocampus is thought to provide a cognitive map that supports flexible navigation and planning ^1,2^. An influential normative account posits that this map is predictive: place cells encode not merely where an animal is, but where it expects to be in the future ^3^. Such a predictive map is formalised as the successor representation (SR) ^4^, and explains various properties of place cell tuning curves that remain fundamentally unimodal in typical enclosures used in laboratory studies ^2,5,6^. The SR has also been shown to provide a neural representation that faithfully clusters task states according to the community structure of their underlying relational graph – as found experimentally in neuroimaging studies of the hippocampus ^3,7^.

However, the explanatory power of SR for neural representations in the hippocampus has only been established in small enclosures, in which most place cells have a single compact firing field, resulting in unimodal spatial tuning. This seems to break down in environments that animals navigate under naturalistic conditions that can be orders of magnitude larger. For example, radio-tracking studies of free-ranging rats have documented home ranges extending over hundreds of metres ^8^, while Egyptian fruit bats commute tens of kilometres each night between their roost and foraging sites ^9^. In contrast to the textbook picture of unimodal place-cell spatial tuning, rat and bat CA1 place cells express multiple, irregularly spaced place fields in environments of naturalistic scale ^10,11,12,13,14^. In particular, the number of place fields in large environments exhibits an overdispersed distribution across cells ^11,15^, with fields of different sizes coexisting within single cells ^12,14^. These properties are at odds with the simple place fields predicted by SR and instead have been suggested to arise from a random process without fulfilling any particular computational goal ^16^. Yet, even in large environments, not all is random in place cell statistics: the population activity of place cells bears the specific topological signature of a hyperbolic rather than a Euclidean geometry ^17^, and several measures of place cell activity exhibit a lawful dependence on the size of the environment ^11,12,14^. Here we provide a unifying explanation of all these aspects of heterogeneous place-cell spatial tuning across different environmental sizes. Our account is based on a single computational principle: the construction of a generalisable and scalable predictive map.

Our approach starts from recognising two computational limitations of the standard SR. First, a predictive map with a single spatial scale faces a trade-off in large environments: broad fields support long-range prediction but blur the local estimate of position and assign similar representations to distant locations (spatial aliasing) ^10,18^, whereas narrow fields preserve local precision but face a capacity limitation when a fixed population must cover a large space. This trade-off cannot be flexibly adjusted under the standard SR, because its spatial scale is fixed by a single discount factor. A second, related, limitation is that the SR assumes that actions are chosen at a fixed temporal resolution. This makes planning in large environments unfeasible. Instead, in large environments, using temporally extended, subgoal-directed subroutines (called “options”), such as “go to the end of the corridor”, have been shown to greatly improve the efficiency of planning ^19^. Indeed, animals do not traverse or plan in large environments one step at a time: their behaviour is organised hierarchically, with long trajectories decomposed into segments directed at subgoals such as doorways, landmarks or the ends of corridors ^20,21,22^. However, it has been unclear how predictive maps might support such hierarchical control.

To overcome the computational limitations of the standard SR, we developed the factorised hierarchical successor representation (fHSR) ^23^. The fHSR generalises the construction of a predictive map by two innovations: (1) it extends to hierarchical behaviour, which combines primitive one-step actions with temporally extended options, and (2) uses a factorisation that is provably optimal for task transfer. As a result, we show that, unlike the standard SR, the fHSR both supports flexible generalisation across different tasks in the same environment and scales favourably to large environments. Critically, the fHSR provides a set of features that predict the environmental size-dependent multi-scale organisation of place cell activity of rat and bat CA1 neurons, at both the single-cell and population levels. In addition, the fHSR made a novel prediction about population firing rates showing spatial periodicity in simple linear environments, which we confirmed in experimental data. Our results thus point to a key role of multi-scale place fields in establishing a predictive map for hierarchically organised, flexible behaviour.

## Results

### The HSR and its factorisation support simultaneously scalable and generalisable learning

Under the SR framework, the value of a state (expected discounted future return from the state ^25^) is the sum of the future occupancies of all states predicted under the current behavioural policy, each weighted by the reward available there ^4^ (Equation 5). The SR therefore supports efficient generalisation across tasks: when the task or reward changes, the updated value function can be recomputed immediately by re-weighting the predictive map with the new reward function, rather than being relearned from scratch ^4,26^. To demonstrate this, we trained reinforcement-learning agents in a four-room environment on a series of 20 tasks, each requiring navigation from the same start location to a different goal (Figure 1a). All agents learned action values as a linear readout of their state representation ^24^. The representation was learned incrementally across tasks. For SR agents, it was the running average of the SRs under the policies learned for the completed tasks, an estimate of the task-averaged SR (analogous procedures were used for all other agents; Methods). All agents started from one-hot state features in the first task. Because the readout weights were re-initialised for every task, only the representation carried information from one task to the next, so any improvement across tasks measures how well a representation supports transfer to new tasks (Extended Data Figure 1a,b). SR features conferred a clear advantage in transfer: SR agents learned to reach later goals in far fewer training episodes, whereas naive agents with one-hot state encoding did not improve across tasks (Figure 1d,f,m, left panels).

**Figure 1:**
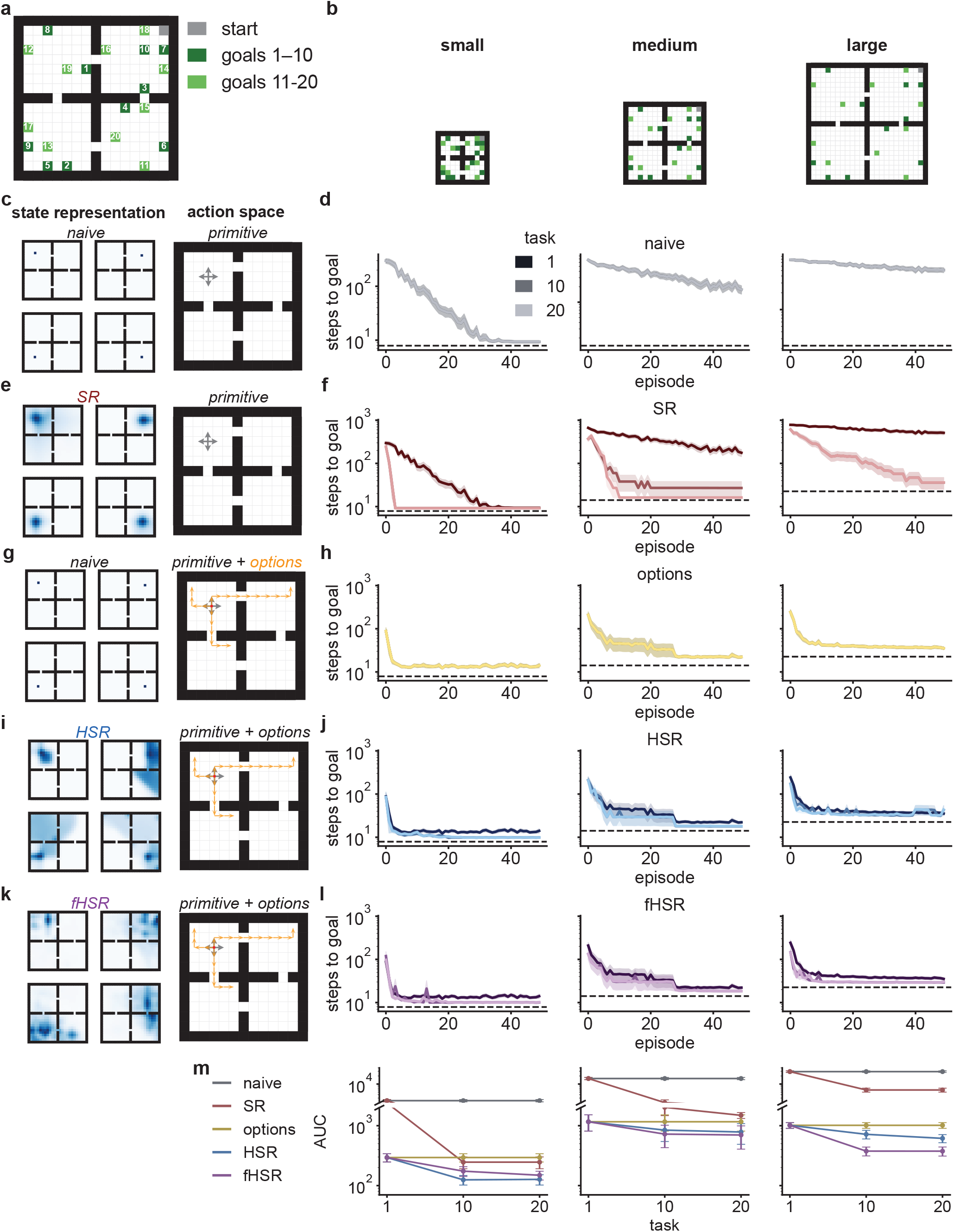
The hierarchical successor representation (HSR) and its factorisation (fHSR) support learning that both generalises across goals and scales with environment size. a. Four-room navigation task. From a fixed start state (grey), agents navigate to a different goal in each of a series of tasks. Numbers label the 20 goal locations (dark green, goals 1-10; light green, goals 11-20). **b**. Small, medium and large versions of the environment, with the same room-and-doorway topology and room-relative goal (green) and start (grey) locations. **c, e, g, i, k**. Example state features (left) and action spaces (right) of five agents: one-hot state features with primitive actions (naive; c), SR features with primitive actions (SR; e), one-hot state features with primitive actions and eigenoptions (options; g; orange arrows, example option trajectories), HSR features with primitive actions and eigenoptions (HSR; i), and the factorised HSR, an overcomplete non-negative basis of the HSR (Methods), as state features with primitive actions and eigenoptions (fHSR; k). **d, f, h, j, l**. Learning curves (steps to goal per episode) of the agents in c, e, g, i and k in the small (left), medium (middle) and large (right) environments, for the 1st, 10th and 20th task of the series (tasks 1, 10 and 20). Dashed lines, optimal path length averaged over goals. **m**. Area under the learning curve relative to the optimal path length (AUC; lower is better; symmetric logarithmic axis) as a function of task for the five agents in each environment. All agents learn linear Q-functions ^24^ whose readout weights are reset at each change of goal. State representations are learned incrementally across tasks, as the running average of the representations under the policies learned for the completed tasks, starting from one-hot features in task 1 (Methods). Mean±s.e.m. over 10 random seeds for each of 9 goals.

The SR, however, scales poorly with environment size. Although its features summarise multi-step predictions, the agent still selects among primitive actions. The number of decisions separating start from goal therefore grows with the size of the environment, and over these decisions the goal must be found by exploration and its value propagated back to the start. Accordingly, in topologically identical environments of increasing size (Figure 1b), learning slowed with environment size at similar rates in naive and SR agents, although SR agents still improved across tasks (Figure 1d,f,m, middle and right panels).

To address this limitation, we turned to hierarchical control with options, which substantially shorten the temporal horizon of trial-and-error learning ^19^. Here we used eigenoptions, which are discovered from the slowest eigenmodes of the transition dynamics under a random-walk policy ^27^ and lead the agent towards topologically significant states such as doorways and corners (Figure 1g; Extended Data Figure 2; Methods). Augmenting naive agents with options (options agents; Extended Data Figure 1c) yielded the opposite profile to that of SR agents: they markedly accelerated learning in larger environments but did not transfer to new goals (Figure 1h,m). The predictive representation and options thus deliver either generalisation or scalability, but not both.

An immediate resolution is to combine the two. However, naively augmenting SR agents with options (SR + options; Extended Data Figure 1d; Methods) impaired both scalability and generalisation. Over successive tasks, learning became slower rather than faster (negative transfer), until it was no more efficient than in naive agents (Extended Data Figure 3a,b,e). In this situation, the SR is nevertheless computed at the leve of one-step transitions and ignores how options persist and terminate. It therefore misrepresents the dynamics induced by the hierarchical policy and hinders rather than helps learning. To resolve this mismatch, we proposed the HSR^23^, in which each option enters the construction of the map as a single compound transition linking its initial and terminal states, rather than as a sequence of one-step transitions through every intermediate state along the option’s trajectory (Equations 7; 8; Methods). The HSR therefore represents states consistently with the high-level policy and supports the similar linear decomposition of state values under that policy (Methods). Agents with HSR features and eigenoptions (HSR; Extended Data Figure 1e) achieved both scalable and generalisable learning, outperforming the naive combination of SR features with options (Figure 1i,j,m; cf. Extended Data Figure 3e). The benefit of the HSR thus comes from predicting the dynamics at the level at which behaviour is controlled, which naively adding options to the SR agent does not achieve. The advantage held across a range of option numbers and for an alternative definition of options (Extended Data Figure 4). It also persisted when the predictive representations (both SR and HSR) were learned online by temporal-difference learning (Extended Data Figure 5; Methods), and when the options themselves were discovered online from the agent’s own experience (Extended Data Figures 6a-e; 1h-j). When online option construction was applied on a linear track, HSR agents acquired ecologically plausible, ballistic movement after initial exploration, as observed in rodents ^28^ (Extended Data Figure 6f,g). Because the HSR is constructed from a task-agnostic set of options, it also changed substantially less than the standard SR between goals, and so depended less on the policy learned for any one goal (Extended Data Figures 3f-h and 6d,e).

The representations themselves show where this advantage comes from. Because the SR is locally diffusive, it assigns nearly identical predictive profiles to states within the same room and collapses each room onto nearly a single point. The HSR instead preserves distinct profiles within rooms alongside a clear between-room block structure (Extended Data Figure 3f,g,i). The same distinction held beyond space. In a non-spatial community-structure task ^7,29^, the HSR again captured the representational geometry measured in the human hippocampus more closely than the standard SR (Extended Data Figure 7).

Despite being more robust to policy changes than the SR, the HSR is nevertheless strictly policy-dependent. We therefore sought a single basis shared across tasks and proposed the fHSR, from which every task-dependent HSR can be linearly reconstructed with task-dependent loadings (Extended Data Figure 8a,b; Methods). For rewards drawn independently across states, finding the shared basis that minimises this reconstruction error is equivalent to finding the shared basis that minimises the error of linearly approximating the value functions of all tasks, again with task-dependent loadings (Supplementary Information, Section 3.3):

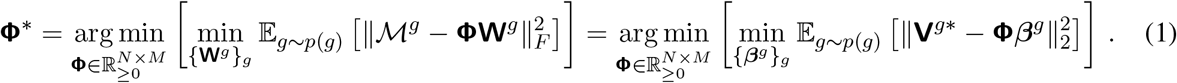

where **V**^*g*\*^ and ^*g*^ denote the vector containing the state values and the HSR matrix corresponding to the optimal policy for task *g*, respectively. The fHSR is therefore the optimal shared state representation for linear value approximation across tasks. Used directly as the agents’ state representation (Figure 1k; Extended Data Figure 1g), the fHSR matched HSR features in the smaller environments and improved transfer beyond them in the largest one (Figure 1l,m). By contrast, agents whose state representation was the same factorisation of the SR transferred poorly (fSR + options; Extended Data Figures 1f and 3c-e), so the factorisation helps only when both ingredients of the HSR, predictive representation and hierarchical control, are integrated appropriately. We constrained the fHSR to be non-negative and overcomplete, and these constraints cost little computationally: agents learned nearly as efficiently as with an unconstrained factorisation, particularly at higher degrees of overcompleteness (Extended Data Figure 8c-g). These additional constraints are, however, what allows the fHSR to model place cells tuning curves, as we show next.

### The fHSR reproduces the statistical characteristics of place-cell spatial tuning

We next asked whether the fHSR, the representation that provably supported optimally scalable and generalisable learning, could also account for the heterogeneous spatial tuning of place cells across environments of varying sizes. Specifically in larger environments, the distribution of field counts over all cells is overdispersed, and can be accurately described by a gamma-Poisson model ^11,15^: each cell has a Gamma-distributed ‘propensity’, defined as the expected number of fields per unit length, and its fields are drawn from a memoryless spatial Poisson process with the rate computed given the propensity. Reproducing the statistical consequences of this descriptive account from first principles is therefore a stringent test.

Classical predictive-map theories identify place cells with the columns of the predictive map ^3^, which comes with two caveats. First, the number of modelled neurons would then be capped at the number of states, which depends on the spatial resolution and the size of the environment. Second, both the SR and the HSR have a dominant diagonal, so every modelled neuron would express at least one field, contradicting the finding that only a minority of CA1 pyramidal neurons express place fields in any given environment ^11,30,31^.

We therefore modelled place cells as the fHSR, the task-agnostic basis obtained by an overcomplete, non-negative factorisation of the HSRs of all tasks (Equation 1; Methods). Each column of the fHSR models the spatial tuning of one model place cell. Non-negativity yields a parts-based, spatially localised code, and overcompleteness allows a single cell to carry several spatially separated fields (Supplementary Information, Section 3.5). The predictive map is then a property of the population, and individual cells are its reusable parts. On linear tracks, the tasks are the behaviours that a track affords, such as running rightward, running leftward and a random walk. As controls, we considered two alternative models: the factorised SR (fSR), obtained by applying the identical decomposition to the random-walk SR, and raw HSR columns. Place fields were identified with an identical, experimentally validated procedure in all simulated and real place cells (Extended Data Figure 9a; Methods). Given the normative nature of the theory, we sought qualitative and structural correspondence with the recordings rather than a statistical fit, and fixed one set of parameters for simulations across all tracks and both species (Methods).

The fHSR reproduced multi-field tuning with the correct population statistics. On the 48-m track, the number of fields per cell was heavy-tailed and favoured the gamma-Poisson over the equal-Poisson description by a wide margin, as in rat CA1 recordings on the same track ^11^ (Figure 2a,b). The same held in the much larger regime probed in bats: on a 200-m track, the fHSR again followed gamma-Poisson statistics that mirrored bat CA1 recordings ^12,14^ (Figure 2c,d). Neither control reproduced this structure (Figure 2b,d), indicating that place-cell-like tuning requires both the HSR and its decomposition (Supplementary Information, Sections 3 and 4).

**Figure 2:**
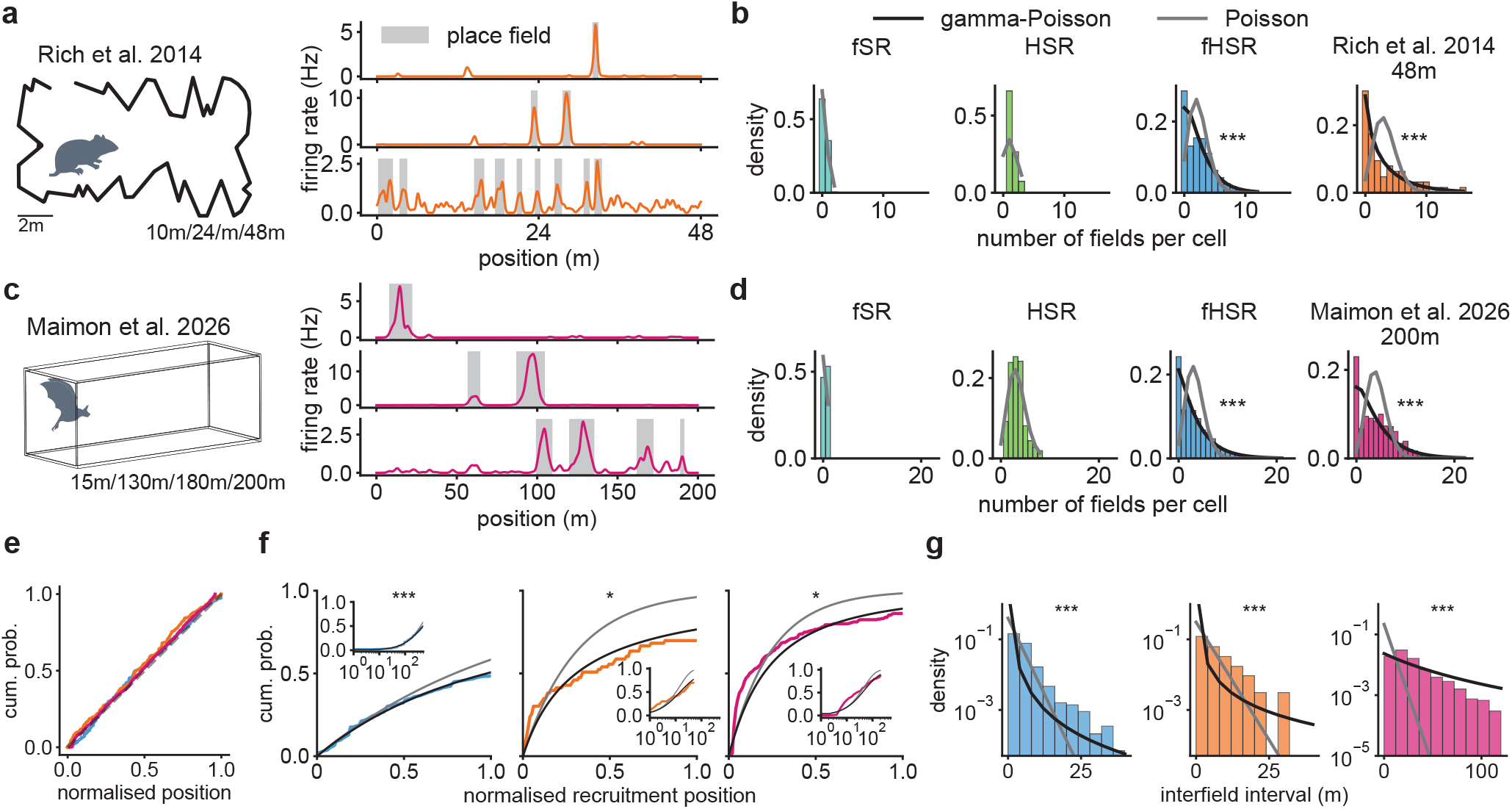
The factorised HSR (fHSR) reproduces the multi-field statistics of place cells in large environments. a. Left, rats run on linear tracks of 10, 24 and 48 m ^11^ (48-m track shown; scale bar, 2 m). Right, spatial tuning of three rat CA1 place cells on the 48-m track. Grey shading, identified fields. **b**. Distributions of the number of fields per cell on the 48-m track for (left to right) the factorised SR (fSR), raw HSR columns (HSR), the fHSR and rat CA1 place cells, with maximum-likelihood fits of the gamma-Poisson (black) and equal-Poisson (grey) descriptions ^11,15^ (likelihood ratio tests; fHSR: *χ*^2^(1) = 860.50, *p* = 3.8 10^*−*189^; rat: *χ*^2^(1) = 111.67, *p* = 4.2 × 10^*−*26^). **c**. Same as a, for bats flying along linear tracks of 15–200 m ^12,14^; tuning of three bat CA1 place cells on the 200-m track. **d**. Same as b, for the 200-m track (fHSR: *χ*^2^(1) = 3,597.35, *p <* 10^*−*308^; bat: *χ*^2^(1) = 733.22, *p* = 1.8 × 10^*−*161^). **e**. Cumulative distributions of normalised field locations for the fHSR (blue), rat (orange) and bat (magenta) place cells. Dashed line, uniform distribution. **f**. Cumulative distributions of the normalised recruitment position (location of each cell’s first field) for the fHSR (left), rat (middle) and bat (right) place cells, with the distributions implied by the gamma-Poisson (black) and equal-Poisson (grey) descriptions. Insets, same data on a logarithmic axis (fHSR: *χ*^2^(1) = 27.22, *p* = 1.8 × 10^*−*7^; rat: *χ*^2^(1) = 4.12, *p* = 0.042; bat: *χ*^2^(1) = 3.97, *p* = 0.046). **g**. Distributions of the intervals between successive fields within a cell, pooled over cells, with maximum-likelihood Lomax (gamma-Poisson; black) and exponential (equal-Poisson; grey) fits (fHSR: *χ*^2^(1) = 43,056.6, *p <* 10^*−*308^; rat: *χ*^2^(1) = 4,934.3, *p <* 10^*−*308^; bat: *χ*^2^(1) = 302.67, *p* = 8.7 × 10^*−*68^). Model results are pooled over 10 random seeds (*n* = 2000 cells); data, *n* = 253 rat cells from 5 rats and *n* = 291 bat cells from 8 bats. \**p <* 0.05; \*\**p <* 0.01; \*\*\**p <* 0.001.

Beyond field counts, the fHSR reproduced the finer spatial organisation of place fields. Field locations were distributed approximately uniformly along the track, as in rat and bat recordings, consistent with the homogeneous Poisson placement of fields (Figure 2e). The recruitment position, the location of each cell’s first field, followed the heavy-tailed distribution implied by the gamma-Poisson description, in both the model and real data (Figure 2f). Similarly, the intervals between successive fields, when pooled across cells, it followed a Lomax rather than an exponential distribution (Figure 2g), as expected if the intervals within each cell are exponential with a Gamma-distributed rate.

These structures originate in the range of scales spanned by the options. On a linear track, eigenoptions of increasing index partition the environment into progressively finer segments (Extended Data Figures 2c-e and 10a,b), so that a hierarchical policy selecting among them makes jumps of many lengths, distributed as a power law (Extended Data Figure 10c). The HSR therefore predicts occupancy far beyond the reach of standard SRs with the same discount, decaying logarithmically rather than exponentially with distance, and contains structure at every scale between the finest and the coarsest option (Extended Data Figure 10d,e; Supplementary Information, Section 2.3). Because its predictions remain coherent over such distances, a cell can split its support into several separated fields without losing predictive value. Doing so spreads its overlap with other cells thinly over many partners instead of concentrating it on a few, which the decomposition favours (Supplementary Information, Section 3.5). Each cell’s number of fields therefore grows with the range of scales it spans, and because this range varies multiplicatively across cells, field propensity becomes approximately Gamma-distributed (Supplementary Information, Section 4). Accordingly, replacing the eigenoptions with options of a single scale markedly reduced both the number of fields per cell and their overdispersion (Extended Data Figure 10f,g). The place-cell statistics is thus contingent on acting with options at multiple scales; temporal abstraction at a single scale does not suffice.

These statistics were robust to the optimisation procedure used to extract the basis (Extended Data Figure 11), to the parameters of the HSR and its decomposition (Extended Data Figure 9b-f), and to the criteria used to identify firing fields, which we applied identically to modelled and real place cells (Extended Data Figure 9g-k).

### Within-cell multi-scale fields support accurate and scalable spatial decoding

The multi-scale origin of these statistics makes a further prediction for individual cells: a cell that spans a wide range of scales should carry fields of markedly different sizes rather than repeats of one characteristic width (Supplementary Information, Section 3.5) ^12,14^. Among fHSR cells with at least two fields, the largest field was typically several times wider than the smallest, and this ratio spanned more than an order of magnitude across cells (Figure 3a). Bat ^14^ and rat ^11^ CA1 place cells showed the same within-cell multi-scale tuning (Figure 3b,c).

**Figure 3:**
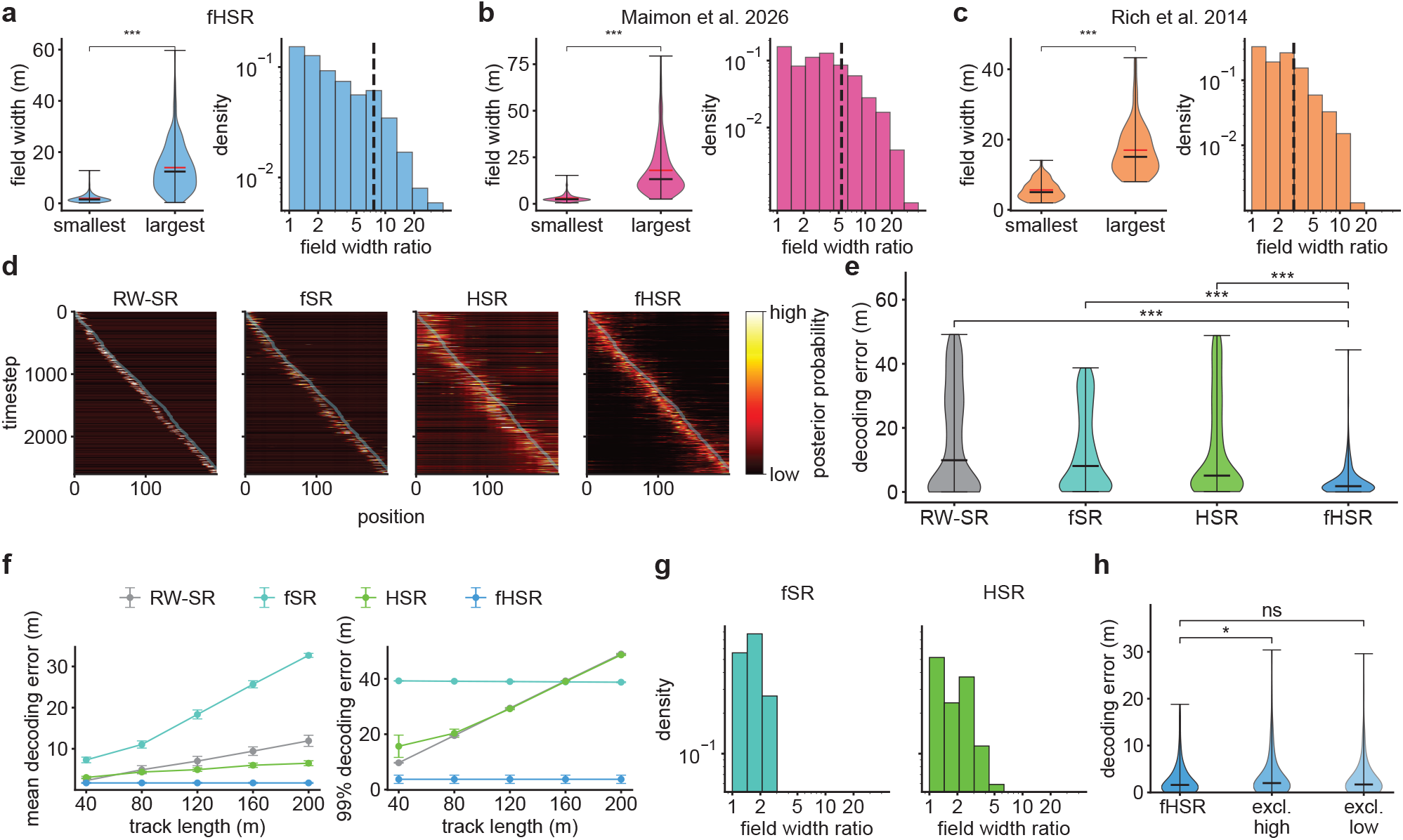
Within-cell multi-scale tuning in the factorised HSR (fHSR) supports accurate and scalable spatial decoding. a. Left, widths of the smallest and largest field of each fHSR cell with at least two fields on the 200-m track (*n* = 1,235 cells; black and red horizontal lines, median and mean). Right, distribution of the within-cell field-width ratio (largest/smallest); dashed line, median ratio. **b**. Same as a, for bat CA1 place cells ^14^ (*n* = 433 cells). **c**. Same as a, for rat CA1 place cells ^11^ (*n* = 253 cells). **d**. Posterior probability over position decoded from simulated population activity along a rightward trajectory on the 200-m track (true position, cyan) for RW-SR columns ^3^, the factorised SR (fSR), raw HSR columns (HSR) and the fHSR. **e**. Distributions of per-step decoding error for the four representations in d (two-sided Mann–Whitney *U* tests, *n* = 651 decoding steps per representation; RW-SR columns versus fHSR: *U* = 290,520.5, *p* = 4.5 × 10^*−*31^; fSR versus fHSR: *U* = 306,710.0, *p* = 2.1 × 10^*−*44^; HSR columns versus fHSR: *U* = 326,934.0, *p* = 1.6 × 10^*−*64^). **f**. Mean (left) and 99th-percentile (catastrophic; right) decoding error as a function of track length (mean ± s.e.m. over 10 random seeds per length). **g**. Distributions of the within-cell field-width ratio for the fSR (left) and raw HSR columns (right). RW-SR columns have exactly one field per cell and therefore no within-cell ratio. **h**. Per-step decoding error for a random 90% subsample of fHSR cells (fHSR) and after excluding the decile of cells with the highest (excl. high) or lowest (excl. low) within-cell field-width ratios (two-sided Mann–Whitney *U* tests, *n* = 651 steps; subsample versus excl. high: *U* = 195,089.5, *p* = 0.013; subsample versus excl. low: *U* = 207,167.5, *p* = 0.49). Violin plots: horizontal lines, median. \**p <* 0.05; \*\*\**p <* 0.001; ns, not significant.

A representation that supports learning in large environments should also remain informative about self-position as the environment grows. We tested this by Bayesian decoding of position from simulated population activity along a 200-m track (Methods), comparing random-walk SR (RW-SR) columns ^3^, the fSR, raw HSR columns and the fHSR (Figure 3d). The fHSR yielded markedly more accurate spatial decoding than all three controls (Figure 3e), and its advantage grew with environment size: across tracks of 40-200 m, both its mean and its catastrophic (99th-percentile) decoding errors remained small, whereas the mean errors of all three controls grew with track length and their catastrophic errors reached tens of metres (Figure 3f). A population code with multi-scale fields escapes the spatial aliasing induced by the single-scale trade-off^17^: narrow fields set local precision, while broad fields rule out distant candidate locations (Supplementary Information, Section 5). Consistent with the decoding results, the field-width ratios of the fSR and of raw HSR columns were concentrated near one (Figure 3g), and RW-SR columns carry a single field per cell. Selectively removing cells from the fHSR further revealed an asymmetric contribution of the two ends of the scale range: excluding the decile of cells with the highest within-cell field-width ratios impaired decoding relative to a random 90% subsample, whereas excluding the decile with the lowest ratios did not (Figure 3h). Cells that span the widest range of scales thus contribute disproportionately to accurate decoding.

### The fHSR predicts how field statistics scale with environment size

The comparisons so far were made at single track lengths, but real place cells exhibit markedly different firing profiles across environment sizes (Figure 4a). In the fHSR, such differences are expected from the way the option hierarchy grows with the environment. The finest option scale is set by the correlation length of the RW-SR and does not depend on track length, whereas the coarsest option spans the whole track. The range of option scales therefore grows in proportion to track length (Methods; Supplementary Information, Section 7.1). Because each cell’s number of fields grows with the range of scales it spans (see above), field statistics should change systematically, and approximately as power laws, with track length (Supplementary Information, Section 4). We tested this prediction on tracks matching each length in the rat (10, 24 and 48 m) and bat (15, 130, 180 and 200 m) recordings. The spatial resolution and all model parameters were fixed, so that only the number of states, and with it the number of options, changed (Methods). In both the modelled and real place cells, field counts favoured the gamma-Poisson over the equal-Poisson description at every length (Figure 4b), except for bat place cells on the 15-m track, possibly because few neurons were recorded there.

**Figure 4:**
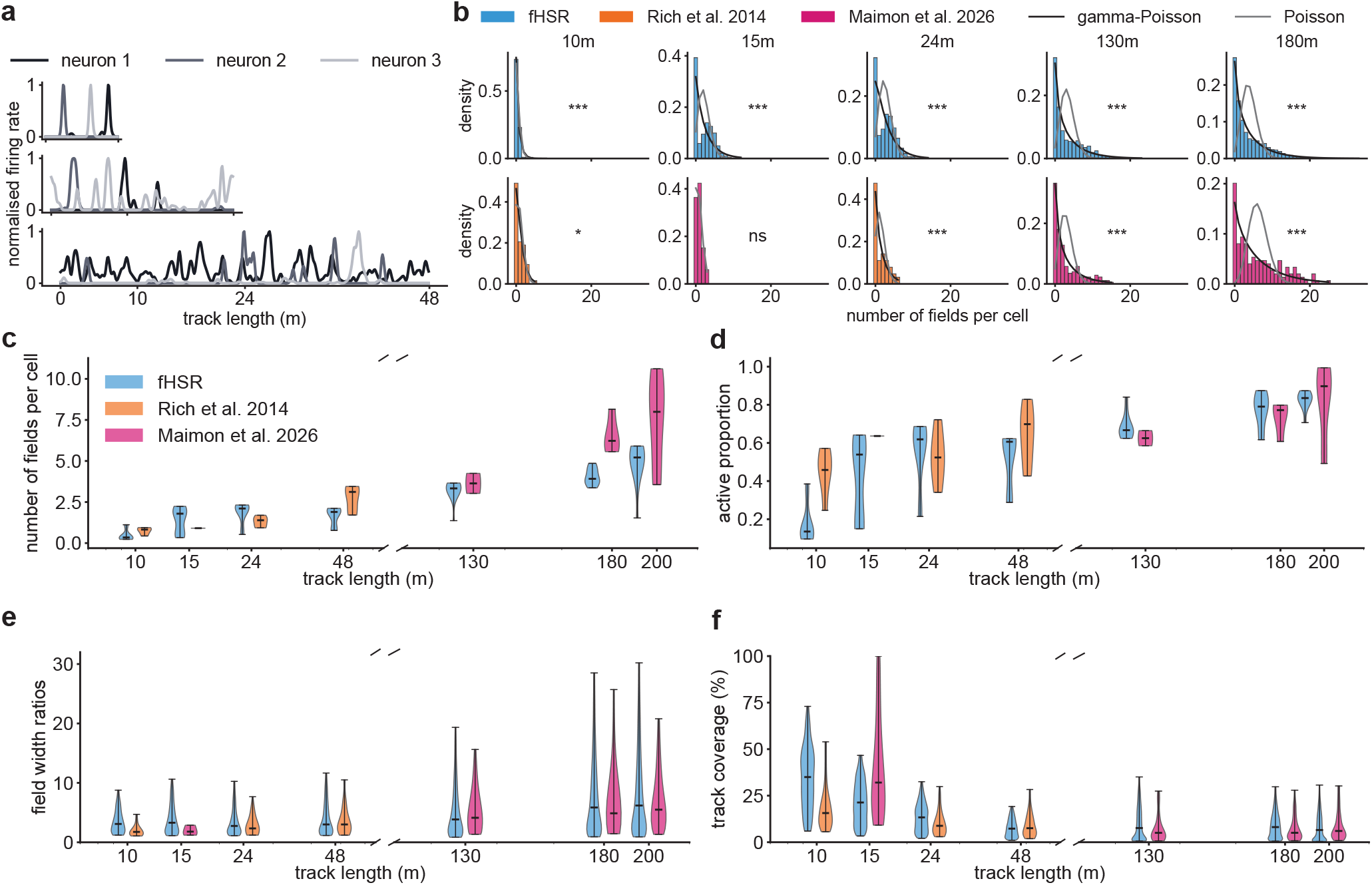
The factorised HSR (fHSR) reproduces how place-cell statistics scale with environment size. a. Normalised spatial tuning of three rat CA1 place cells recorded on the 10-m (top), 24-m (middle) and 48-m (bottom) tracks ^11^. **b**. Distributions of the number of fields per cell for the fHSR (top) and recorded place cells (bottom; rat, orange; bat, magenta) on tracks of 10, 15, 24, 130 and 180 m, with maximum-likelihood fits of the gamma-Poisson (black) and equal-Poisson (grey) descriptions (likelihood ratio tests; fHSR 10 m: *χ*^2^(1) = 164.54, *p* = 1.2 10^*−*37^; fHSR 15 m: *χ*^2^(1) = 1,227.03, *p* = 8.2 × 10^*−*269^; fHSR 24 m: *χ*^2^(1) = 1,173.73, *p* = 3.1 × 10^*−*257^; fHSR 130 m: *χ*^2^(1) = 5,138.18, *p <* 10^*−*308^; fHSR 180 m: *χ*^2^(1) = 7,511.12, *p <* 10^*−*308^; rat 10 m: *χ*^2^(1) = 4.27, *p* = 0.039; bat 15 m: *χ*^2^(1) = 0.0, *p* = 1.0; rat 24 m: *χ*^2^(1) = 27.88, *p* = 1.3 × 10^*−*7^; bat 130 m: *χ*^2^(1) = 341.90, *p* = 2.5 × 10^*−*76^; bat 180 m: *χ*^2^(1) = 491.90, *p* = 5.5 × 10^*−*109^). Results for 48 and 200 m are shown in Figure 2b,d. **c-f**. Mean number of fields per cell (c), proportion of active cells (cells with at least one field; d), within-cell field-width ratio (e) and per-cell track coverage (fraction of the track covered by a cell’s fields; f) as a function of track length, for the fHSR (blue), rat (orange) and bat (magenta) place cells. Horizontal ticks, medians. All model, decomposition and field-detection parameters, and the spatial resolution, are identical at every length (Methods). \**p <* 0.05; \*\*\**p <* 0.001; ns, not significant.

Model and data shared four scaling trends. The mean number of fields per cell, the proportion of active cells and the within-cell field-width ratio increased with track length, whereas the fraction of the track covered by each cell’s fields decreased (Figure 4c-f). Despite differences in species, behaviour and recording methods, the rat and bat data followed approximately continuous trends across the full range of lengths, and the model captured the trends of both species. As predicted, these statistics scaled approximately as power laws of track length in the model, with exponents that were robust to variations in model, decomposition and detection parameters (Extended Data Figure 12).

The random-process account of place fields ^16^ deals with environment size differently, which reproduced the sizes, spacing and shapes of fields by modelling each cell’s tuning curve as a random function, drawn independently from a stationary Gaussian process and rectified at a threshold. The keys parameters of the model were fitted separately to each environment, and were shown to exhibit expected scaling with environment size. However, it does not explain why its parameters, and the associated field statistics, should change with size. In contrast, this change follows from the expansion of the option hierarchy with environment size under the fHSR framework (Supplementary Information, Section 4 and 7.1).

### Population activity of the fHSR shows hyperbolic-like topology

Moving from single cells to the population, we compared the organisation of co-activity in the model and the data using the clique-topology analysis with which the hyperbolic signature of rat CA1 activity on the 48-m track was identified ^17^. Clique topology summarises a matrix of pairwise correlations between neuronal activities (Figure 5a) by the number of *n*-dimensional holes (Betti numbers *β*_*n*_) in the sequence of undirected graphs obtained by progressively connecting the most strongly correlated pairs of cells (Figure 5b). Because it depends only on the rank order of correlations, it reveals the intrinsic structure of co-activity independently of how correlations relate to distances ^32^. On the same track, the fHSR yielded Betti curves that closely resembled those of the rat recordings and of a three-dimensional hyperbolic reference model (Figure 5c), and the integrated Betti numbers of the data fell within the range spanned by model realisations (Figure 5d). The hyperbolic-like topology arises because each fHSR cell is characterised by both a position and a spatial scale (Supplementary Information, Section 6). Such a code resembles a tree, in which each coarse field is subdivided by finer ones, so that the number of distinguishable positions doubles with every halving of scale. This exponential branching is the defining feature of hyperbolic space ^33^. None of the control representations showed this topology: RW-SR columns, the fSR and raw HSR columns produced Betti numbers one to two orders of magnitude larger than those of the data (Extended Data Figure 13a). Our analysis is based on correlations between spatial tuning maps, and closely tracked correlations computed from spike trains ^17^ in rat recordings: both yielded indistinguishable Betti curves (Extended Data Figure 13b-g).

**Figure 5:**
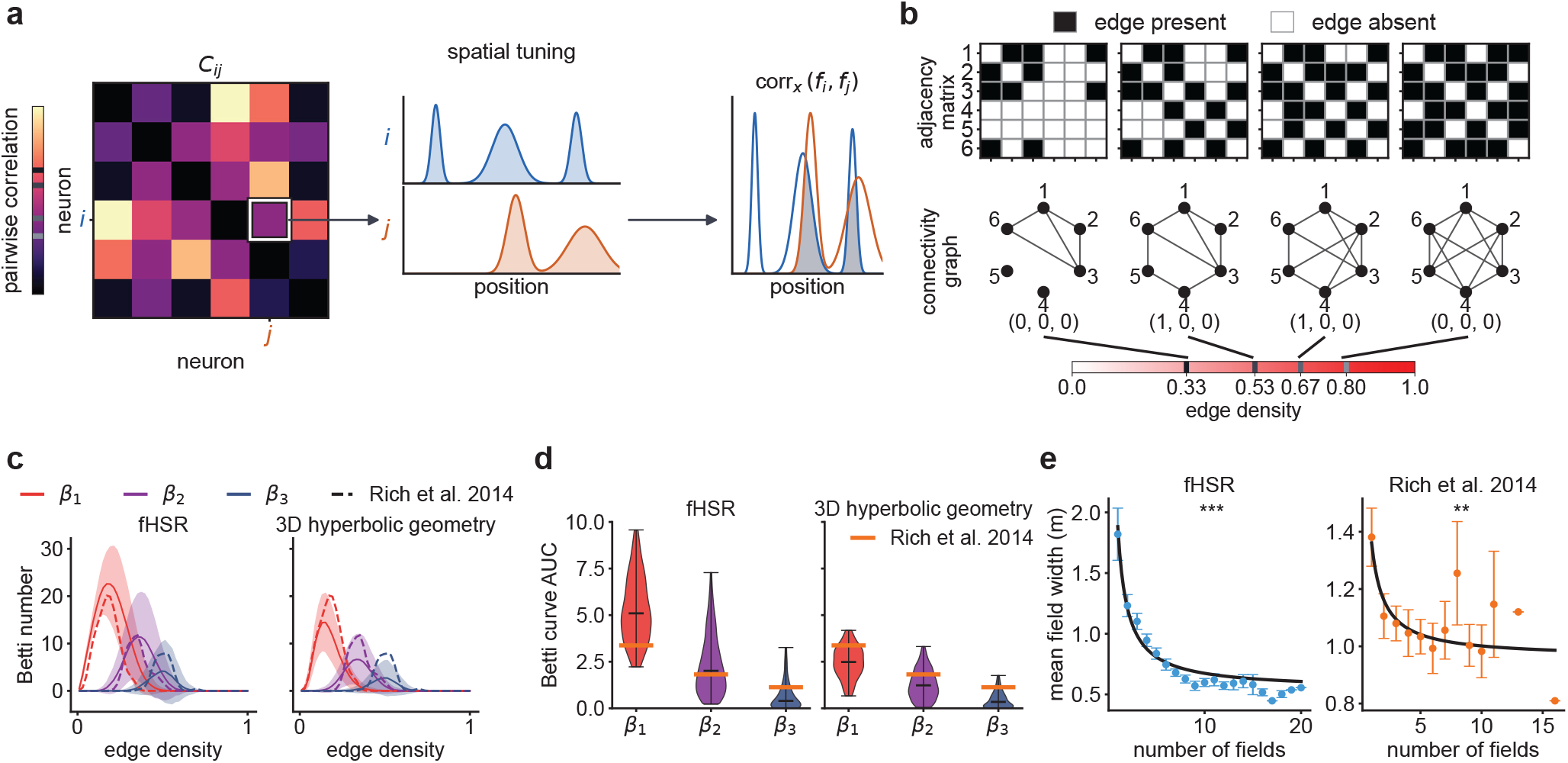
The factorised HSR (fHSR) reproduces the hyperbolic-like topology of hippocampal population activity. **a**. Construction of the correlation matrix *C*_*ij*_: for each pair of cells *i* and *j*, the correlation between their spatial tuning curves *f*_*i*_ and *f*_*j*_. **b**. Clique topology ^32^. Edges are added between pairs of cells in order of decreasing correlation; at each edge density, the adjacency matrix (top) defines a graph (bottom) whose one-, two- and three-dimensional cycles are counted by the Betti numbers (*β*_1_, *β*_2_, *β*_3_) (in brackets). Betti numbers as a function of edge density define Betti curves. Example with six cells. **c**. Betti curves of the fHSR (left) and of a three-dimensional hyperbolic reference model ^17^ (right), mean±s.d. over 300 random seeds. Dashed lines, rat CA1 place cells on the 48-m track ^11^. **d**. Integrated Betti curves (area under the curve) across random seeds for the fHSR (left) and the hyperbolic reference (right). Orange lines, rat CA1 data. **e**. Mean field width as a function of the number of fields per cell for the fHSR (left) and rat CA1 place cells (right) (mean±s.e.m. across cells with a given number of fields). Solid lines, least-squares fits of *y* = *b* + *a/x* (fHSR: *a* = 2.70, *b* = 0.82; rat: *a* = 0.40, *b* = 0.96; two-sided Wald tests on *a*, fHSR: *p* = 3.5 × 10^*−*26^; rat: *p* = 0.0022).

We next asked whether irregular multi-field tuning alone, modelled by independent random Gaussian process functions ^16^, could produce this topology. Relocating each cell’s fields at random, preserving their number and widths, inflated all three Betti curves of the rat data (Extended Data Figure 13h,i). Nor did the topology emerge from the random-function model (see above), simulated with the parameters fitted to the rat data on the 48-m track (Methods). Random-function populations matched the data in one-dimensional cycles. However, in their canonical, noiseless form they lacked the three-dimensional cycles of the data, and with tuning curves re-estimated from Poisson spike counts they produced an excess of two- and three-dimensional cycles, whether the threshold was fixed or varied across cells (Extended Data Figure 13j,k). The hyperbolic-like topology therefore reflects how fields are arranged across cells, rather than the irregularity of individual cells.

The hyperbolic-geometry account, however, is agnostic to how fields are arranged within cells. It models each neuron as a point in a hyperbolic space, and where it considers multiple fields per cell, their number is drawn from the empirical gamma-Poisson statistics and their sizes are drawn independently of that number ^17^. The fHSR, in contrast, predicts how fields are arranged within cells. Because the decomposition creates multiple fields by splitting a cell’s support rather than by adding to it (Supplementary Information, Section 3.5), a cell’s total field extent should be approximately conserved, so cells with more fields should have narrower fields. Both modelled and rat place cells showed this reciprocal relationship (Figure 5e), which the hyperbolic geometry alone does not determine.

### Population activity shows a predicted spatial periodicity independent of environment size

At the population level, the theory makes a previously untested prediction. On a linear track, each eigenoption terminates on a regular lattice of locations (Extended Data Figures 2d and 10b), so cells that draw on the same option scale inherit a common spatial phase. Although each cell’s fields are irregularly placed, the summed activity of the population should therefore carry a weak spatially periodic modulation, with a wavelength set by the finest option scale (Supplementary Information, Sections 7.2-7.4). Because this scale does not depend on track length (see above), neither should the wavelength.

Summed activity of the fHSR was indeed periodically modulated along 10-, 24- and 48-m tracks (Figure 6a). Its spatial autocorrelation peaked at similar non-zero lags across track lengths, with no detectable trend in this characteristic wavelength (Figure 6b,c). The centroid of its power spectral density was likewise similar across lengths (Figure 6g,i). Both statistics differed significantly from those of shuffled populations in which each cell’s fields were relocated at random, preserving their number and widths (Figure 6b,g). The periodicity is compatible with the near-uniform distribution of field locations (Figure 2e) because it is carried by a weak modulation of activity across cells, anchored to option terminal states (Extended Data Figure 14), rather than by a clustering of fields at particular locations. The descriptive account did not reproduce this modulation. Populations whose fields were placed independently, with counts drawn from either equal-Poisson or gamma-Poisson processes matched to the model, lacked it (Figure 6j; Supplementary Information, Section 7.4).

**Figure 6:**
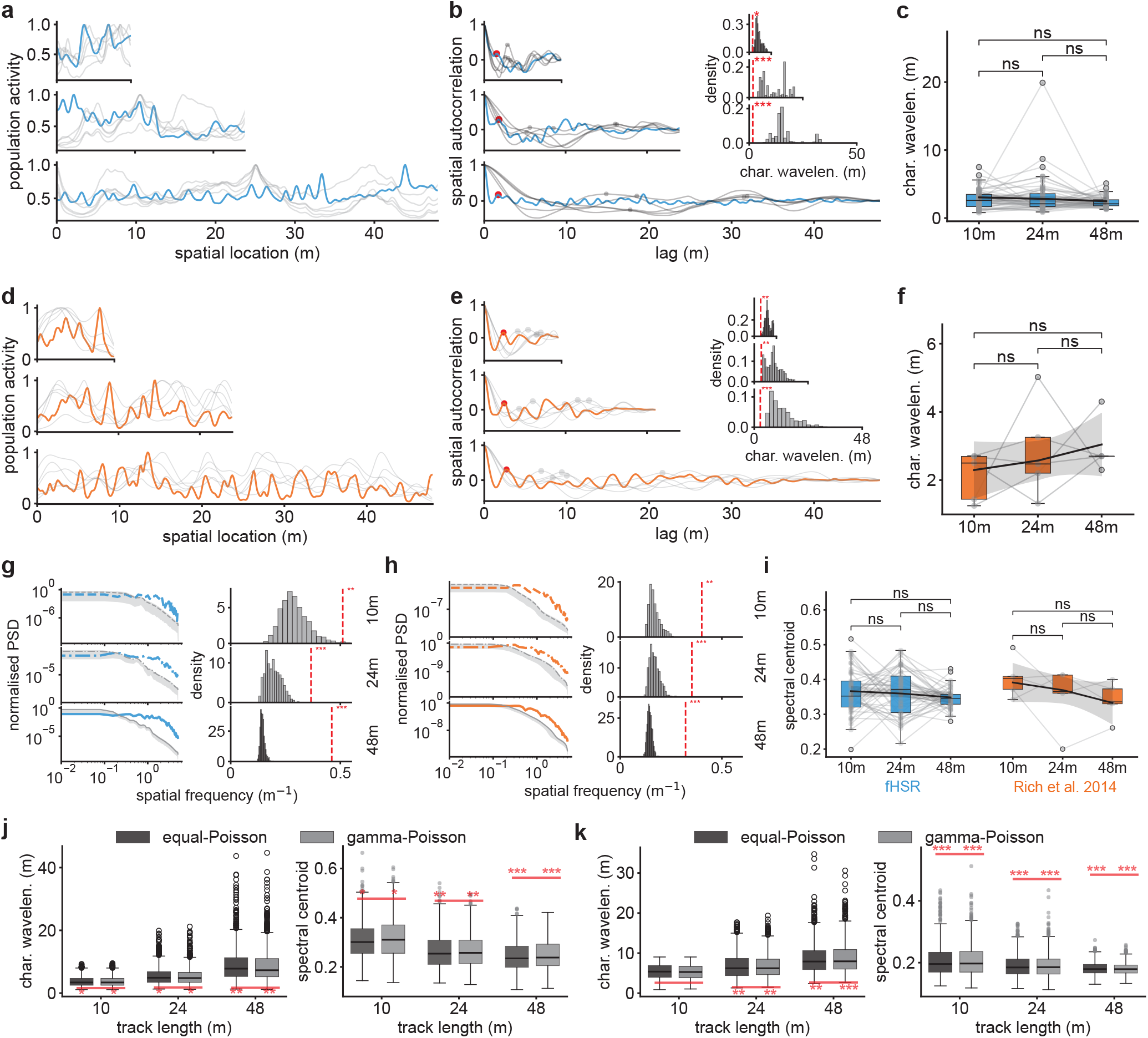
The factorised HSR (fHSR) predicts a spatially periodic modulation of population activity whose wavelength is independent of environment size, as observed in rat CA1. a. Summed population activity of the fHSR along 10-, 24- and 48-m tracks (blue). Grey, example populations in which each cell’s fields are relocated at random, preserving their number and widths (shuffles). **b**. Spatial autocorrelations of the profiles in a. Red dots, characteristic wavelength (lag of the first non-zero peak); grey, shuffles. Insets, null distributions of the characteristic wavelength over 1,000 shuffles; red dashed lines, model (10 m: *p* = 0.018; 24 m and 48 m: *p <* 0.001). **c**. Characteristic wavelengths over different track lengths across random seeds (*n* = 100; grey lines connect seeds). Two-sided Wilcoxon signed-rank tests: 10 versus 24 m, *W* = 468.5, *p* = 0.76; 10 versus 48 m, *W* = 331.5, *p* = 0.087; 24 versus 48 m, *W* = 386.0, *p* = 0.17. Black line, least-squares linear fit (slope *a* = −0.015, intercept *b* = 3.20 m; two-sided Wald test on *a, p* = 0.18). Linear mixed-effects model (LMM) of the characteristic wavelength on track length (Methods), *β* = −0.0151, *p* = 0.1791. **d-f**. Same as a-c, for rat CA1 place cells ^11^ (shuffle tests, 10 and 24 m: *p* = 0.003; 48 m: *p <* 0.001; Wilcoxon signed-rank tests across *n* = 5 rats, 10 versus 24 m: *W* = 5.0, *p* = 0.63; 10 versus 48 m: *W* = 2.0, *p* = 0.19; 24 versus 48 m: *W* = 7.0, *p* = 1.0; linear fit, *a* = 0.011, *b* = 2.10 m, Wald test on *a, p* = 0.25; shading, 95% confidence interval; LMM, *β* = 0.0197, *p* = 0.2508). **g**. Left, normalised spatial power spectral density (PSD) of the summed activity of the fHSR on each track (blue) and of shuffles (grey; mean ± s.d. over 1,000 shuffles). Right, null distributions of the spectral centroid over shuffles; red dashed lines, model (10 m: *p* = 0.007; 24 and 48 m: *p <* 0.001). **h**. Same as g, for rat CA1 place cells (10 m: *p <* 0.001; 24 m: *p* = 0.001; 48 m: *p <* 0.001). **i**. Spectral centroids across random seeds (fHSR, *n* = 45) and rats (*n* = 5). Two-sided Wilcoxon signed-rank tests, fHSR, 10 versus 24 m: *W* = 509.0, *p* = 0.93; 24 versus 48 m: *W* = 372.0, *p* = 0.10; 10 versus 48 m: *W* = 397.0, *p* = 0.18; rats, 10 versus 24 m: *W* = 4.0, *p* = 0.44; 24 versus 48 m: *W* = 6.0, *p* = 0.81; 10 versus 48 m: *W* = 1.0, *p* = 0.13. Black lines, least-squares linear fits. **j**. Characteristic wavelength (left) and spectral centroid (right) of populations in which fields are placed independently, with field counts drawn from equal-Poisson (dark grey) and gamma-Poisson (light grey) processes matched to the fHSR (1,000 resamples); red lines, values of the fHSR. Empirical *p* values, 10/24/48 m: characteristic wavelength, equal-Poisson, *p* = 0.027*/*0.018*/*0.001; gamma-Poisson, *p* = 0.022*/*0.013*/*0.004; spectral centroid, equal-Poisson, *p* = 0.021*/*0.006*/<*0.001; gamma-Poisson, *p* = 0.034*/*0.003*/<*0.001. **k**. Same as j, for rat CA1 place cells (1,000 resamples; characteristic wavelength, equal-Poisson, *p* = 0.075*/*0.001*/*0.002; gamma-Poisson, *p* = 0.069*/*0.001*/<*0.001; spectral centroid, equal-Poisson and gamma-Poisson, *p <* 0.001 on all tracks). Box plots: centre line, median; box, interquartile range; whiskers, 1.5× interquartile range; points, outliers. \**p <* 0.05; \*\**p <* 0.01; \*\*\**p <* 0.001; ns, not significant.

We found qualitatively similar periodicity in rat CA1 place cells, again with no detectable trend in wavelength with track length (Figure 6d-f,h,i). As in the model, the periodicity departed from that of independently placed equal- and gamma-Poisson populations (Figure 6k). It did not arise from the many turns of the winding tracks (Extended Data Figure 15a-c). Population activity was not locked to turns (Extended Data Figure 15d,e), and its phase at turns was uniformly distributed (Extended Data Figure 15f). The same pattern held in bats. Summed activity of bat CA1 place cells ^14^ on 130-200 m tracks was also periodically modulated, with a wavelength an order of magnitude longer that did not differ detectably between track lengths (Extended Data Figure 15g-j). Summed activity given tuning curves modelled by random functions ^16^, by contrast, did not exhibit any significant periodicity comparing to that of shuffled populations (Extended Data Figure 15k), so the periodicity found in both species is not a generic consequence of irregular multi-field tuning at the single-cell level.

## Discussion

Our theoretical framework traces the irregular, multi-field and multi-scale place cells in naturalistically large environments to a principled computational account: learning and planning that both scale with environment size and generalise across tasks. Among the representations we tested, only a predictive map computed under hierarchical control met both demands, and the options that make learning scalable also shape the neural code. The range of scales they span sets how many fields each cell expresses and how that number changes with the environment, gives the population its hyperbolic-like geometry and, through its finest scale, fixes the wavelength of the population periodicity. What makes a representation useful for learning thus also determines the statistics of the cells that carry it. On this view, the multi-field place cells of large environments are the form that an efficient predictive map takes when behaviour is organised hierarchically, rather than a by-product of biological implementation.

Previous predictive-map accounts proposed multiple spatial scales from SRs arising from varying planning horizon (discount factor) ^3,34,35^, possibly ordered along the dorsoventral hippocampal axis ^36^. Regardless of the scale, SRs still predict step-by-step behaviour, so they do not shorten the sequences of decisions over which learning and planning operate, and they yield cells with single fields whose size differs between cells, not cells with several fields of different sizes. In the HSR, multiple scales arise from a single discount through the nested durations of options, and its factorisation distributes them within cells, providing a normative origin for the gamma-Poisson descriptive account of place-field distributions ^11,15^.

The random-function model offers a mechanistic account of the same statistics ^16^. It assumes that random projections from afferent neurons make each place cell’s tuning curve resemble a random function drawn from a Gaussian process, with place fields defined as the excursions of this function above a pre-specified threshold. We derived the field statistics of the fHSR with a similar threshold-crossing analysis, but the spatial correlations of its tuning curves are set by the option hierarchy rather than fitted to the experimental data (Supplementary Information, Section 4). Irregular field arrangements, compatible with predominantly random synaptic input to CA1^16^, can equally arise in an optimised code, because the decomposition constrains field positions only through the overlap between cells (Supplementary Information, Section 3.7). The two accounts diverge at the population level: given the independent sampling of each cell’s tuning maps, the random-function model lacked both the hyperbolic-like topology and the spatial periodicity of CA1 population activity (Extended Data Figures 13j,k and 15k), whereas the fHSR derives both from the option hierarchy shared by all cells.

The components of the HSR formulation maps onto an entorhinal-prefrontal-hippocampal circuit. Eigenoptions derive from the eigenvectors of the random-walk SR. With a uniform default policy and step costs, the random-walk SR is equivalent, up to a scale factor, to the goal-independent default representation in linear reinforcement learning, with its eigenvectors resemble entorhinal grid fields ^3,37^. Grid modules, with their discrete progression of scales ^38^, are thus a natural source of options at nested scales. Executing options requires tracking progress towards subgoals, a signal that could be carried by medial frontal neurons in mice ^39^ and by prefrontal and cingulate cortex in humans ^20,21,40^. The hippocampus would then learn the HSR under the resulting hierarchical policy, for which neural implementations of such online learning at the synaptic level have been proposed for the standard SR^41,42^. Hippocampal sequences, which can traverse long distances in single jumps ^43,44^ with superdiffusive statistics ^45,46^, would in turn sample a map that itself predicts long, heavy-tailed jumps rather than local diffusion (Supplementary Information, Section 2.3), as planning over subgoals requires ^47^.

The periodicity in population activity is a direct test of the theory. It inherits its wavelength and phase from the eigenoptions (Supplementary Information, Sections 7.1 and 7.2), and options without a shared lattice of terminal states would not produce it. Its presence in rats and bats is therefore consistent with options derived from shared eigenmodes, and its wavelength may read out the finest scale at which the animal segments its behaviour. Disrupting grid-cell input should erode the within-cell multi-scale structure of place cells, whereas inactivating prefrontal cortex, which removes options from behaviour and hence from the map, should shift place cells towards the single-field code of the fSR (Figure 2b,d), abolish the periodicity and make learning slow down more steeply with environment size (Figure 1m).

The fHSR is a sparse, overcomplete, non-negative code of the kind proposed for sensory cortex ^48^, but optimised to reconstruct predictive maps, and hence values, across tasks rather than sensory input (Supplementary Information, Section 3.3). This code also admits a neurally plausible implementation. Because the fHSR depends on the HSR only through the similarity between the futures predicted from different states, and because refining its initial factorisation barely changes it (Extended Data Figure 11), the basis is, to a close approximation, a symmetric non-negative factorisation of that predictive similarity (Methods). A single-layer recurrent network with Hebbian feedforward and anti-Hebbian lateral plasticity could perform this computation online ^49^, and under specific instantiations, its output neurons develop localised, place-field-like receptive fields when its inputs vary smoothly with position ^50^. Here we computed the HSR and its decomposition offline, for familiar environments and a fixed option set. Future work could extend to learning both online, with the HSR updated by temporal-difference learning while a Hebbian/anti-Hebbian network continuously re-factorises it into place cells. Further combined with online option construction (Extended Data Figure 6), such a model would predict how place fields form during exploration and how they reorganise when the environment or the option set changes.

The current study has several limitations that point to future work. The quantitative deviations between model and data (Figures 4c,d and 5e) suggest that factors outside the normative theory, such as heterogeneous excitability ^15^, also shape field statistics. Modelling place cells as a single task-agnostic basis gives each model cell identical fields in the two running directions, at odds with the directional firing observed on linear tracks ^51,52^. The theory could accommodate this with small task-specific populations, each obtained by applying the same decomposition to the HSR of one behaviourally salient task, such as leftward or rightward running. Each population would be active only during its own task, and hence in one running direction, with the task-agnostic basis serving all other tasks. Our comparisons with real data were restricted to linear tracks, whereas natural navigation unfolds in large two-dimensional spaces, where place cells likewise express multiple fields ^10,13^. In such spaces, the theory predicts elevated population activity at bottlenecks and other topologically significant locations where options terminate (Extended Data Figure 14). This is consistent with the accumulation of fields near goals and other salient locations ^53,54,55^, if these also serve as option terminals. More generally, place-cell statistics should change when the hierarchical organisation of behaviour changes, even if the geometry of the environment does not: intermediate rewards or landmarks should create new option terminals and redistribute fields, and learning a shortcut should reshape the predictive geometry that the fHSR encodes. Recordings in large, structured two-dimensional environments could thus reveal how an animal decomposes its world into subgoals.

## Methods

Notation follows the Supplementary Information: bold symbols denote vectors and matrices, and calligraphic symbols denote the HSR and its operators. *N* is the number of states, *M* the number of model cells and *K* the number of options.

### Environments and tasks

Navigation experiments were based on deterministic, discrete Markov decision processes ^25^, in which movements into a boundary leave the agent in place. Primitive actions are the four cardinal moves in two-dimensional environments and left and right moves on linear tracks. The four-room environment consists of four square rooms connected pairwise by single-state doorways. Small, medium and large versions (grids of side 11, 17 and 25 states) were obtained by enlarging each room while preserving the room-and-doorway topology and the room-relative positions of the start and goal states (Figure 1b). Entering the goal of the current task yields a reward of 1 and immediately terminates the episode, and all other transitions yield 0. Goal locations are selected such that no goal coincides with the start state or with an option terminal state, and are maximally uniformly distributed within the four rooms. Each task lasts 50 episodes of at most 1,000 steps, all starting from the start state. Optimal path lengths were computed by dynamic programming.

In the main navigation experiments (Figure 1; Extended Data Figures 3, 4 and 8), agents faced a series of tasks that share the start state and differ in their goal, drawn from 20 goal locations placed at the same room-relative positions in every environment size (Figure 1a). We report performance on the 1st, 10th and 20th task of the series. Each of 9 goals (goals 1-9 in Figure 1a) served in turn as the goal of the reported task, with the preceding tasks drawn from the remaining goals, and each condition was repeated with 10 random seeds. A simpler transfer setting (Extended Data Figures 5 and 6a-e) used three goals, one in each room other than that of the start state (Extended Data Figure 5a), one per task, presented in all six orderings with 10 random seeds each.

Place-cell analyses used linear tracks discretised at a fixed resolution of Δ*x* = 0.1 m per state, with lengths matching the recordings: 10, 24 and 48 m for rats ^11^ and 15, 130, 180 and 200 m for bats ^12,14^, with additional tracks of 40–200 m for decoding. Longer tracks therefore have proportionally more states. Tracks of 100 states were used to illustrate the decomposition (Extended Data Figure 8b) and online option construction (Extended Data Figure 6f,g), in which agents ran alternately to the two ends of the track. The analyses of option terminal states (Extended Data Figure 14) used an open arena, the four-room environment, a two-room environment with an off-centre door and a four-room environment with shifted doors.

### Reinforcement-learning agents

All agents learned action values by Q-learning with linear function approximation ^24^, 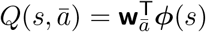, where *ā* is a primitive action or an option and ***ϕ***(*s*) is the feature vector of state *s* (Extended Data Figure 1). The features are a one-hot vector, the row of the SR or of the HSR corresponding to *s*, or the corresponding row of the factorised SR or HSR (fSR or fHSR) – the non-negative bases obtained by decomposing the SR or the HSR (see below). The action set comprises either the primitive actions alone or the primitive actions augmented with the option set. Their combinations define seven agents (Extended Data Figure 1a-g). Naive (one-hot features) and SR agents use primitive actions only. Options (one-hot features), SR + options and fSR + options agents add the option set, as do HSR and fHSR agents, whose representations are defined with respect to it. SR agents use the SR of their own policy. For agents that combine the SR with options (SR + options and fSR + options), the SR was computed from the one-step transition matrix induced by the hierarchical policy *µ*, **T**_*µ*_(*s, s*^′^) = _*ā*_ *µ*(*ā s*) _*a*_ *π*_*ā*_(*a s*) *P* (*s*^′^ *s, a*), in which a selected option contributes only the primitive action its policy *π*_*ā*_ takes at the current state (for a primitive action *a, π*_*a*_(*a*^′^ *s*) = 1(*a*^′^ = *a*)), so that the persistence and termination of options are ignored. HSR and fHSR agents instead use the HSR with respect to *µ* (see below). All agents were instantiated with *ϵ*-greedily (*ϵ* = 0.1). A selected option ran until termination. After each action, primitive or option, the readout weights of the executed action were updated by Q-learning, extended to options by semi-Markov Q-learning ^19^, in which the TD error accumulates the discounted reward over the option’s execution,

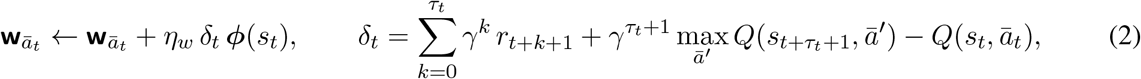

where the action *ā*_*t*_ initiated in state *s*_*t*_ lasts *τ*_*t*_ + 1 primitive steps (*τ*_*t*_ = 0 for primitive actions, for which this becomes standard Q-learning), *r*_*t*+1_ is the reward received on entering *s*_*t*+1_, and the bootstrap term is zero once the goal is reached. We used *η*_*w*_ = 0.1 and *γ* = 0.9. The readout weights were reset to zero at each change of goal, so that only the state representation could carry information between tasks.

We used three learning regimes. In the main regime (Figure 1; Extended Data Figures 3, 4 and 8), the representation was learned incrementally across tasks. Every agent used one-hot features in the first task, in which it therefore behaved as the naive or options agent with the same action set. At the end of each task *k*, the map **M**^(*k*)^ under the greedy policy learned in that task (the SR or the HSR, depending on the agent) was computed in closed form (Equations 5 and 7), and the representation was updated as a running average over the completed tasks,

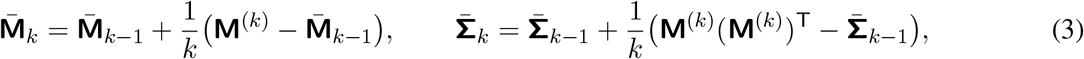

that is, a sample-average estimate (step size 1*/k*) of the task-averaged map, updated once per task. The rows of 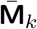 served as the features of SR and HSR agents in task *k* + 1. Note that the fSR + options and fHSR agents used the factorisation re-extracted from the averaged predictive similarity 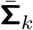 (Equation 10 with *p*(*g*) uniform over the completed tasks). The representation was held fixed within a task. In the second regime (Extended Data Figure 5), the SR and HSR were instead learned online by temporal-difference (TD) learning, with a given option set, starting from the identity (one-hot features) and carried over both between and within tasks. The SR was learned with the one-step TD rule ^4^, applied at every primitive transition, including those made during option execution,

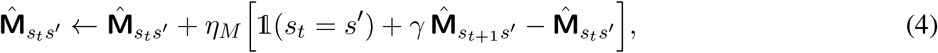

and the HSR with its option-level counterpart (Equation 8), applied whenever an action terminated, both with learning rate *η*_*M*_ = 0.1. In the third regime (Extended Data Figure 6), the options were themselves constructed online ^23^. During 10 warm-up episodes of random exploration, the agent recorded the primitive transitions it experienced in a subjective transition graph (Extended Data Figure 6a); eigenoptions were then constructed as described below, from a separate SR estimate of this graph (options and HSR agents) or from the SR that also served as the state representation (SR + options agents; Extended Data Figure 1h-j), added to the action set and reconstructed every 10 episodes as experience accumulated.

Learning efficiency is quantified through the area between the learning curve and the optimal path length, 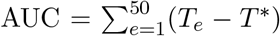, where *T*_*e*_ is the number of steps in episode *e* and *T* ^*^ the optimal number of steps (Figure 1m). Representational change is the relative squared Frobenius distance between the representations used in successive tasks, 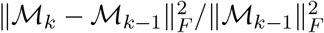, and analogously for the SR (Extended Data Figure 3h). In the online regime, the same quantity was computed between successive episodes (Extended Data Figure 6d) and summarised by its area within each task (Extended Data Figure 6e). Low-dimensional embeddings of state features were computed with two-dimensional t-SNE^56^, with perplexity 30.

### Successor representation

For a policy *π* with state-transition matrix **T**_*π*_, the SR is the expected discounted future occupancy of each state ^4^,

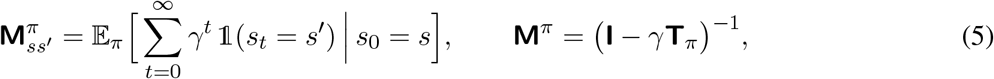

so that values factorise as **V**^*π*^ = **M**^*π*^**R** and can be recomputed when the reward changes. The random-walk SR (RW-SR), **M**_0_ = (**I** − *γ***T**_0_)^−1^, is the SR under the uniform policy. On a linear track, its entries decay exponentially with distance over a single correlation length (Supplementary Information, Section 2.2). Its columns served as the single-field RW-SR place-cell control ^3^.

### Eigenoptions and option selection

An option ^19^ is defined via a tuple, *O* = ⟨*I*^O^, *π*^O^, ***β***^O^⟩, encompassing the initiation set (*I*^O^) where the option *O* can be executed, the intra-option policy (*π*^O^), and the termination probability (***β***^O^ ∈ [0, 1]^|S|^). Options were constructed as eigenoptions ^27^ unless stated otherwise, which are given by the eigendecomposition of the RW-SR, **M**_0_**v**^(*i*)^ = *ζ*_*i*_**v**^(*i*)^, with eigenvalues ordered as *ζ*_0_ ≥ *ζ*_1_ ≥ · · · ≥ *ζ*_*N*−1_. Each eigenbasis *i* defines an options, with the eigenvector value (real, if necessary) defines a pseudo-reward function, 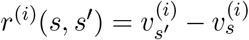. The option-specific policy follows the greedy policy with respect to the pseudo-action values, 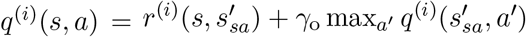, computed by value iteration, where 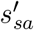 is the state reached by taking the primitive action *a* in *s* and *γ*_o_ is the discounting factor for computing the option-specific policies. The policy terminates whenever no further positive pseudo-return is available, that is, its termination vector has entries 1 (max_*a*_ *q*^(*i*)^(*s, a*) ≤ 0), which places its terminal states at the local maxima of **v**^(*i*)^. In compartmentalised environments these lie at corners and doorways (Extended Data Figure 2a,b). Note that the leading eigenvector, **v**^(0)^, is constant and defines a flat pseudo-reward, hence was excluded (Supplementary Information, Section 7.1).

The number of options is not set by hand. Instead, it is determined by the same criterion in every environment (the four-room environments, the linear tracks and the community graph): the explained-variance ratio of the RW-SR eigenvalue spectrum: we retained the eigenvectors **v**^(1)^, …, **v**^(*K*)^, with

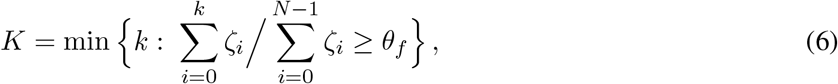

where *θ*_*f*_ = 0.95 unless stated otherwise (Extended Data Figure 2e). Because the spectrum of the RW-SR depends on the size of the environment, larger environments recruit more options. On a linear track, the eigenvectors are standing waves, so the option of index *i* partitions the track into segments of length *L/i* and terminates on a regular lattice (Extended Data Figures 2d and 10b). The criterion then makes *K* grow in proportion to *L*, so that the finest option scale is independent of track length (Supplementary Information, Section 7.1). The threshold *θ*_*f*_ and the option discount *γ*_o_ are the only parameters of option construction, and both were fixed across environments and analyses, so that the number of options changes between environments only through Equation 6. To test robustness of the place cell spatial tunings modelled by fHSR with respect to the number of options used for constructing the HSR, we varied *θ*_*f*_ between 0.8 and 0.99 (Extended Data Figure 12a), set the number of options directly (*K* = 4, 8, 16 or 32 on the 48-m track, Extended Data Figure 9b; *K* = 2, 4, 8, 16 or 32 in the large four-room environment, Extended Data Figure 4a-d) and varied *γ*_o_ (Extended Data Figures 9c; 12b). As controls, we replaced eigenoptions with options that each lead along shortest paths to a single terminal state, with terminal states chosen to be maximally dispersed across the environment (maximum-dispersion options; Extended Data Figure 4e-h), or, on tracks, with options of a single spatial scale that move the agent 3, 6, 12 or 24 m (Extended Data Figure 10d-g).

### Hierarchical successor representation

The HSR is the expected discounted future occupancy under a hierarchical policy *µ* that selects amongst primitive actions and options ^23^. For each action in the augmented action space, *ā* ∈ *Ā* = *A* ∪ *O*, it contributes to the computation of HSR given its intra-option occupancy **M**^*ā*^ (the SR of the action, or option’s policy until it terminates) and its discounted termination kernel **F**^*ā*^ = **M**^*ā*^ diag(***β***_*ā*_), where ***β***_*ā*_ is its termination vector; and a primitive action can be interpreted as an option of unit duration. The HSR satisfies the following semi-Markovian Bellman recursion ^23^.

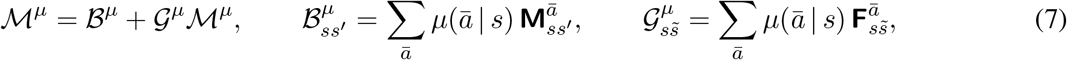

in which *B*^*µ*^ collects the occupancy accrued while options execute, and *G*^*µ*^ passes control, in a single step, from the state where an option begins to the state where it ends. The associated operator is a contraction for *γ <* 1, so the HSR is its unique fixed point, *M*^*µ*^ = (**I** – *G*^*µ*^)^−1 *Bµ*^, which we used for closed-form computation ^23^. Given the Bellman equation, the HSR can be learned online with a TD error that accumulates over the executed action.

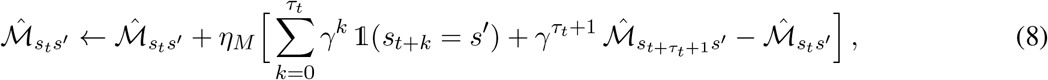

where the action initiated at time *t* lasts *τ*_*t*_ + 1 primitive steps, as in Equation 2, and *η*_*M*_ is the learning rate. The target accumulates the discounted occupancy along the executed option and bootstraps from the state in which it terminates; for primitive actions (*τ*_*t*_ = 0), Equation 8 reduces to the one-step TD rule of the SR (Equation 4). We write *M*^*g*^ for the HSR under the optimal hierarchical policy *µ*_*g*_ learned for task *g*, and denote the HSR under the uniform high-level policy *µ*_0_, which selects uniformly among the primitive actions and the options available at each state, as the random-walk HSR (RW-HSR). On a linear track, uniform selection among the eigenoptions produces jumps whose lengths are distributed as a power law above the finest option scale (Extended Data Figure 10a-c), so that predicted occupancy decays logarithmically rather than exponentially with distance and extends over a horizon far beyond the SR (Supplementary Information, Section 2.3).

### Factorised HSR: overcomplete non-negative decomposition of the HSR

We modelled place cells are the columns of the factorised HSR (fHSR), a non-negative basis 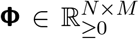, with *M* ≫ *N*, shared across a distribution of tasks *g* ∼ *p*(*g*). For computing the fHSR on linear tracks, *p*(*g*) assigns equal probability to running rightward (*µ*_→_), running leftward (*µ*_←_) and a random walk (*µ*_0_) (Extended Data Figure 9a); *µ*_→_ and *µ*_←_ are the greedy hierarchical policies of agents trained, as above, to run to the right and to the left end of the track. The basis minimises the linear reconsutrction error of every task’s HSR with task-dependent optimal loadings **W**^*g*^, with ridge-regularisation.

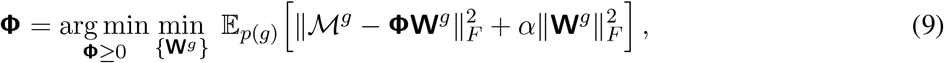

Eliminating the optimal loadings reduces this objective (Supplementary Information, Section 3.1), up to a constant, the factorisation loss objective reduces to the following.

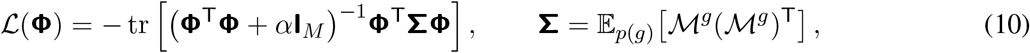

which depends on the HSRs only through **Σ**, the similarity between the futures predicted from different states. For isotropic reward distributions across *g* ∼ *p*(*g*), *L* is also, up to scale and a constant, equivalent to the expected error of linearly reconstructing the value functions of all tasks with a shared basis, which makes the basis optimal for linear value approximation across tasks (Supplementary Information, Section 3.3). We note that the ridge term is essential: without it, we would end up in a degenerate scenario where every basis that spans the state space would attain the same loss when *M > N* (Supplementary Information, Section 3.5).

To remove the scale ambiguity between basis and loadings, and to model each cell’s firing propensity explicitly, we express **Φ** = **U** diag(**a**), with unit-norm, non-negative atoms **u**_*m*_ and non-negative gains *a*_*m*_, and minimised

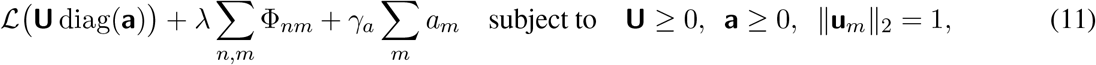

with *L*_1_ penalties on the basis (*λ*) and on the gains (*γ*_*a*_). Without the gains, normalisation would force nearly every cell to express a field. Gvein the close analogy between the loss loss obejctive and the symmetric non-neagtive matrix factorisation (symmetric NMF; Supplementary Information, Section 3.2.3), we used a heuristic initialisation scheme such that the basis is initialised given the symmetric NMF of the predictive similarity,

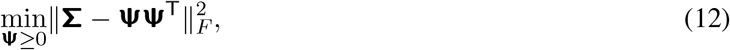

which shares its data term with *L* (Supplementary Information, Section 3.2.3), is a relaxation of kernel *k*-means clustering ^57^ and can be computed online by a network with Hebbian feedforward and anti-Hebbian lateral plasticity ^49^. We then refined the basis by multiplicative updates ^58^, which we refer to as quasi-NMF: because the regularised inverse in makes its gradient sign-indefinite, the gradient was split elementwise into positive and negative parts, and the atoms were updated along the Riemannian gradient on the unit sphere ^59^ (Supplementary Information, Section 3.2). Refinement was iterated until the relative change in between consecutive iterations fell below 10^−4^. To quantify its effect (Extended Data Figure 11), we compared the refined basis on the 48-m track with its initialisation (10 random seeds) in terms of the relative change in the total objective and in the trace loss, the cosine similarity between each basis vector and its own initialisation, and the field statistics.

Unless stated otherwise, we used *M* = 2,000 cells, *α* = 0.2, *λ* = 0.1, *γ*_*a*_ = 0.007, *θ*_*f*_ = 0.95, an option discount of *γ*_o_ = 0.99 and an HSR discount of *γ* = 0.9. These values were identical on every track, for both species and in all analyses, so that only the number of states, and with it the number of options selected by Equation 6, changed with track length. Sensitivity analysis with respect to these hyperparameters were performed by varying one parameter at a time around these defaults (number of options or *θ*_*f*_, *γ*_o_, *M, γ*_*a*_ and *λ*; Extended Data Figures 9b-f; 12a-d). The factorised SR (fSR) was obtained by applying the identical pipeline to the random-walk SR, and raw HSR columns, the columns of the HSR without decomposition, served as a second control. In the navigation experiments, the fHSR and fSR were obtained by applying the same decomposition to the four-room HSR and SR, with *M* = 2*N* . Extended Data Figure 8 compares this default with full-rank (*M* = *N* ) factorisations without (free-form) and with the non-negativity constraint, and with an overcomplete non-negative factorisation with *M* = 1.5*N* .

### Firing fields and field statistics

The tuning curve of model cell *m* is *f*_*m*_(*x*_*n*_) = *a*_*m*_*U*_*nm*_, scaled to a firing rate by a single global factor (1 Hz per unit); recorded cells contributed their empirical tuning curves. Tuning curves were smoothed with a Gaussian kernel (*σ* = 2 spatial bins) and thresholded at 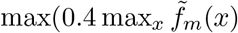, *r*_floor_), with *r*_floor_ = 0.35 Hz. Contiguous supra-threshold intervals at least 0.2 m wide were taken as fields, intervals separated by less than 0.2 m were merged, and cells without fields were counted as inactive. From each cell’s fields we computed the field count, the field locations normalised by track length, the within-cell field-width ratio (largest/smallest, for cells with at least two fields) and the coverage (summed field width divided by track length), and, across cells, the proportion of active cells. Model and recorded tuning curves were processed identically. To test robustness, the smoothing kernel width, the absolute and relative thresholds and the minimum field width were varied one at a time, and the rat data were re-analysed with the same parameters (Extended Data Figures 9g-k; 12e-g).

In the gamma-Poisson description ^11,15^, a cell’s field count is Poisson-distributed with a cell-specific propensity drawn from a Gamma distribution, so that counts across cells are negative-binomial; the equal-Poisson description assigns all cells the same rate. Both were fitted by maximum likelihood and compared with likelihood-ratio tests with one degree of freedom. The two descriptions also predict the distributions of recruitment positions (the normalised location of each cell’s first field) and of interfield intervals pooled across cells: exponential under equal-Poisson, and heavy-tailed under gamma-Poisson (Lomax, for the intervals). These were fitted and compared in the same way. Excess dispersion of field counts (Extended Data Figure 9) was quantified as the Fano factor (variance divided by mean) minus 1. Scaling exponents and values at 48 m are the slopes and intercepts of least-squares fits of the logarithm of each statistic against the logarithm of track length relative to 48 m (Extended Data Figure 12).

### Bayesian decoding

Population spike counts were simulated along a trajectory given a stochastic rightward policy (probability of moving rightward and leftward are 0.95 and 0.05, respectively), by drawing *k*_*m*_ ∼ Poisson(Δ*tf*_*m*_(*x*)) in bins of Δ*t* = 10 ms, and position was decoded as the maximum of the posterior under a uniform prior ^60,61^,

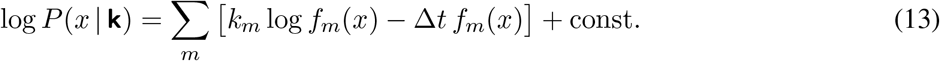

We report the mean and the 99th percentile (catastrophic error) of the absolute decoding error. All representations comprised the same number of cells (2000) and were scaled to the same mean population rate (5 Hz). Decoding was repeated on tracks of 40–200 m (10 random seeds per length). To assess the contribution of multi-scale cells, we compared a random 90% subsample of fHSR cells with versions lacking the decile of cells with the highest or the lowest within-cell field-width ratios.

### Population topology

Pairwise co-activity between neurons can be computed as the Pearson correlation between spatial tuning curves, or as the integral of the normalised spike-train cross-correlogram within±1,000 ms (Extended Data Figure 13a). The two co-activity matrices agree to a large extent over all sessions in the rat data, which is quantified by the Spearman correlation of their entries, and compared with a shuffled null (10,000 shuffles; Extended Data Figure 13b-d). Following refs. ^17,32^, cell pairs were ranked by co-activity and connected in order of decreasing co-activity, and the Betti numbers *β*_1_–*β*_3_ of the resulting clique complexes (*n*-degree holes for *n* ∈ [1, 2, 3]) were computed as functions of edge density up to 1 (Betti curves) and integrated over edge density (integrated Betti curves). Because Betti curves depend only on the rank order of co-activities, they are invariant to monotonic transformations and reflect the geometry of any space in which co-activity decreases with distance ^32^. Every population was subsampled to the number of simultaneously recorded rat cells in corresponding sessions. The hyperbolic reference sampled the same number of points uniformly from a three-dimensional hyperbolic ball of radius 15.5 and converted geodesic distances into co-activities by a decreasing function ^17^ (300 random seeds). Betti curves of the rat data were compared with those of populations in which each cell’s fields were relocated at random, preserving their number and widths (1,000 shuffles; Extended Data Figure 13h,i). The relationship between a cell’s number of fields *x* and its mean field width *y* was fitted by least squares with *y* = *b* + *a/x*, and *a* was tested with a two-sided Wald test.

### Population periodicity

Summed population activity, *A*(*x*) = Σ_*m*_ *f*_*m*_(*x*), was computed from smoothed, unthresholded tuning curves and normalised to its maximum. With *δA* its deviation from the mean, we computed the spatial autocorrelation and the spectral centroid,

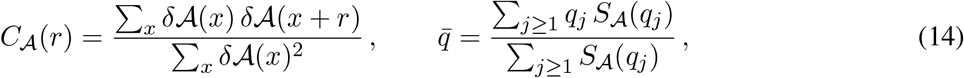

where *S*_*A*_(*q*_*j*_) is the power spectral density of *δA* at wavenumber *q*_*j*_ = 2*πj/L*. The characteristic wavelength is the smallest non-zero spatial lag at which *C*_*A*_ peaks. Analytically, the characteristic wavelength can be viewed as a proxy to the finest option scale (Supplementary Information, Section 7.3). Both measures were compared with 1,000 shuffles in which each cell’s fields were relocated at random, preserving their number and widths (one-sided empirical *p* values), and with 1,000 populations whose fields were placed independently, with counts drawn from equal-Poisson or gamma-Poisson processes matched to the observed statistics. Independent placement preserves each cell’s own contribution to the spectrum but removes the cross-cell term, so departures from these controls indicate alignment of fields across cells (Supplementary Information, Sections 7.2 and 7.4). Dependence on track length was assessed with two-sided Wilcoxon signed-rank tests across seeds (model) or rats, with two-sided Mann-Whitney *U* tests across bats, and with a least-squares linear fit and a linear mixed-effects model (LMM) of each measure on track length.

To test whether the periodicity reflected track geometry (Extended Data Figure 15a-f), we detected turns as peaks in the curvature of each track’s two-dimensional layout, with turning angles above 20^°^ (all turns) or 45^°^ (major turns). Summed activity was z-scored and band-pass filtered (0.2-0.7 cycles m^−1^), with the ends of the track excluded. Turn-triggered averages (±2 m) were compared with a null obtained by randomly shifting turn positions (1,000 shuffles) and with a positive control in which every field was shifted to its nearest turn. The phase of the band-passed activity at turns, obtained from its Hilbert transform, was tested for uniformity with Rayleigh tests, and its concentration was summarised by the mean resultant length. The same periodicity analyses were applied to the bat data on 130-200 m tracks (Extended Data Figure 15g-j).

### Random-function model of place-cell tuning

As a non-computational control, we simulated place cells with the random-function model of Mainali et al. 2025^16^. In this model, each CA1 cell sums the activity of many spatially tuned neurons through random synaptic weights. By the central limit theorem, this summed input, as a function of spatial location along the track, converges in distribution to a Gaussian process, a random function whose values at any set of locations are jointly Gaussian. The cell fires wherever its input exceeds a threshold, so its firing fields are the stretches of track where the input happens to be high. The tuning curve of cell *m* is

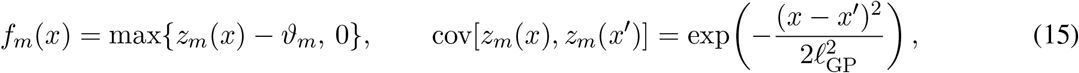

where *z*_*m*_ is a zero-mean, unit-variance Gaussian process, sampled independently for each cell, *l*_GP_ is its correlation length and *ϑ*_*m*_ is the threshold. A longer correlation length gives fewer, wider fields, and a higher threshold gives fewer, narrower ones. Because cells are sampled independently, the model describes the tuning of each cell but no relationship between cells.

We used the same parameters the original paper ^16^ fitted for the rat recordings on the 48-m track ^11^ by matching the mean field size and the mean number of fields per cell: *l*_GP_ = 0.34 m and *ϑ*_*m*_ = 1.8 for all cells (fixed threshold). We also used their extension in which the threshold varies across cells, drawn from a skew-normal distribution restricted to positive values, with mean 1.8 and spread and skewness as in ref. ^16^ (variable threshold). For each random seed, we sampled 1000 cells on the spatial bins of the 48-m track. Tuning curves were either used directly (noiseless) or, to mimic the noise of rate maps estimated from recorded spikes, and re-estimated from Poisson spike counts generated from them (Poisson). We analysed three configurations (fixed threshold, noiseless; fixed threshold, Poisson; variable threshold, Poisson), each over 300 random seeds. Betti curves were computed as for the other populations, and the integrated Betti curves of the rat data were compared with their distribution across seeds by two-sided permutation tests (Extended Data Figure 13j,k). Summed activity was analysed as described under Population periodicity, with 1000 shuffles (Extended Data Figure 15k).

### Option terminal states in two-dimensional environments

For each environment in Extended Data Figure 14, we computed the density of option terminal states and the normalised betweenness centrality of each state ^62^, which is high at bottlenecks, and classified states as doorway, door flank, corner, wall, near wall or interior. We compared the fraction of option terminal states with the fraction of all states in each class. The fHSR and fSR were obtained by the decomposition above from the RW-HSR and the RW-SR of each environment. Their summed activity, normalised to the median of each room, was analysed as a function of the distance from the nearest doorway and compared between the two bases within each state class with two-sided Wilcoxon signed-rank tests across random seeds.

### Non-spatial community structure

We computed the RW-SR and the RW-HSR on the 15-node community-structured graph of refs. ^7,29^, using the eigenoptions selected by Equation 6 (with *θ*_*f*_ = 0.99; 8 options; Extended Data Figure 7). Node representations (the columns of each matrix) were embedded in two dimensions by multidimensional scaling ^63^, and pairwise distances between embedded nodes were compared with those derived from bilateral hippocampal BOLD activity ^7^, with numerical values estimated from the published figures from Stachenfeld et al. 2017^3^, by Spearman correlation and mean squared error. Community structure was summarised by the ratio of the mean between-community to the mean within-community distance, which we also computed for the RW-HSR as a function of the number of eigenoptions.

### Experimental datasets

We reanalysed previously published recordings; no new animal experiments were performed, and all procedures were approved by the relevant institutional animal care and use committees, as described in the original publications ^11,12,14^. Rat CA1 recordings were obtained from 5 rats running on 10-, 24- and 48-m linear tracks (253 cells; 8 sessions on the 48-m track) ^11^, and bat CA1 recordings from 14 Egyptian fruit bats flying along linear tracks of 15, 130, 180 and 200 m (621 cells) ^12,14^. In both datasets, direction-specific tuning curves were computed from periods of active movements (rats, ≥ 5 cm/s) or flight (bats, ≥ 1 m/s), with occupancy normalisation and spatial bins of 0.1 m (rats) and 0.2 m (bats), and all field, topology and periodicity analyses were applied identically to model and data. The numbers of cells entering each analysis are given in the figure legends.

### Statistics and reproducibility

Unless stated otherwise, comparisons used two-sided non-parametric tests: the Wilcoxon signed-rank test for paired and the Mann-Whitney *U* test for unpaired comparisons. Nested distributional comparisons used likelihood-ratio tests, and fitted slopes were tested with two-sided Wald tests. Linear mixed-effects models of a measure on track length included a random intercept for each seed (model) or animal (data), and their fixed-effect slope *β* was tested with a two-sided Wald test. Rank correlations were quantified by Spearman’s *ρ*, and circular uniformity was tested with Rayleigh tests. Unless stated otherwise, shuffle-based tests report empirical *p* values from 1,000 shuffles, so that *p <* 0.001 indicates that no shuffle was as extreme as the observed value. Exact *p* values are reported, except below the floating-point range (*p <* 10^−308^). No correction for multiple comparisons was applied unless stated. The numbers of seeds, animals and cells are given in the figure legends. No statistical methods were used to predetermine sample sizes; the sizes of the experimental datasets were determined by the original studies.

## Data availability

The rat CA1 data are publicly available from CNCRS^64^. Requests for access to the bat CA1 data ^12,14^ should be directed to the corresponding authors of those papers.

## Code availability

Simulation and analysis code will be made available in a public repository upon publication.

## Acknowledgements

C.Y. was supported by a Leverhulme Early Career Fellowship (ECF-2026-076). M.L. was supported by a Wellcome Trust Investigator Award in Science (212262/Z/18/Z). M.L. and N.U. were supported by the Simons Collaboration on Ecological Neuroscience (SFI-AN-NC-SCN-00007276). A.K.L. was supported by the Howard Hughes Medical Institute.

## Author contributions

C.Y. conceptualised the project. C.Y. performed the mathematical derivations and model simulations, with input from M.L. C.Y. performed data analysis with input from P.D.R. and A.K.L. (rat data), and S.R.M. and N.U. (bat data). C.Y., P.D.R., A.K.L., N.U. and M.L. discussed and interpreted the results. P.D.R. and A.K.L. collected the rat data. S.R.M., T.E., L.L. and N.U. collected the bat data. C.Y. and M.L. wrote the manuscript.

## Extended Data Figures

**Extended Data Figure 1:**
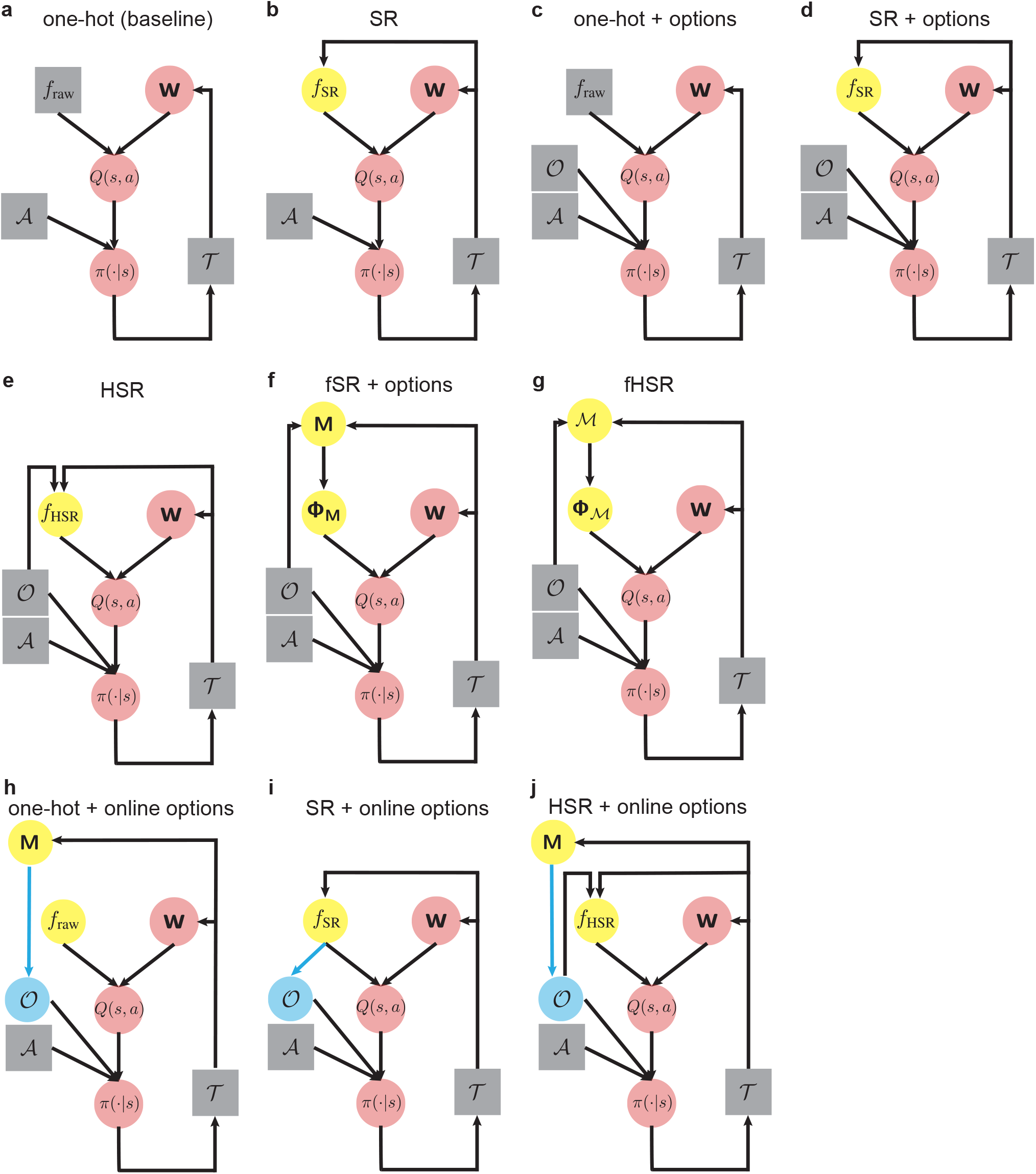
Reinforcement-learning agents with different state representations and action spaces. **a**. Baseline agent with one-hot state features (*f*_raw_; naive; Figure 1c,d). The action-value function *Q*(*s, a*) is a linear function of the state features with weight vector **w**. The behavioural policy *π*(·|*s*) is *ϵ*-greedy with respect to *Q* and selects primitive actions (*A*), which generate transitions *T* = (*s, a, s*^*′*^, *r*) that are used to update **w**. At each change of goal, **w** is reset to zero, which simultaneously resets the value function and the policy (red); the baseline agent therefore has no source of transfer. **b**. SR agent (Figure 1e,f), whose state features (*f*_SR_) are given by the SR, which depends on the agent’s behaviour and is learned from experienced transitions: incrementally across tasks, as the running average of the SRs under the policies learned for the completed tasks (Figure 1; Extended Data Figures 3, 4 and 8), or online by temporal-difference learning on every transition (Extended Data Figures 5 and 6). **c**. Agent with one-hot state features and options (*O*; options; Figure 1g,h), whose hierarchical policy selects among primitive actions and options. **d**. SR agent with options (SR + options; Extended Data Figure 3a,b). In this case, the SR is computed from the one-step transition dynamics induced by the hierarchical policy (Methods). **e**. HSR agent (Figure 1i,j), whose state features (*f*_HSR_) depend on both the primitive actions and the option set at the level of compound actions. **f**. Factorised SR agent with options (fSR + options; Extended Data Figure 3c,d). The agent computes the SR (**M**) at the level of one-step transitions, as the SR + options agent does, and its state features are given by the factorised SR (**Φ**_**M**_), the overcomplete, non-negative decomposition of the SR (Equation 1; Methods). **g**. Factorised HSR agent (fHSR; Figure 1k,l), whose state features are given, as for the fSR + options agent, by the factorised HSR (**Φ**_*M*_), the overcomplete, non-negative decomposition of the HSR (*M*). **h**. Agent with one-hot state features and online option construction (options; Extended Data Figure 6). The agent initially has access only to primitive actions, and its policy is effectively random. It maintains a separate estimate of the SR learned from primitive transitions (**M**). After a prespecified number of warm-up episodes, eigenoptions are derived from the eigenvectors of this estimate (blue arrow); they augment the action set and are periodically updated as experience accumulates. **i**. SR agent with online option construction (SR + options), in which options are constructed from the SR that simultaneously serves as the state representation. **j**. HSR agent with online option construction.

**Extended Data Figure 2:**
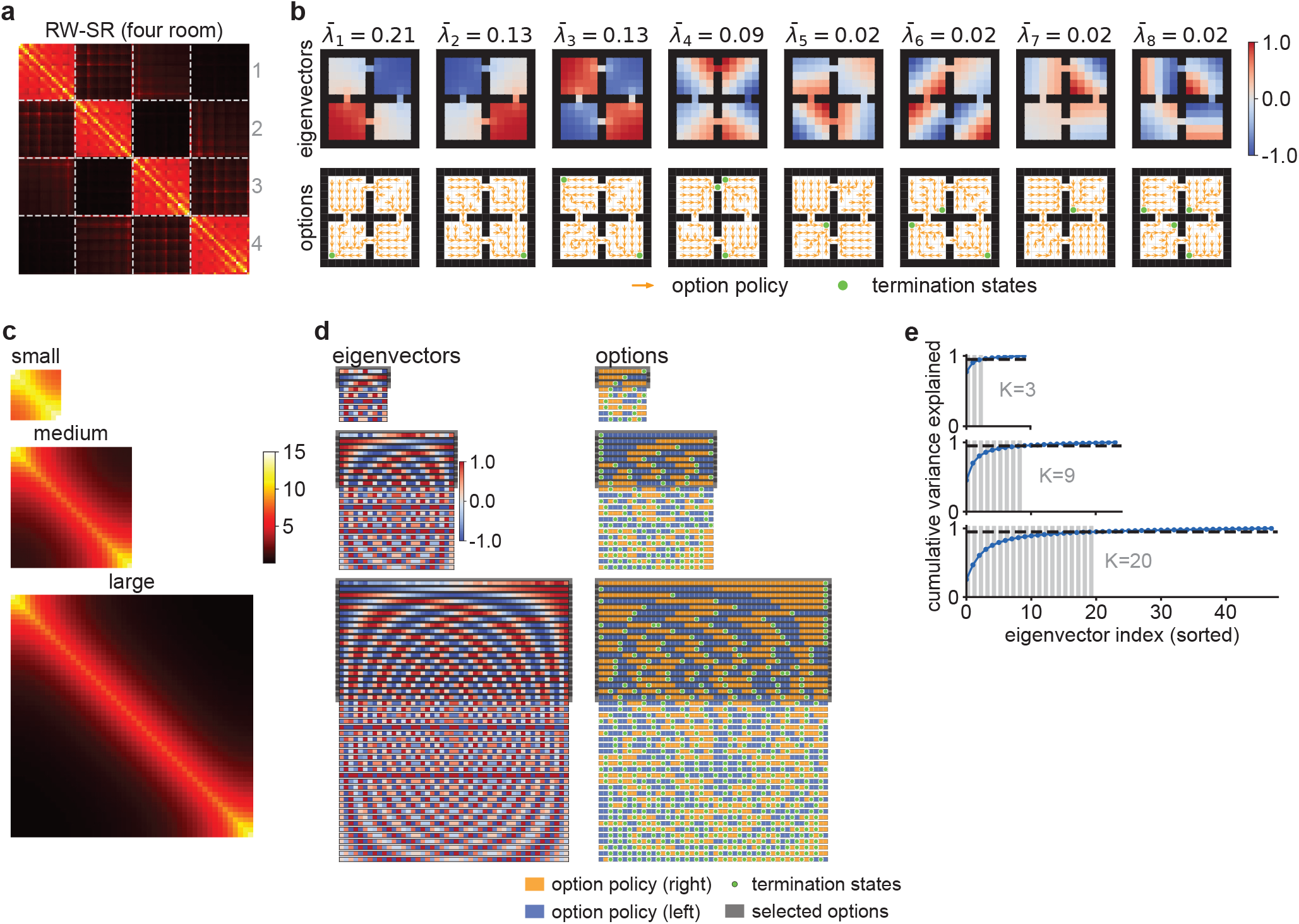
Eigenoptions identify topologically relevant subroutines from the spectral structure of the environment’s transition dynamics. **a**. Random-walk SR (RW-SR) matrix of the four-room environment, with rows and columns ordered by room (dashed lines; rooms 1-4). Warm colours indicate high expected future occupancy. **b**. Top, the eight leading eigenvectors of the matrix in a, shown as pseudo-reward functions over the environment and ranked by their normalised eigenvalue (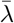, fraction of the eigenvalue spectrum). Bottom, the corresponding option policies (orange arrows) and terminal states (green), obtained by solving the MDP defined by each pseudo-reward. Each option drives the agent towards a local maximum of its pseudo-reward, which coincides with corners and doorways. **c**. RW-SR matrices of small, medium and large linear tracks (10, 24 and 48 states). **d**. Left, eigenvectors of the matrices in c, which are standing waves of increasing spatial frequency. Right, the corresponding option policies (orange, rightward; blue, leftward) and terminal states (green), which partition the track into progressively finer segments with increasing eigenvector index. Grey, options retained by the selection criterion in e. **e**. Cumulative fraction of the eigenvalue spectrum explained by the sorted eigenvectors. The number of options *K* is the smallest number of leading eigenvectors whose cumulative share exceeds 0.95 (dashed line), so that longer tracks recruit more options (*K* = 3, 9 and 20).

**Extended Data Figure 3:**
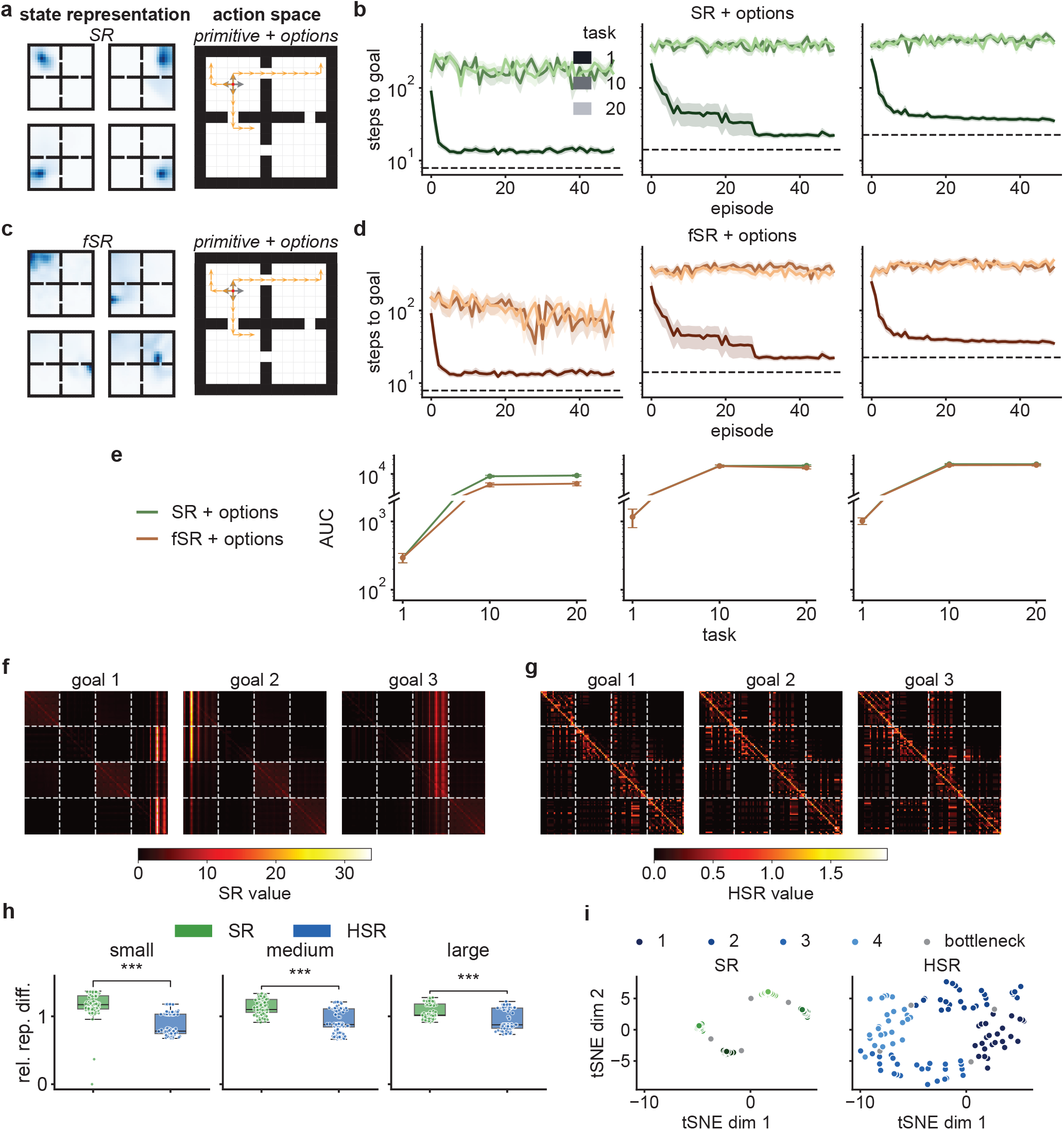
Learning curves of the remaining agent configurations, and stability and geometry of SR and HSR features. **a, c**. Example state features (left) and action spaces (right) of agents with SR features (SR + options; a) and with the factorised SR as state features (fSR + options; c), each with primitive actions and eigenoptions. **b, d**. Learning curves of the agents in a and c for tasks 1, 10 and 20, as in Figure 1d (mean ± s.e.m. over 10 random seeds for each of 9 goals). Dashed lines, optimal path length. **e**. AUC, as in Figure 1m, of the agents in a and c as a function of task in each environment. **f, g**. SR (f) and HSR (g) matrices under the policies learned for goals 1, 2 and 3, with states ordered by room (dashed lines, room boundaries). **h**. Relative difference between the representations of successive tasks (Methods) for SR (green) and HSR (blue) agents in each environment (two-sided Mann–Whitney *U* tests; small: *U* = 29,717.0, *p* = 5.8 × 10^*−*43^; medium: *U* = 26,518.0, *p* = 7.3 × 10^*−*26^; large: *U* = 25,340.0, *p* = 1.0 × 10^*−*20^; 3 pairwise comparisons per seed over *n* = 20 random seeds). Box plots as in Figure 6; points, individual comparisons. **i**. Two-dimensional t-SNE embeddings ^56^ of SR (left) and HSR (right) state features in the large environment, coloured by room (1-4) and with doorway (bottleneck) states in grey. \*\*\**p <* 0.001.

**Extended Data Figure 4:**
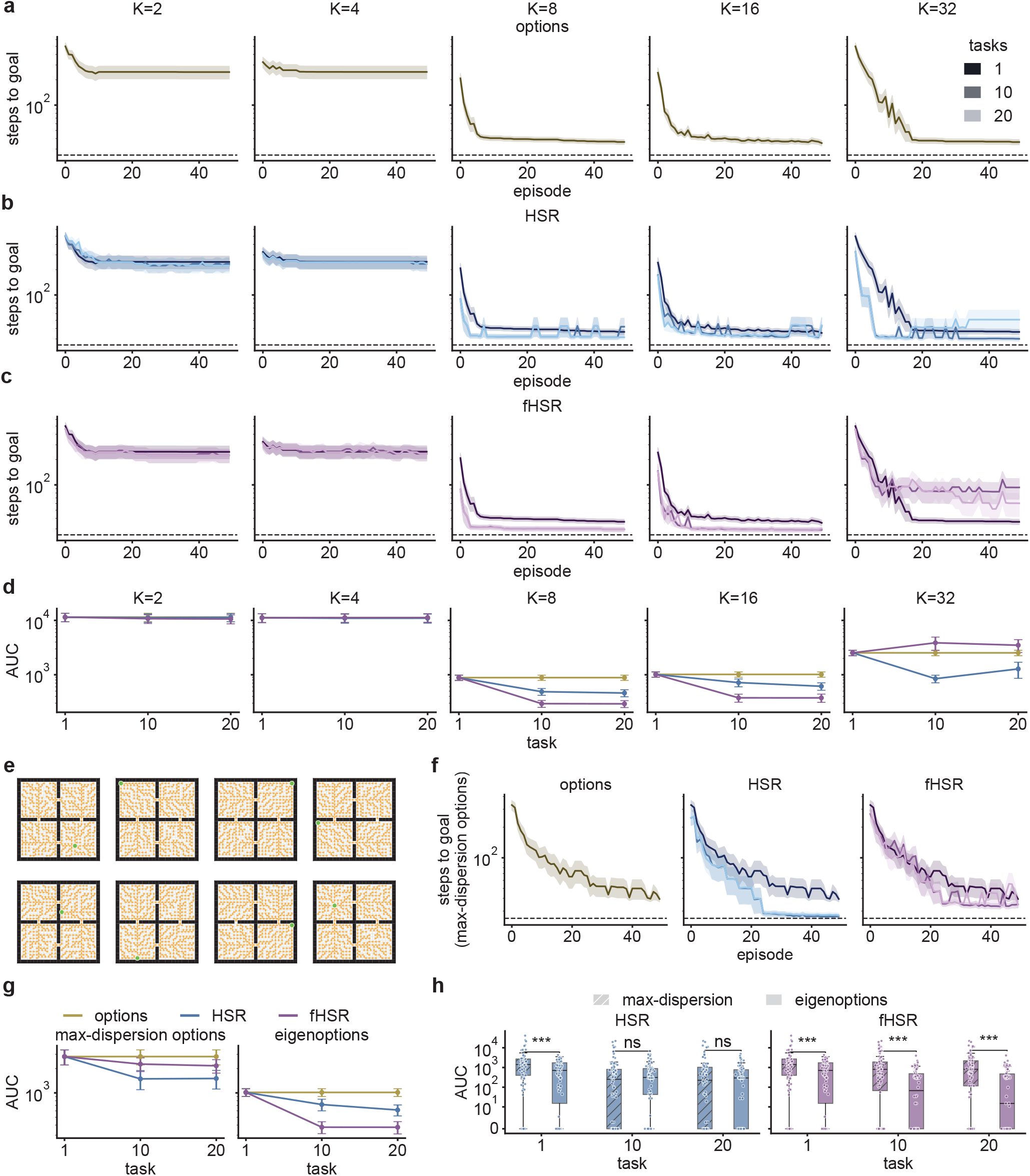
The navigational advantages of the HSR hold across a range of option numbers and for an alternative definition of options. **a-c**. Learning curves (steps to goal per episode; mean ± s.e.m., 10 random seeds per goal over 9 goals) for tasks 1, 10 and 20 in the large four-room environment for options (a), HSR (b) and fHSR (c) agents, each with primitive actions and *K* = 2, 4, 8, 16 or 32 eigenoptions (columns). Dashed lines, optimal path length. **d**. Learning curve AUC as a function of task for the three configurations in a-c, for each *K*. With two or four options, the three agents perform alike. **e**. Example option policies (orange arrows) and terminal states (green) of eight maximum-dispersion options, an alternative option class (Methods), in the large four-room environment. **f, g**. Learning curves (f) and AUC as a function of task (g; left), as in a-d, for agents with 16 maximum-dispersion options in the large four-room environment; right panel in g, the same agents with eigenoptions. HSR and fHSR agents retain their advantage in scalability and transfer. **h**. AUC of HSR (left) and fHSR (right) agents with maximum-dispersion options (hatched) and with eigenoptions in tasks 1, 10 and 20 (two-sided Mann–Whitney *U* tests; HSR task 1: *U* = 5128.0, *p* = 0.0008; task 10: *U* = 3683.5, *p* = 0.2898; task 20: *U* = 4156.5, *p* = 0.7570; fHSR task 1: *U* = 5128.0, *p* = 0.0008; task 10: *U* = 5666.5, *p* = 2.3491 × 10^*−*6^; task 20: *U* = 5855.0, *p* = 1.3932 × 10^*−*7^). Eigenoptions yield more efficient transfer than maximum-dispersion options in fHSR agents, but not in HSR agents. Box plots as in Figure 6; points, individual runs. \*\*\**p <* 0.001; ns, not significant.

**Extended Data Figure 5:**
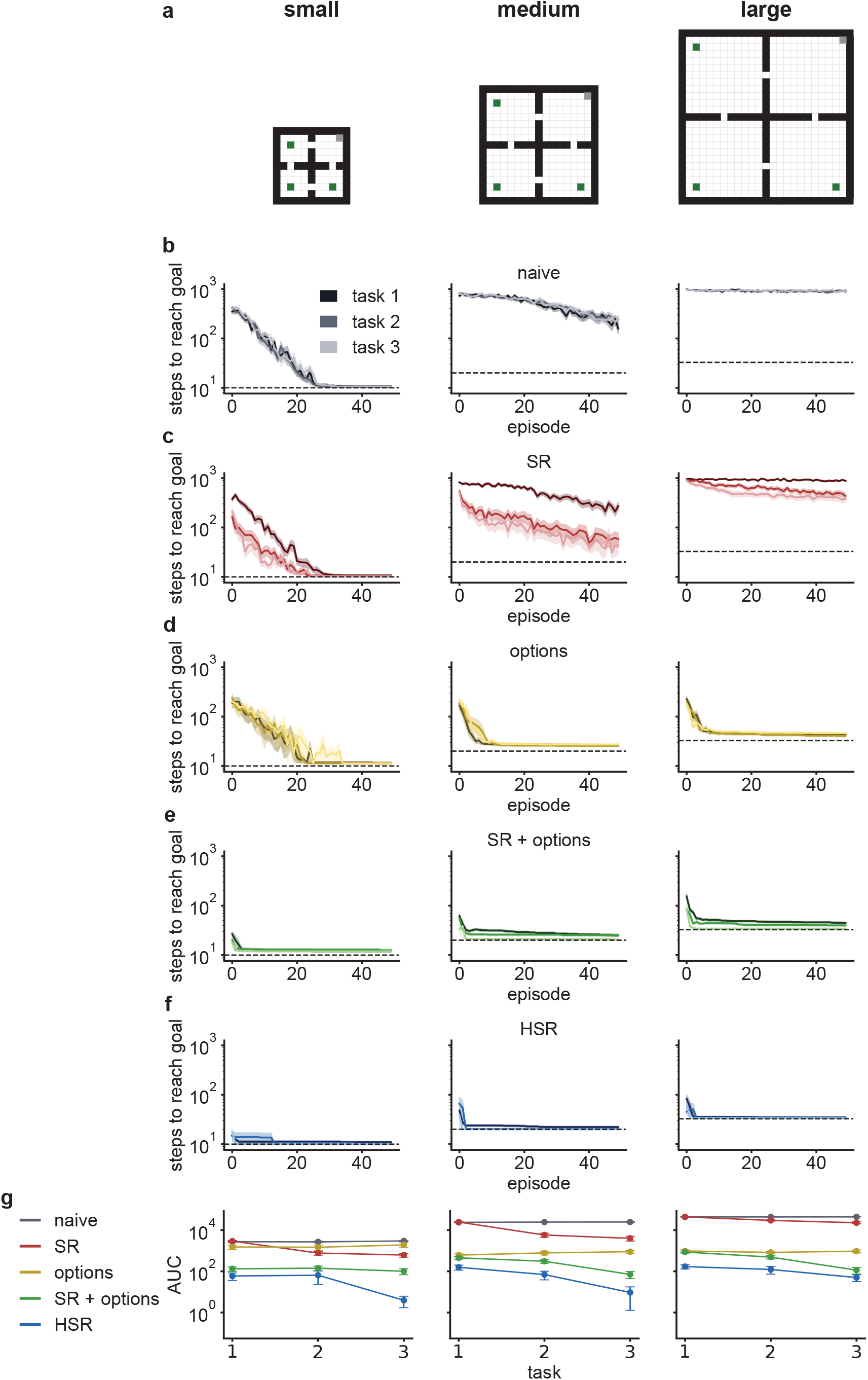
The navigational advantages of the HSR persist when state representations are learned online. **a**. Simpler transfer setting in the small, medium and large four-room environments: a start state (grey) and three goals (green), one per task, in all six orderings (Methods). **b-f**. Learning curves for tasks 1-3 of naive (b), SR (c), options (d), SR + options (e) and HSR (f) agents, with the SR and HSR learned online by temporal-difference learning and a given option set (Methods). Mean ± s.e.m., *n* = 60 runs (6 goal orderings × 10 random seeds). Dashed lines, optimal path length. **g**. Learning curve AUC of the five agents as a function of task in each environment.

**Extended Data Figure 6:**
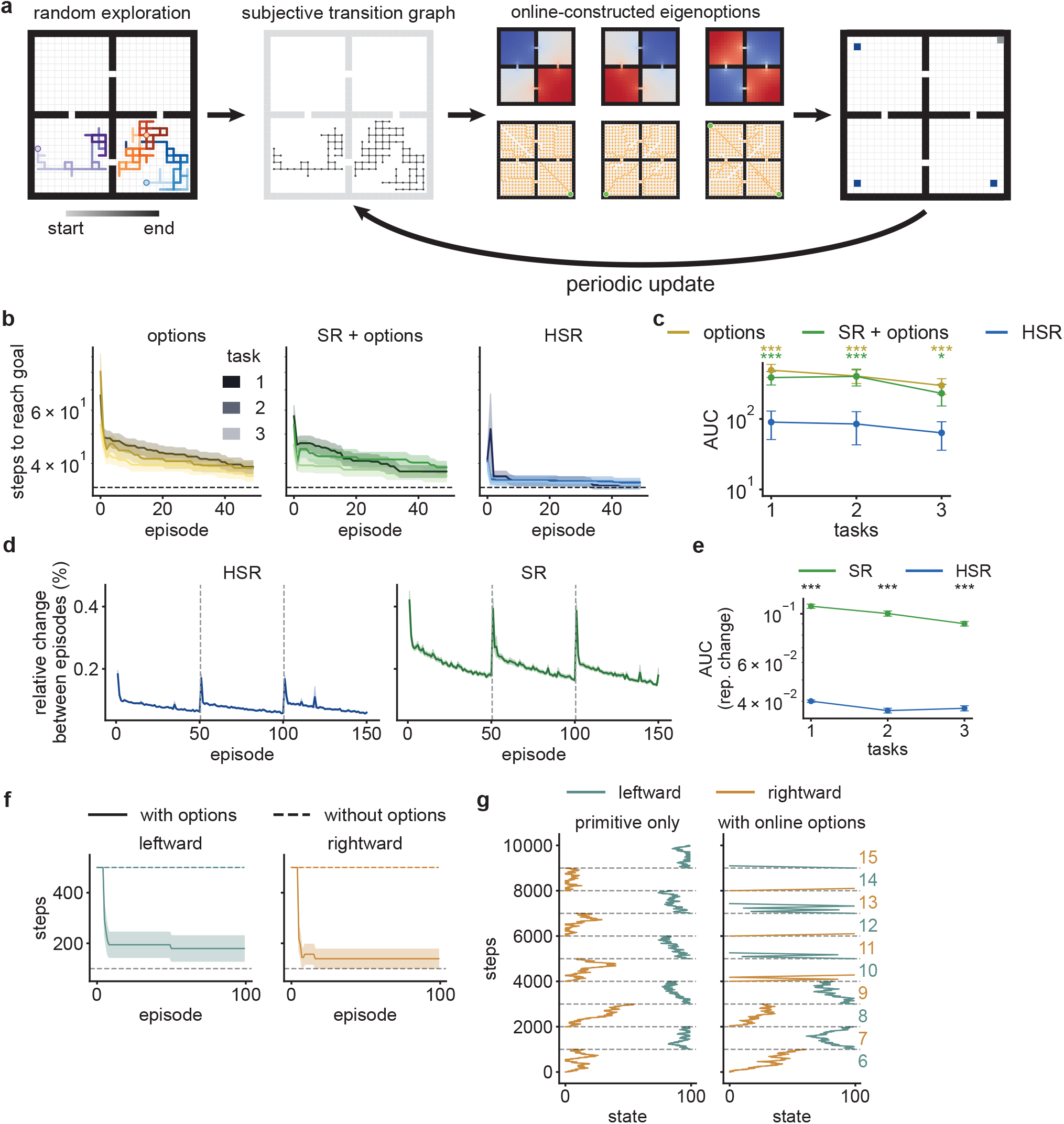
The HSR supports efficient transfer and ecologically plausible movement patterns under online-constructed options. **a**. Online option construction (Extended Data Figure 1h-j). During random exploration of the four-room environment, the agent builds a subjective transition graph, from which eigenoptions are constructed; these are added to the action set and periodically updated during subsequent goal-directed tasks. **b**. Learning curves (steps to goal per evaluation episode; mean ± s.e.m. over all six goal orderings with 10 random seeds each in the three-goal transfer setting; Extended Data Figure 5a) across tasks 1-3 for options (left), SR + options (middle) and HSR (right) agents, all with online-constructed options. Dashed lines, optimal path length averaged over goals. **c**. AUC of the three agents in b (mean ± s.e.m.; two-sided Mann–Whitney *U* tests; options versus HSR, task 1: *U* = 1,121.0, *p* = 4.4 × 10^*−*5^; task 2: *U* = 786.5, *p* = 2.9 × 10^*−*9^; task 3: *U* = 1,092.5, *p* = 7.6 × 10^*−*6^; SR + options versus HSR, task 1: *U* = 1,195.5, *p* = 2.0 × 10^*−*4^; task 2: *U* = 1,278.0, *p* = 4.0 10^*−*4^; task 3: *U* = 1,541.0, *p* = 0.034). **d**. Relative change in the HSR (left) and SR (right) between successive episodes; dashed lines, changes of goal (mean ± s.e.m.). **e**. Area under the representational-change curves in d within each task (AUC; two-sided Mann–Whitney *U* tests; task 1: *U* = 0.0, *p* = 3.6 × 10^*−*21^; task 2: *U* = 0.0, *p* = 3.6 × 10^*−*21^; task 3: *U* = 0.0, *p* = 3.6 × 10^*−*21^; *n* = 60 runs). **f**. Steps per episode for leftward (left) and rightward (right) runs on a linear track of 100 states, with (solid) and without (dashed) online-constructed options (mean ± s.e.m. over 10 random seeds). **g**. Example trajectories (position against cumulative steps) with primitive actions only (left) and with online-constructed options (right). Colours, running direction; numbers, episode index; dashed lines, episode boundaries. With options, the agent runs between the ends of the track in laps. \**p <* 0.05; \*\*\**p <* 0.001.

**Extended Data Figure 7:**
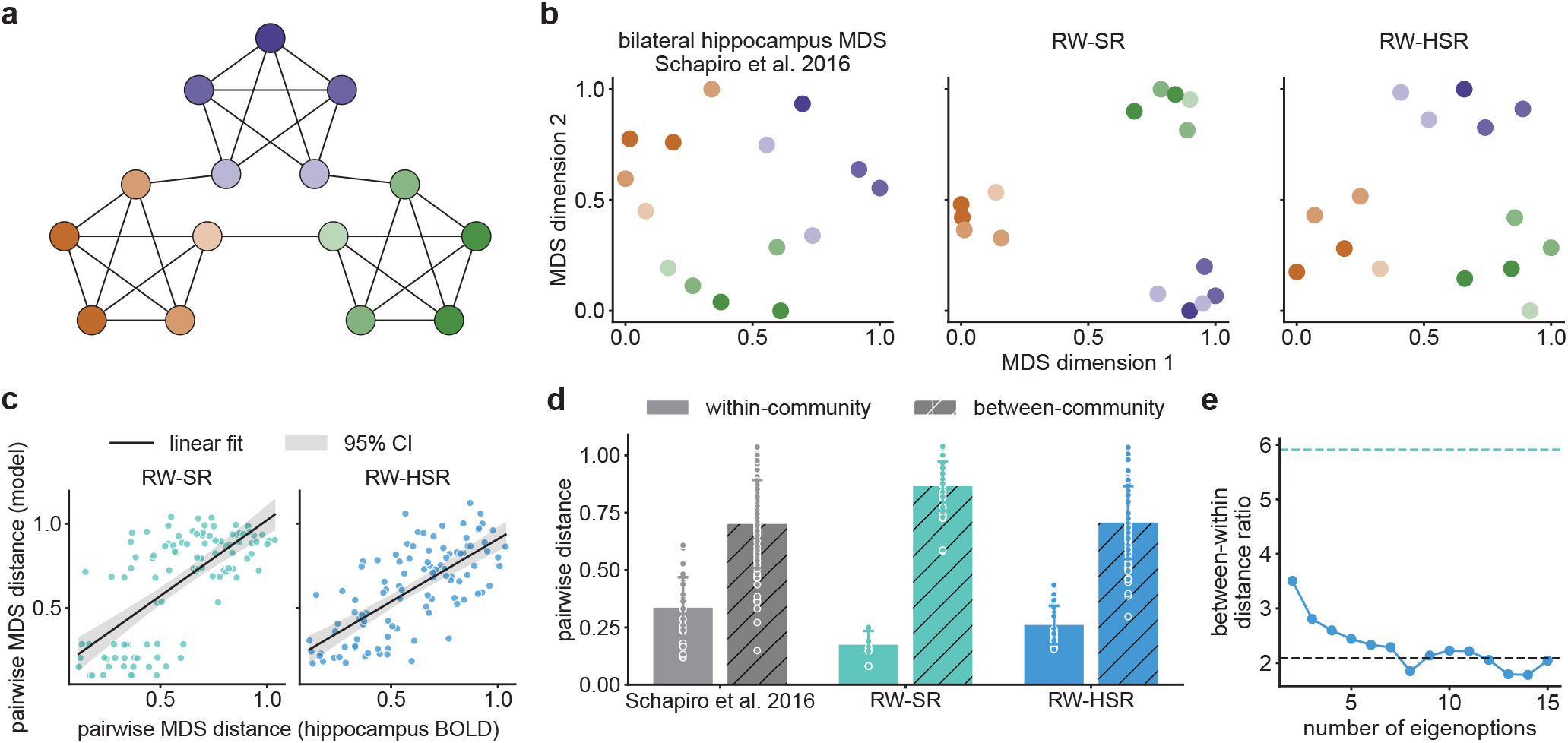
The HSR captures the geometry of hippocampal representations in a non-spatial community-structure task. **a**. Community-structured graph ^7,29^: 15 nodes, each associated with a distinct visual stimulus, arranged in three communities (colours) with dense within-community and sparse between-community connectivity. Human participants view sequences of stimuli generated by a random walk on the graph during functional magnetic resonance imaging (fMRI). We compared the predictive representations of this graph with hippocampal BOLD activity evoked by each node. **b**. Two-dimensional multidimensional scaling (MDS) ^63^ embeddings of node representations from bilateral hippocampal BOLD activity (adapted from ref. ^7^), and from the columns of the SR and HSR matrices under a uniformly random (high-level) policy (RW-SR and RW-HSR). The HSR uses three eigenoptions, selected with the criterion of Extended Data Figure 2e. **c**. Pairwise distances between embedded nodes for RW-SR (left) and RW-HSR (right) against the corresponding distances from hippocampal BOLD data. Lines and shading, least-squares fit and 95% confidence interval (Spearman correlations; RW-SR: *ρ* = 0.64, *p* = 2.2 × 10^*−*13^; RW-HSR: *ρ* = 0.70, *p* = 1.9 × 10^*−*16^; mean squared error, RW-SR: 0.049; RW-HSR: 0.032). **d**. Pairwise distances between embedded nodes within and between communities for hippocampal data ^7^, RW-SR and RW-HSR (mean ± s.d.; points, node pairs). The RW-HSR more closely matches the empirical structure, whereas the RW-SR over-separates communities while collapsing within-community structure. **e**. Ratio of between- to within-community distances for the RW-HSR as a function of the number of eigenoptions. Dashed lines, RW-SR (green) and hippocampal data (black). Increasing the number of options progressively preserves within-community variability, while the ratio stays near the empirical value.

**Extended Data Figure 8:**
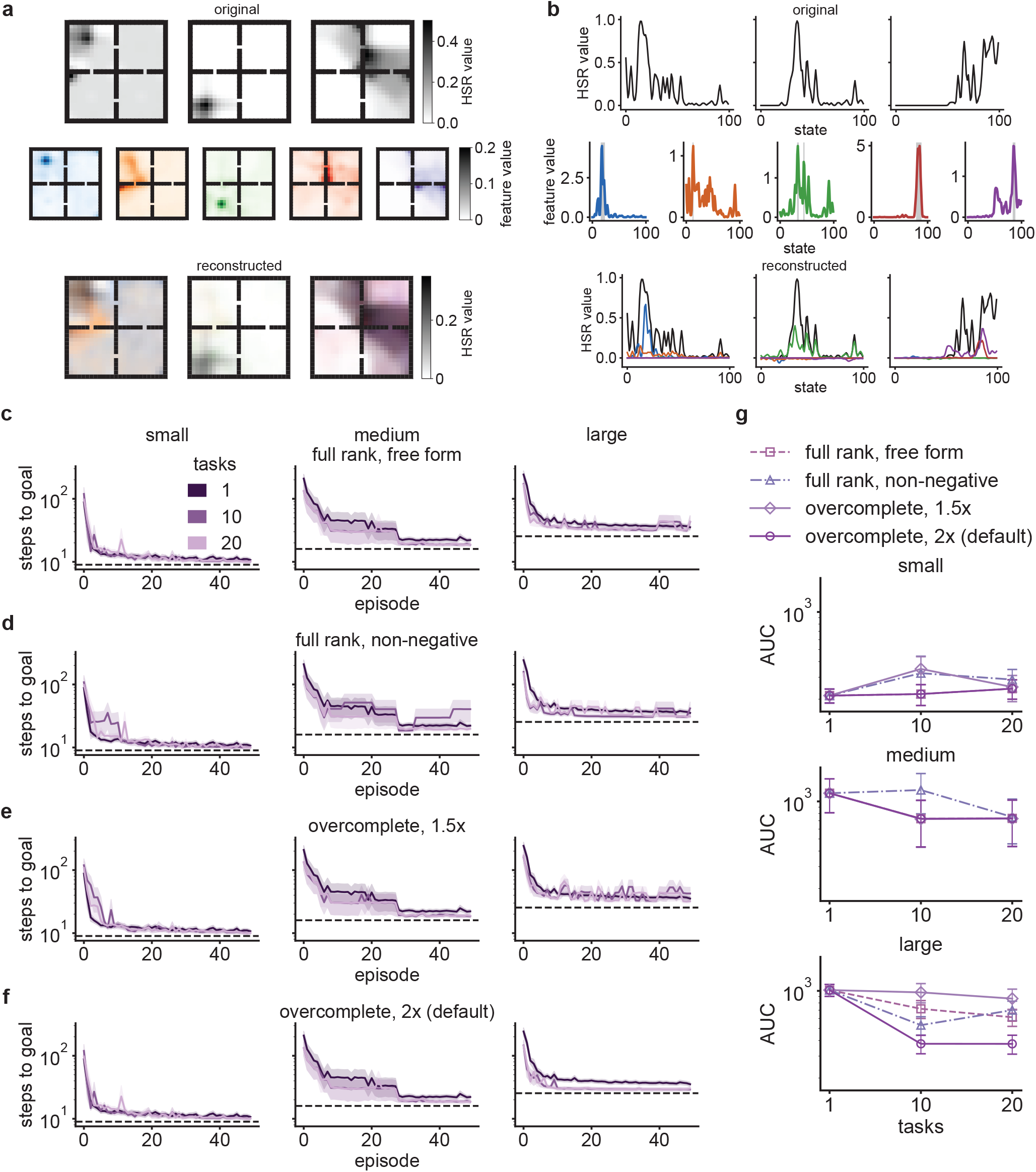
Non-negativity has little effect on the transfer advantage of the factorised HSR (fHSR), particularly at higher degrees of overcompleteness. **a**. Three example HSR columns in the four-room environment (top), five example vectors of the fHSR (middle; colours identify basis vectors) and the reconstruction of the HSR columns as weighted sums of basis vectors (bottom). **b**. Same as a, for a linear track of 100 states. **c-f**. Learning curves for tasks 1, 10 and 20 (as in Figure 1l; mean ± s.e.m. over 10 random seeds for each of 9 goals) of agents whose state features are factors of the HSR obtained by full-rank free-form factorisation (without the non-negativity constraint; c), full-rank non-negative factorisation (d) and overcomplete non-negative factorisation with 1.5 (e) and 2 (default; f; cf. Figure 1l) times as many factors as states. **g**. Learning curve AUC as a function of task for the agents in c-f in the small (top), medium (middle) and large (bottom) environments.

**Extended Data Figure 9:**
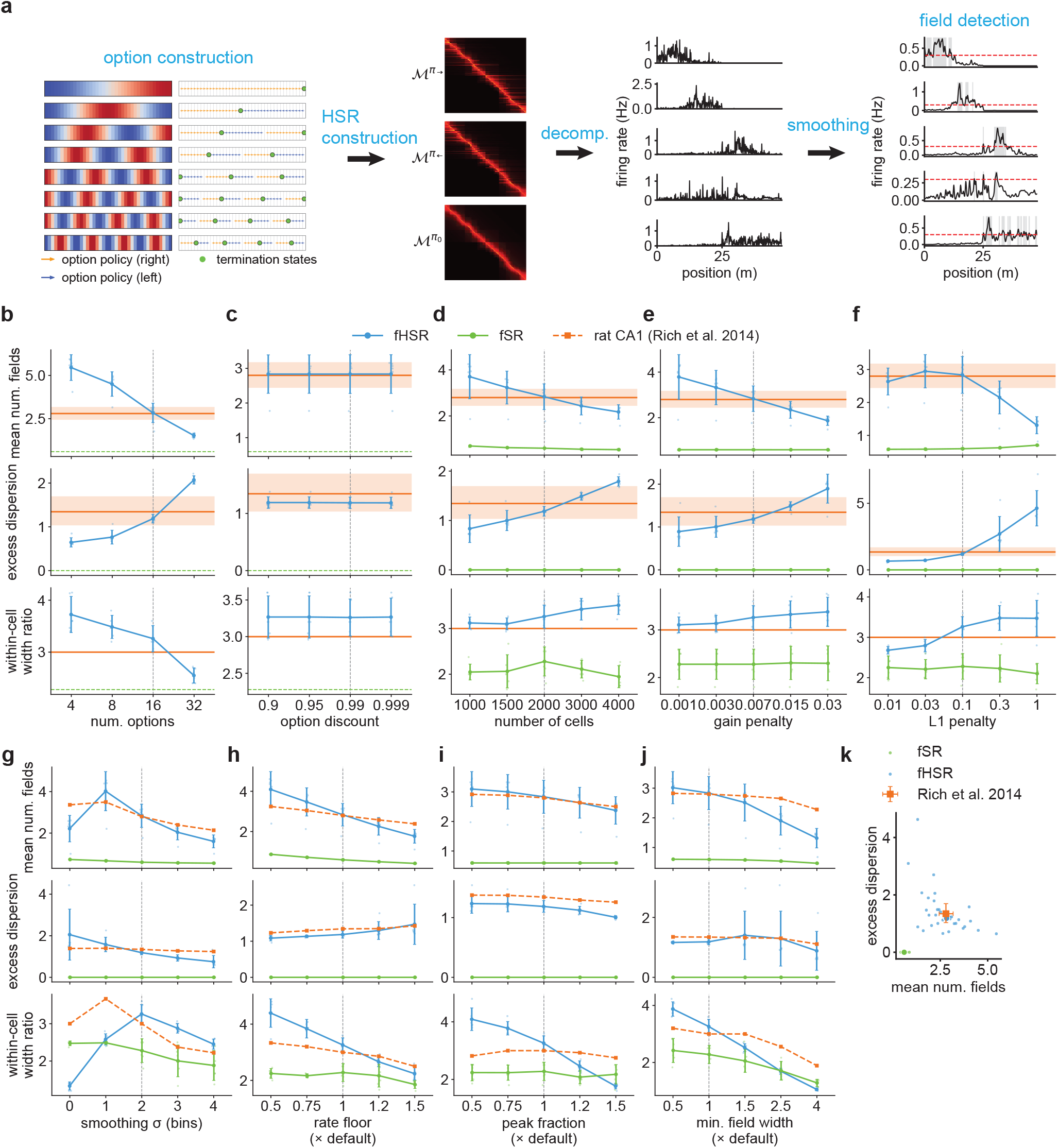
Place-field statistics of the factorised HSR (fHSR) are robust to model, decomposition and field-detection parameters. **a**. Pipeline for simulating fHSR place cells and detecting their place fields on linear tracks (from left to right). Eigenoptions are constructed from the eigenvectors of the random-walk SR (left; cf. Extended Data Figure 2); HSRs are computed under rightward 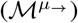, leftward 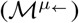 and random-walk 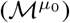 high-level policies; the quasi-NMF decomposition (decomp.) of the task-dependent HSRs yields the fHSR, whose columns are treated as place-cell spatial tuning curves; after smoothing, fields are detected as supra-threshold regions above a minimum field width (red dashed lines, threshold; grey, fields). **b-f**. Mean number of fields per cell (top), excess dispersion of the field-count distribution (middle) and median within-cell field-width ratio (bottom) for simulated place cells on the 48-m track, as a function of the number of options (b) and the option discount *γ*_o_ (c), and of the number of cells *M* (d), the gain penalty *γ*_*a*_ (e) and the *L*_1_ penalty *λ* (f) of the quasi-NMF decomposition, for the fHSR (blue) and the factorised SR (fSR; green). Orange lines and bands, rat CA1 data (mean ± s.e.m.). Dashed vertical lines, default values. **g-j**. Same as b-f, as a function of field-detection parameters, applied identically to model and data: the width of the Gaussian smoothing kernel (g; in spatial bins) and, relative to their defaults, the rate floor (h) and peak fraction (i) of the detection threshold and the minimum field width (j). Orange dashed lines, rat CA1 data re-analysed with the same parameters. **k**. Excess dispersion against mean number of fields per cell across all parameter settings in b-j, for the fHSR (blue), the fSR (green) and rat CA1 data (orange). Error bars, mean ± s.e.m.

**Extended Data Figure 10:**
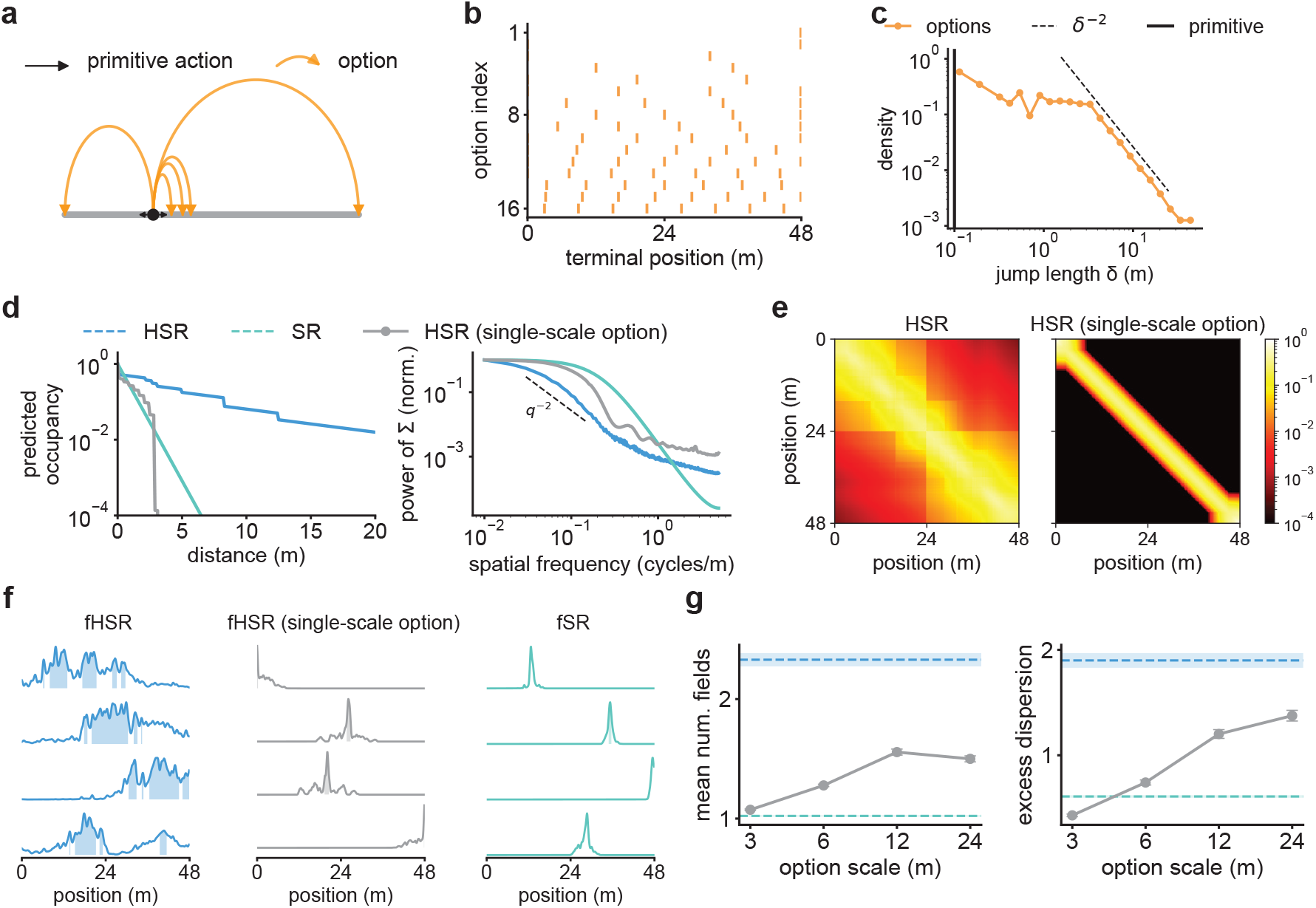
A multi-scale option hierarchy underlies multi-field, multi-scale tuning in the factorised HSR (fHSR). **a**. Schematics of the option-induced jump kernel (Supplementary Information, Section 2.3) on a linear track: primitive actions move the agent by one state (black), whereas options carry it to terminal states at a range of distances (orange). **b**. Terminal positions of the 16 eigenoptions on the 48-m track, ordered by option index. **c**. Distribution of displacement lengths *δ* produced by uniform selection among the options (orange), compared with the theoretically derived inverse-quadratic decay (dashed line; Supplementary Information, Section 2.3). Primitive actions produce unit displacements. **d**. Left, predicted occupancy as a function of distance from the current state for the spatial propagators induced by the HSR (blue), the SR (teal) and an HSR constructed from options of a single scale (grey). Right, power spectrum of the predictive-similarity matrix **Σ** (Equation 10; Methods) and theoretical prediction of the PSD scaling (dashed line; Supplementary Information, Section 4). **e**. HSRs constructed from multi-scale (left) and single-scale (right) option sets on the 48-m track (logarithmic colour scale). **f**. Example tuning curves of the fHSR (left), the fHSR constructed from single-scale options (middle) and the factorised SR (fSR; right). Shading, detected fields. **g**. Mean number of fields per cell (left) and excess dispersion (right) of fHSRs constructed from single-scale options as a function of the option scale (grey; mean ± s.e.m. over 10 random seeds), compared with the fHSR (blue dashed line) and the fSR (teal dashed line).

**Extended Data Figure 11:**
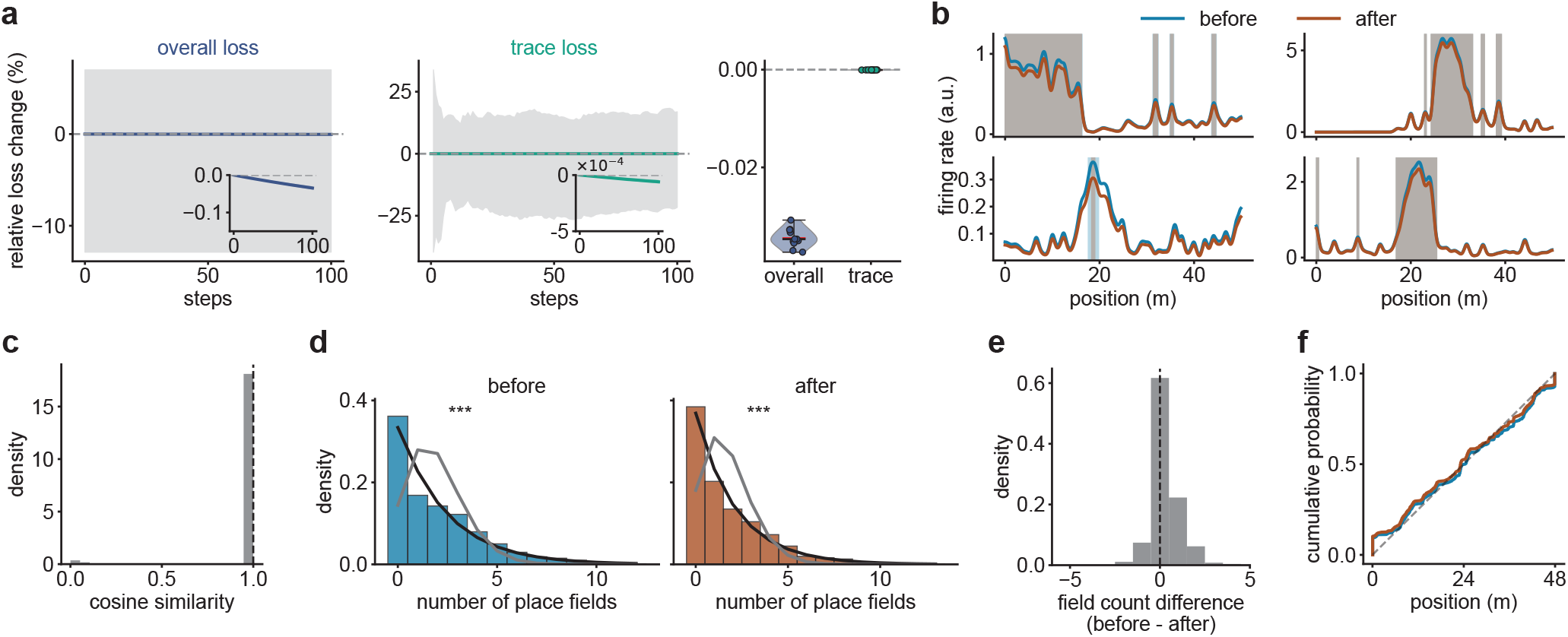
The factorised HSR (fHSR) and its field statistics are robust to the quasi-NMF optimisation. Place cells are extracted from the HSR of the 48-m track by initialisation with sparse symmetric NMF (Methods; Equation 12; Supplementary Information, Section 3.2.3), followed by quasi-NMF optimisation (Methods; Equation 11). To assess how much optimising with respect to the full decomposition objective beyond the heuristic initialisation altered the solution, we compared the basis at initialisation (before) with the basis after the full optimisation until convergence (after). **a**. Relative change (%) in the overall reconstruction loss (left) and in the trace loss (middle) over 100 refinement steps (insets, magnified scale), and the distribution of final relative changes (right). Lines and shading, mean and 95% confidence interval over 10 random seeds. Both losses change by a negligible fraction, indicating that the initialisation already lies close to the optimum. **b**. Tuning curves of four example cells before and after full optimisation. Shading, identified fields. **c**. Distribution of cosine similarities between matched basis vectors before and after full optimisation. Dashed line, median. **d**. Distributions of the number of fields per cell before and after full optimisation, with gamma-Poisson and equal-Poisson fits (likelihood ratio tests; before: *χ*^2^(1) = 1,089.74, *p* = 5.6 × 10^*−*239^; after: *χ*^2^(1) = 1,013.75, *p* = 1.8 × 10^*−*222^). **e**. Per-cell change in the number of fields (before minus after). Dashed line, median. **f**. Cumulative distributions of field locations before and after full optimisation. Dashed line, uniform distribution. Starting from the symmetric NMF of the predictive similarity, the full optimisation leaves the basis, single-cell tuning and population field statistics effectively unchanged, so the place-cell statistics reflect the structure of the HSR rather than details of the optimisation. \*\*\**p <* 0.001.

**Extended Data Figure 12:**
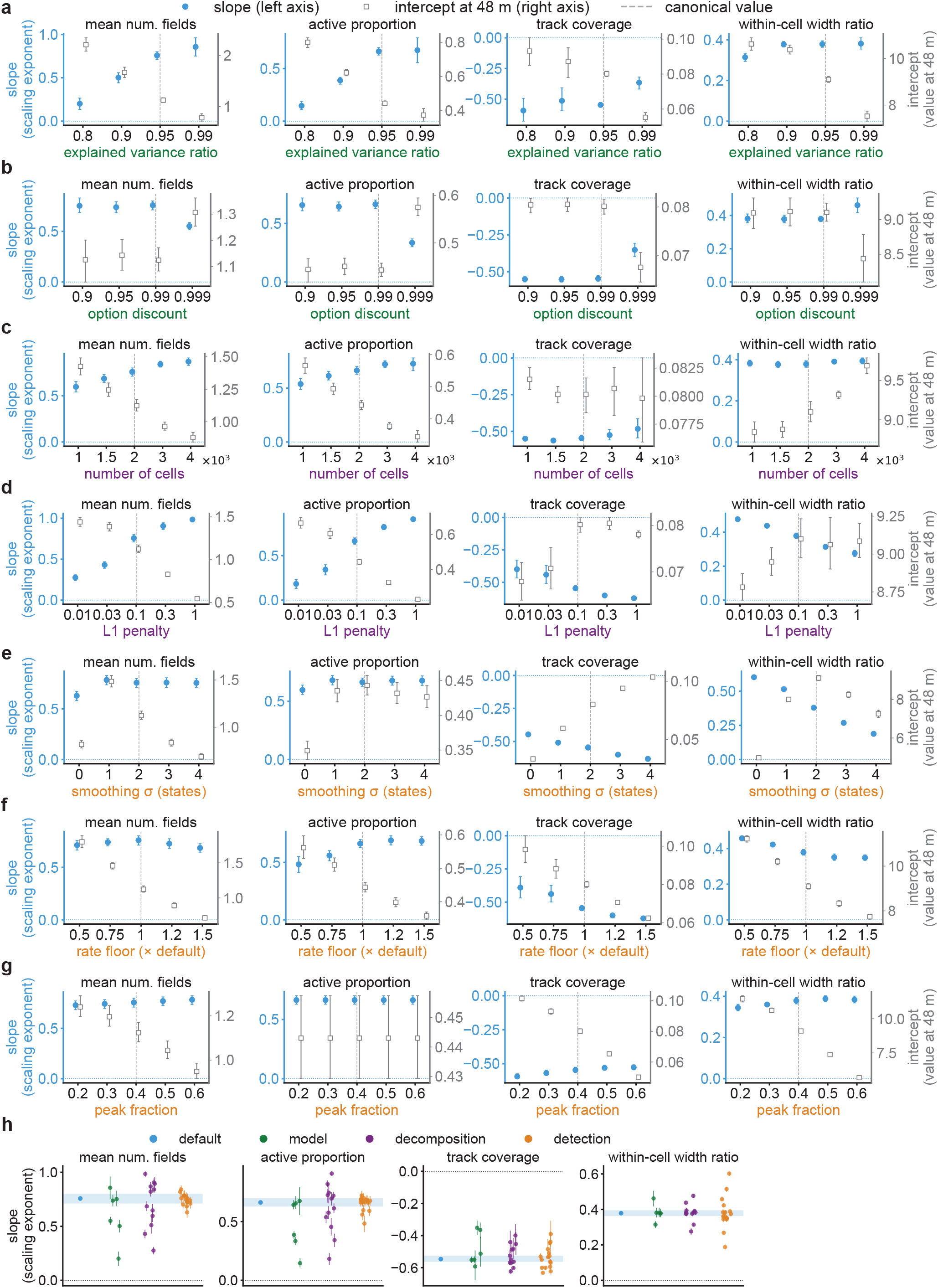
The scaling of place-field statistics with track length is robust to model, decomposition and detection parameters. **a-g**. Scaling exponent (slope of the logarithm of each statistic against the logarithm of track length; filled blue circles, left axes) and value at 48 m (intercept; open squares, right axes) of (from left to right) the mean number of fields per cell, the proportion of active cells, track coverage and the within-cell field-width ratio (cf. Figure 4), as a function of the parameters of option construction (a, option-selection threshold *θ*_*f*_, the explained-variance ratio of Equation 6; b, option discount *γ*_o_), of the quasi-NMF decomposition (c, number of cells *M* ; d, *L*_1_ penalty *λ*) and of field detection (e, width of the Gaussian smoothing kernel; f, rate floor; g, peak fraction of the detection threshold). Dashed vertical lines, default values used in the main text. Error bars, 1 s.e.m. **h**. Scaling exponents across all parameter settings, grouped by parameter class (model, green; decomposition, purple; detection, orange); blue, default setting; band, s.e.m. over 10 random seeds.

**Extended Data Figure 13:**
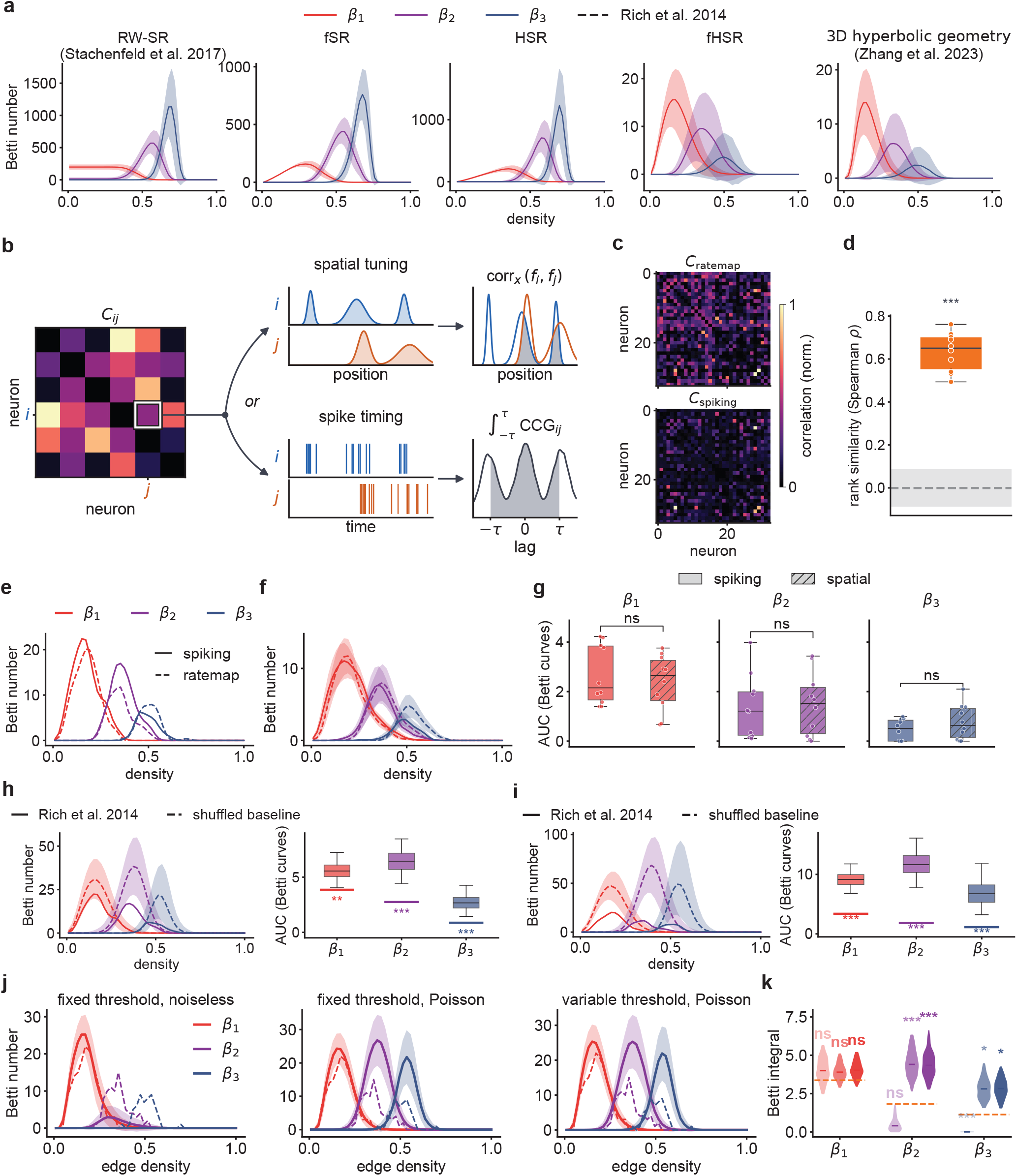
The topological correspondence between the factorised HSR (fHSR) and rat CA1 is specific to the fHSR and robust to the measure of co-activity. **a**. Betti curves of (from left to right) RW-SR columns ^3^, the factorised SR (fSR), raw HSR columns (HSR), the fHSR and the three-dimensional hyperbolic reference ^17^ (mean ± s.d. over 300 random seeds; cf. Figure 5); dashed lines, rat CA1 place cells on the 48-m track ^11^. Note the substantially different scales along the vertical axes. **b**. Two measures of pairwise co-activity: the correlation between spatial tuning curves (top) and the integral of the spike-train cross-correlogram (CCG) within a lag window of ±*τ* (bottom). **c**. Example correlation matrices of the same rat CA1 cells computed from rate maps (top) and spike trains (bottom). **d**. Rank similarity (Spearman *ρ*) between rate-map and spike-train correlation matrices computed from rat CA1 neural activity across 8 sessions on the 48-m linear track (5 rats; points); grey band, shuffled null (10,000 shuffles; 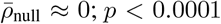. **e**. Betti curves computed from spike-train (solid) and rate-map (dashed) correlations for an example session. **f**. Same as e, mean ±s.d. across all 8 sessions. **g**. Integrated Betti curves from spike-train and rate-map correlations (two-sided Mann–Whitney *U* tests; *β*_1_: *U* = 59.0, *p* = 0.5205; *β*_2_: *U* = 489.5, *p* = 1.0000; *β*_3_: *U* = 37.0, *p* = 0.3383). **h**. Left, Betti curves of rat CA1 place cells computed from the spike-train CCG (solid) and of a shuffled baseline (dashed, mean ± s.d.; fields relocated at random, preserving their number and widths; 1,000 shuffles; Methods). Right, integrated Betti curves of shuffles and data (horizontal lines; *β*_1_: *p* = 0.007, *β*_2_: *p <* 0.001, *β*_3_: *p <* 0.001). **i**. Same as h, for Betti curves computed from tuning-curve correlations (*β*_1_: *p <* 0.001, *β*_2_: *p <* 0.001, *β*_3_: *p <* 0.001). **j**. Betti curves (mean ± s.e. over 300 seeds) of populations whose spatial tuning curves are random functions, drawn from Gaussian processes and rectified at a threshold ^16^ (Methods): (left to right) fixed threshold, with tuning curves used in their canonical, noiseless form; fixed threshold, with tuning curves re-estimated from Poisson spike counts; and variable threshold, with Poisson re-estimation. Dashed lines, rat CA1 place cells on the 48-m track. **k**. Same as h and i, for the integrated Betti curves of the three model configurations in j (left to right within each Betti number); orange dashed lines, rat CA1 data (two-sided permutation test with 300 seeds; fixed threshold + noiseless *β*_1_: *p* = 0.4267, *β*_2_: *p* = 0.0533, *β*_3_: *p <* 0.0033; fixed threshold + Poisson *β*_1_: *p* = 0.2667, *β*_2_: *p <* 0.0033, *β*_3_: *p* = 0.0133; variable threshold + Poisson *β*_1_: *p* = 0.2400, *β*_2_: *p <* 0.0033, *β*_3_: *p* = 0.0133). \**p <* 0.05; \*\**p <* 0.01; \*\*\**p <* 0.001 (h,i) or *p <* 0.0033 (k); ns, not significant.

**Extended Data Figure 14:**
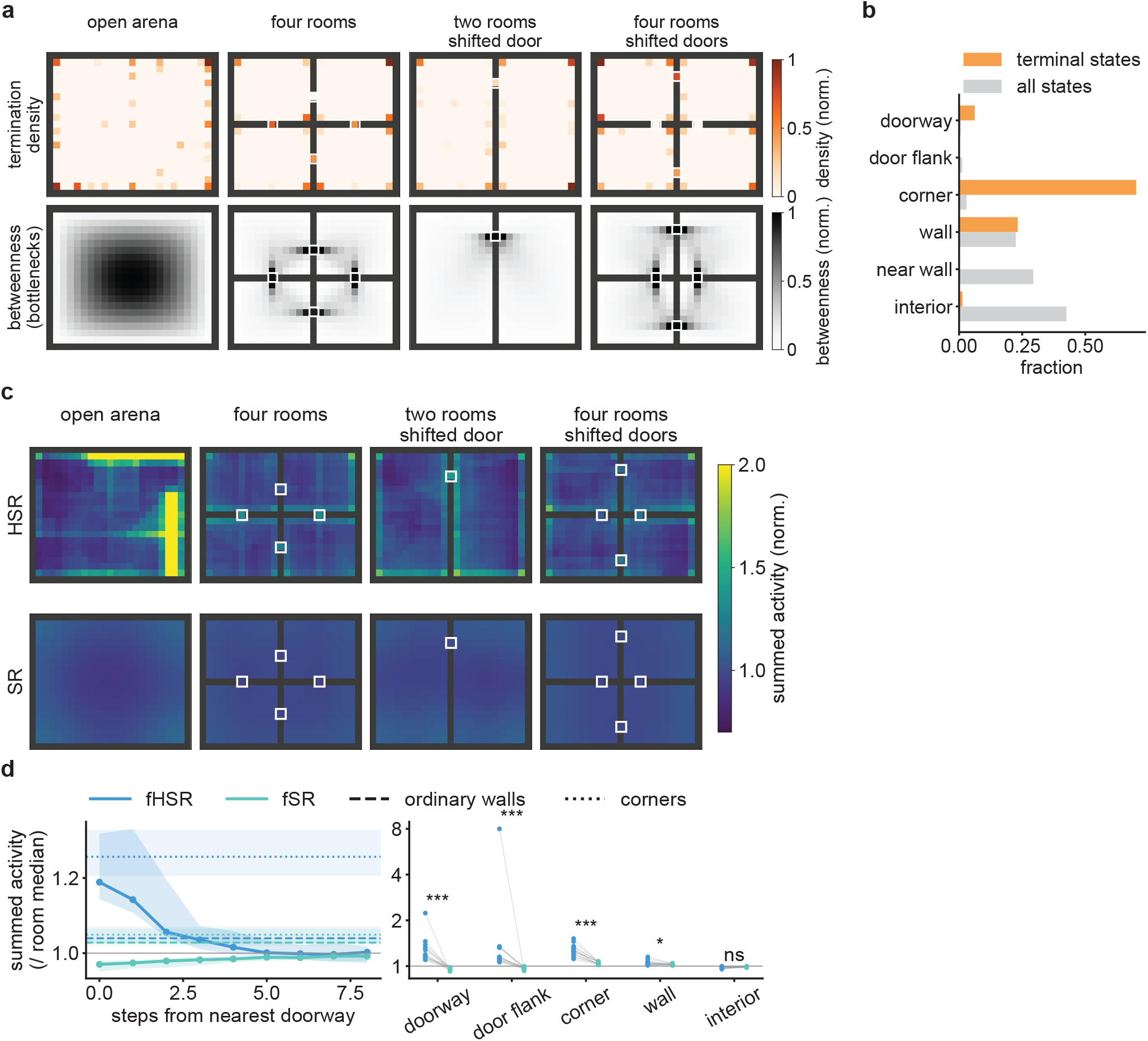
The factorised HSR (fHSR) predicts over-representation of option terminal states, which coincide with topologically significant states in structured environments. **a**. Top, density of option terminal states in an open arena, the four-room environment, a two-room environment and a four-room environment with shifted doors. Bottom, betweenness centrality (normalised), which identifies bottleneck states (doorways). **b**. Fraction of option terminal states (orange) and of all states (grey) in each state class (categorised given betweenness centrality), pooled over environments. **c**. Population summed activity (normalised) of the fHSR (top) and the factorised SR (fSR; bottom). White squares, doorways. **d**. Left, summed activity (normalised to the median of the room) as a function of the distance from the nearest doorway for the fHSR (blue) and the fSR (teal); dashed and dotted lines, mean at ordinary walls and corners (mean ± s.d.). Right, summed activity at states by location category; grey lines connect matched states (two-sided Wilcoxon signed-rank tests, fHSR versus fSR; doorway, *W* = 0.0, *p* = 0.0001; door flank, *W* = 0.0, *p* = 0.0001; corner, *W* = 0.0, *p* = 0.0001; wall, *W* = 15.0, *p* = 0.017; interior, *W* = 25.0, *p* = 0.091; *n* = 15 random seeds). \**p <* 0.05; \*\*\**p <* 0.001; ns, not significant.

**Extended Data Figure 15:**
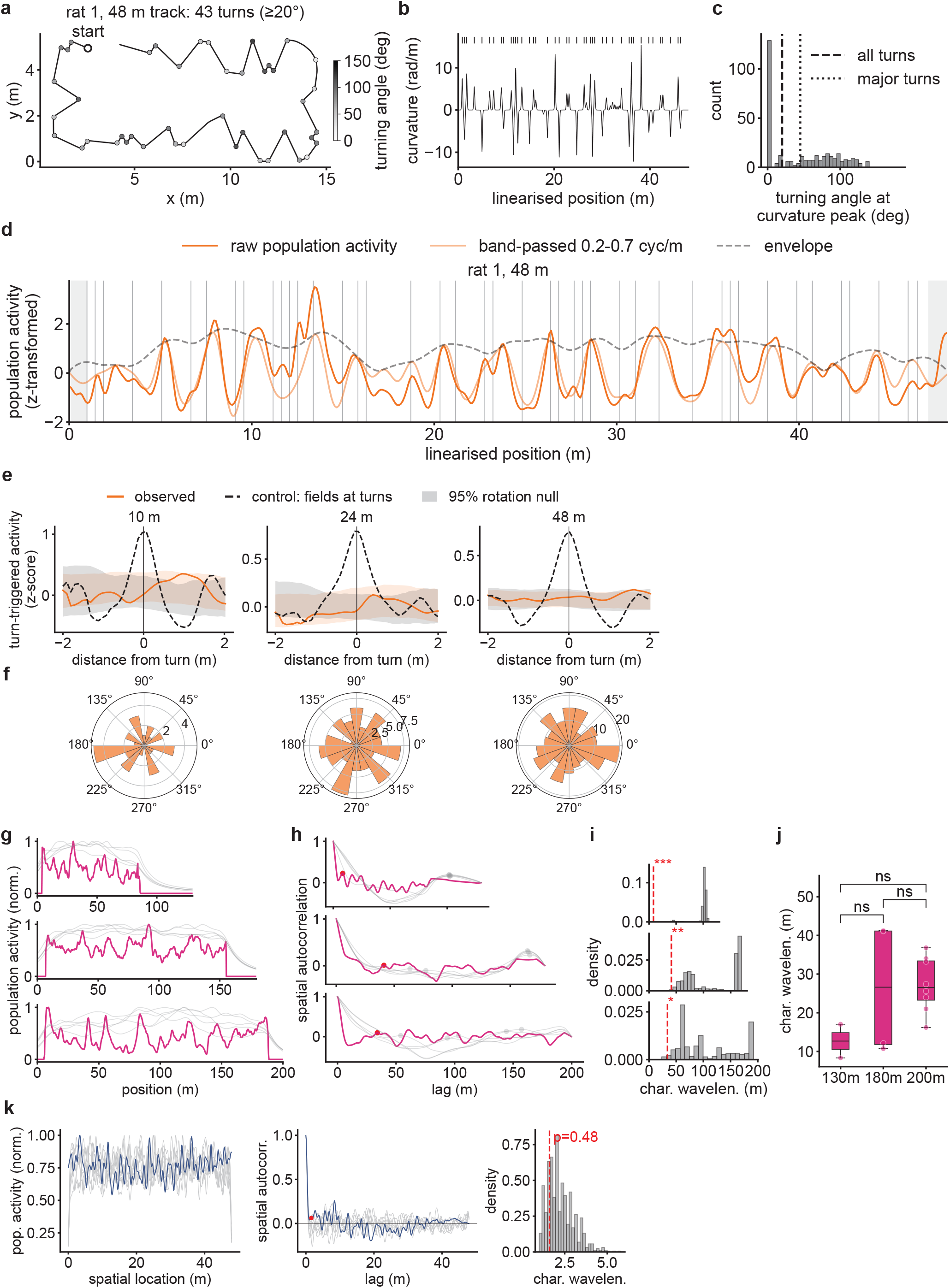
The spatial periodicity of population activity is not locked to track geometry, generalises to bats and is absent from random-function populations. **a**. Layout of the 48-m track of an example rat, with local peaks in the curvature along the track (grey scale, turning angle; 43 turns). **b**. Curvature along the linearised track; top ticks, detected turns. **c**. Distribution of turning angles at curvature peaks (over all 8 sessions). Dashed and dotted lines, thresholds for all turns (turn angle *>* 20^*°*^) and for major turns (turn angle *>* 45^*°*^). **d**. Z-transformed summed population activity of rat CA1 place cells within the example session as a function of the linearised position: raw (solid), band-passed at 0.2–0.7 cycles m^*−*1^ (thin) and its envelope (dashed). Grey vertical lines, turns; grey shading, excluded track ends. **e**. Turn-triggered average of band-passed population activity (orange) on the 10-, 24- and 48-m tracks, compared with the 95% interval of the null distribution obtained by randomly shifting turn positions (grey; mean ± s.d.; 1,000 shuffles) and with a positive control in which all fields are shifted to their nearest turns (dashed; observed, 10 m: *p* = 0.2959, 24 m: *p* = 0.4908, 48 m: *p* = 0.6527; positive control, 10 m: *p* = 0.001, 24 m: *p* = 0.001, 48 m: *p* = 0.001). Summed population activity is not locked to track turns at any of the tested track lengths. **f**. Distributions of the phase of the band-passed activity at turns (Rayleigh tests for uniformity; 10 m: *z* = 0.7029, *p* = 0.4988; 24 m: *z* = 0.3426, *p* = 0.7099; 48 m: *z* = 0.1041, *p* = 0.9012; mean resultant length 10 m: *ρ* = 0.1506, 24 m: *ρ* = 0.0635, 48 m: *ρ* = 0.0219). **g**. Normalised summed population activity of bat CA1 place cells ^14^ along 130-, 180- and 200-m tracks (magenta), with shuffles (grey). **h**. Spatial autocorrelations of the profiles in g. Dots, first non-zero peak. **i**. Null distributions of the characteristic wavelength over 1,000 shuffles; red dashed lines, data (130 m: *p <* 0.001; 180 m: *p* = 0.003; 200 m: *p* = 0.017). **j**. Characteristic wavelengths of 14 bats on the three track lengths (two-sided Mann–Whitney *U* tests; 130 versus 180 m: *U* = 2.0, *p* = 0.5333; 130 versus 200 m: *U* = 1.0, *p* = 0.0889; 180 versus 200 m: *U* = 16.0, *p* = 1.0000; LMM, *β* = 0.2017, *p* = 0.1104). **k**. Same as g-i, for an example population of the random-function model ^16^ on the 48-m track (blue; Extended Data Figure 13j; Methods), with shuffles (grey) and the null distribution of the characteristic wavelength (1000 shuffles; red dashed line, model; *p* = 0.48). \**p <* 0.05; \*\**p <* 0.01; \*\*\**p <* 0.001; ns, not significant.

## Supplementary Information

### 1 Notations

Throughout the note, we will use the following notations.

**Table 1:**
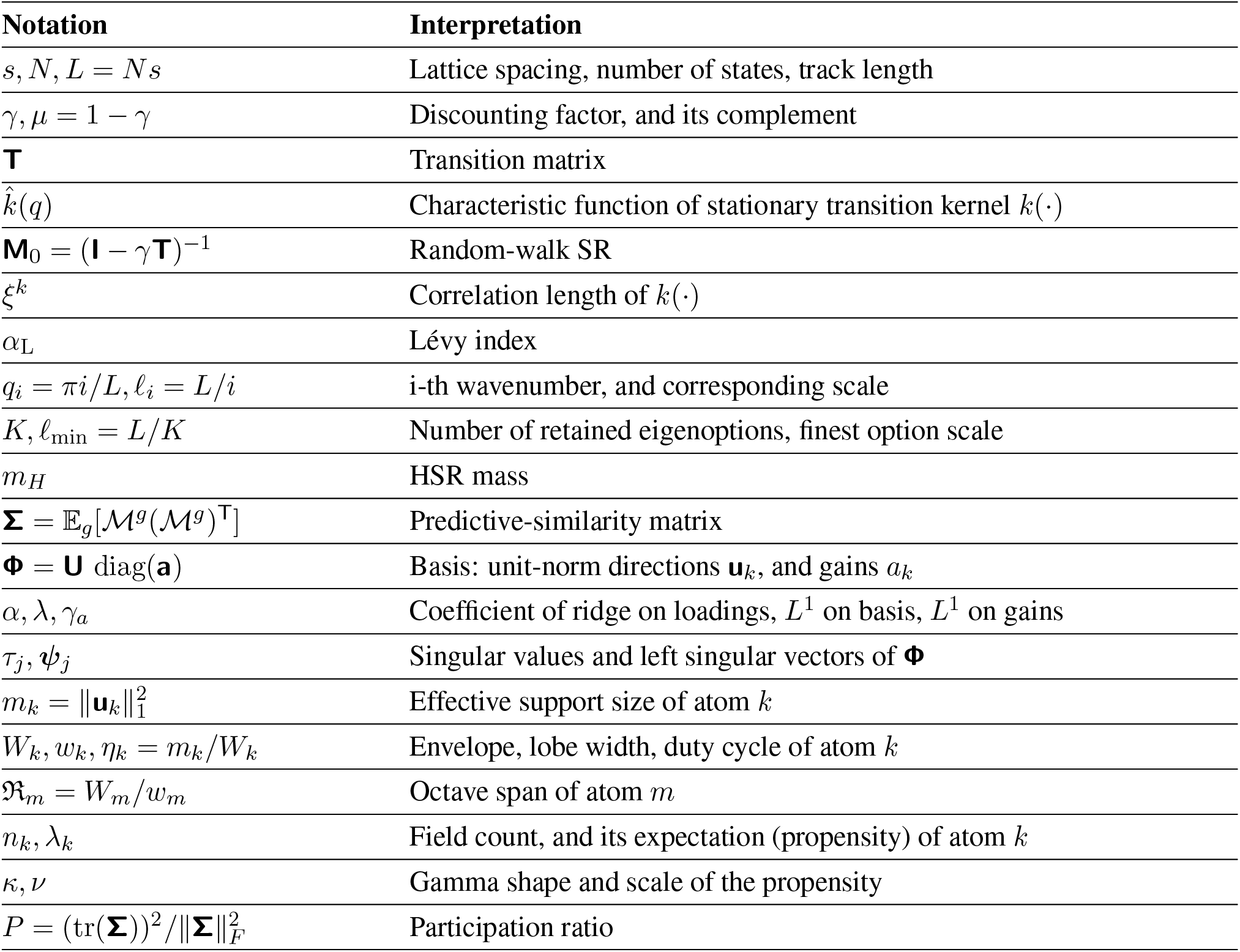
Notation used throughout the text.

| Notation | Interpretation |
|---|---|
| s, N, L = Ns | Lattice spacing, number of states, track length |
| γ, μ = 1 − γ | Discounting factor, and its complement |
| **T** | Transition matrix |
| k̂(q) | Characteristic function of stationary transition kernel k(·) |
| **M**<sub>0</sub> = (**I** − γ**T**)<sup>−1</sup> | Random-walk SR |
| ξ<sup>k</sup> | Correlation length of k(·) |
| α<sub>L</sub> | Lévy index |
| q<sub>i</sub> = πi/L, ℓ<sub>i</sub> = L/i | i-th wavenumber, and corresponding scale |
| K, ℓ<sub>min</sub> = L/K | Number of retained eigenoptions, finest option scale |
| m<sub>H</sub> | HSR mass |
| **Σ** = 𝔼<sub>g</sub>[ℳ<sup>g</sup>(ℳ<sup>g</sup>)<sup>**T**</sup>] | Predictive-similarity matrix |
| **Φ** = **U** diag(**a**) | Basis: unit-norm directions **u**<sub>k</sub>, and gains a<sub>k</sub> |
| α, λ, γ<sub>a</sub> | Coefficient of ridge on loadings, L<sup>1</sup> on basis, L<sup>1</sup> on gains |
| τ<sub>j</sub>, **ψ**<sub>j</sub> | Singular values and left singular vectors of **Φ** |
| m<sub>k</sub> = ‖**u**<sub>k</sub>‖<sub>1</sub><sup>2</sup> | Effective support size of atom k |
| W<sub>k</sub>, w<sub>k</sub>, η<sub>k</sub> = m<sub>k</sub>/W<sub>k</sub> | Envelope, lobe width, duty cycle of atom k |
| ℜ<sub>m</sub> = W<sub>m</sub>/w<sub>m</sub> | Octave span of atom m |
| n<sub>k</sub>, λ<sub>k</sub> | Field count, and its expectation (propensity) of atom k |
| κ, ν | Gamma shape and scale of the propensity |
| P = (tr(**Σ**))<sup>2</sup>/‖**Σ**‖<sub>F</sub><sup>2</sup> | Participation ratio |

### 2 SR and HSR

#### 2.1 Preliminaries

We consider operating on the one-dimensional lattice with *N* sites and spacings *s*, with *x*_*n*_ = *ns* for *n* = 0, …, *N* − 1.

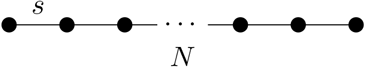

We consider the symmetric nearest-neighbour random walk with reflecting boundaries (movements into a boundary leave the agent in place).

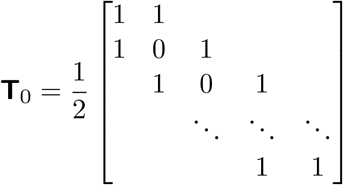

Note that **T**_0_ is symmetric, row-stochastic, and acts on observables via (**T**_0_*f* )(*n*) = E [*f* (*s*_*t*+1_)|*s*_*t*_ = *n*].

The SR is defined as the expected future total (discounted) occupancy ^1^.

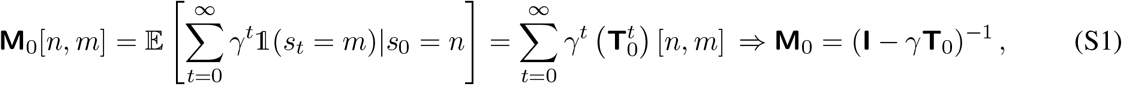

Note that due to **T**_0_ is stochastic, and the spectral radius of *γ***T**_0_ equals *γ <* 1, hence the above resolvent expression of the SR always exists.

#### 2.2 SR as an exponentially decaying diffusion operator

We firstly consider the bulk case where states are away from the reflecting boundaries 1≪ *n* ≪ *N*, then the primitive random walk is spatially homogeneous. Specifically, suppose the current position is *x*, the next position can be expressed as *x*^′^ = *x* + Δ with state-independent increment Δ. Note that in the bulk case, the symmetric random-walk transition operator is translation invariant, hence plane waves diagonalise the transition matrix (the transition operator commutes with shift operation, and is hence a convolution operator).

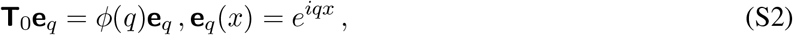

In fact, the plane waves, {**e**_*q*_}, are eigenfunctions for any spatially homogeneous random walk. For the primitive one-step random walk, 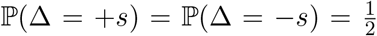, hence 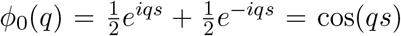 are the symbols (Fourier transform) of the transition operator **T**_0_.

Now given the symbol of **T**_0_, **I** − *γ***T**_0_ has symbol 1 − *γ* cos(*qs*), and is non-vanishing since |*γ* cos(*qs*)| ≤ *γ <* 1. Hence the symbol of the random-walk SR is.

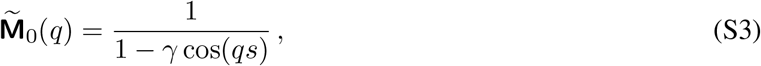

We then apply inverse Fourier transform to 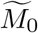 to retrieve the corresponding kernel function for **M**_0_ in the original lattice space.

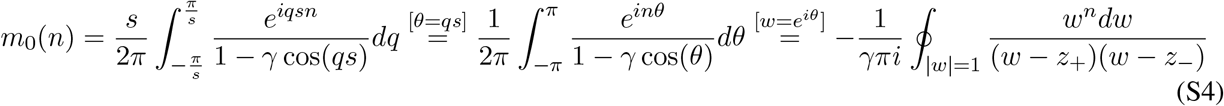

where 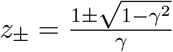 . Since *z*+ *z*_−_ = 1, exactly one root lies within the unit circle. Suppose *z* ≡ *z*_−_ *<* 1. For *n* ≥ 0, the residue at *w* = *z* gives us the value of the integral.

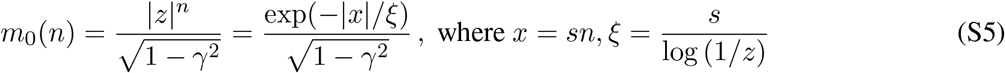

Leveraging *z*_+_*z*_−_ = 1, we have the following simplified expression.

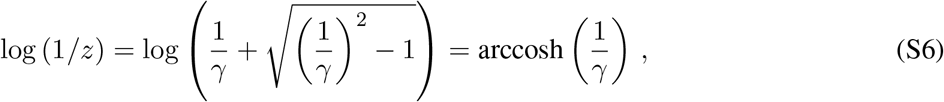

Leveraging first-order Taylor expansion when *γ* is close to 1, we have the following.

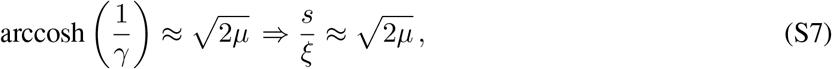

Hence in the spatial bulk, the SR induces an explicit real-space spatial propagator, which decays exponentially with distance.

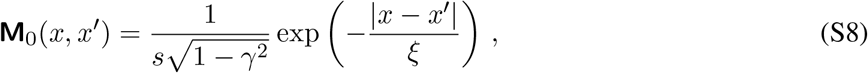

and is fully parameterised by a single parameter, the discounting factor *γ* (Equation S6). Note that Equation S8 specifies a density per unit space, and the actual lattice values are *m*_0_(*n*) = *s***M**_0_(*x*_*n*_, 0) (Equation S5).

##### 2.2.1 Alternative approximation scheme of SR’s correlation lengthscale

For small wavenumbers (*qs* ≪ 1), we expand the cosine term (Equation S3) using its Taylor series around 0.

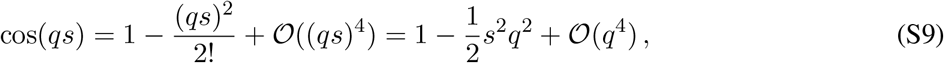

Substitute the denominator approximation into 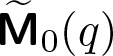.

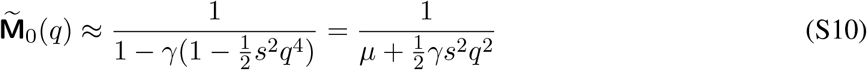

We then match the above approximation to the classical Lorentzian form, 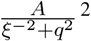 (see Note 2.1), and have the following approximation over the correlation lengthscale.

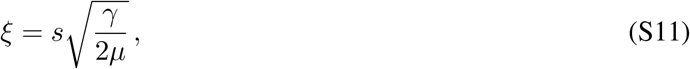

The above approximation accurately matches the exact correlation lengthscale (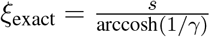; Equation S5), with ∼ 0.9% error for *γ* = 0.9 and ∼ 0.04% error for *γ* = 0.995.

**Note (Lorentzian form) 2.1**

The SR is defined by the matrix inverse (S1), which in position space is equivalent to

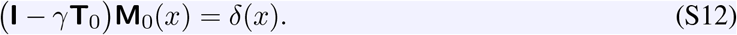

For a single-step symmetric random walk on a grid with spacing *s*, the transition operator acts on **M**_0_(*x*) as follows: *γ γ*

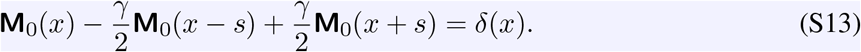

We then subtract and add *γ***M**_0_(*x*) on the left-hand side to group the terms:

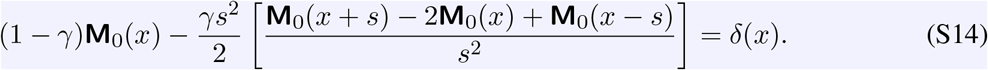

Note the term inside the bracket is the exact discrete second derivative (the discrete Laplacian operator).

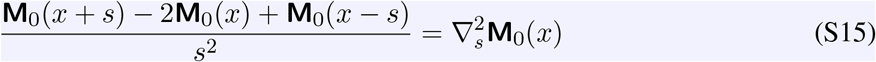

In the continuous limit (*s* → 0), the discrete operator 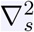 converges to the continuous spatial derivation 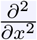. The spatial difference equation hence becomes a continuous differential equation.

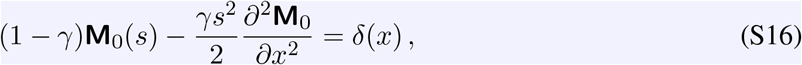

Defining 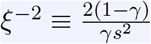, the equation becomes the standard Poisson equation.

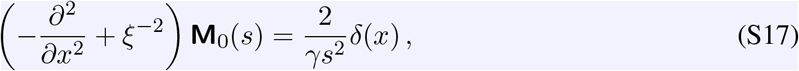

Now taking Fourier transform of this continuous differential equation, and leveraging 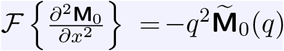.

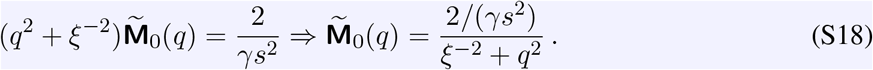

Note that it is also clear to see from Equation S17 that the SR is the lattice Green’s function of a massive diffusion operator, (−∇^2^ + *m*^2^)**M**(·) = *δ*(·), with the mass term inversely proportional to the correlation lengthscale 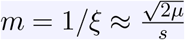, which is solely dependent on the discounting factor, *γ*.

#### 2.3 HSR induces Cauchy-flight spatial propagator

On the one-dimensional lattice with reflecting boundaries, the eigenvectors of the transition matrix (and for the SR) are standing waves, 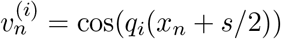, where *q*_*i*_ = *πi/L* and *x* = *ns*. This can be easily validated.

- Interior (1 ≤ *n* ≤ *N* − 2).

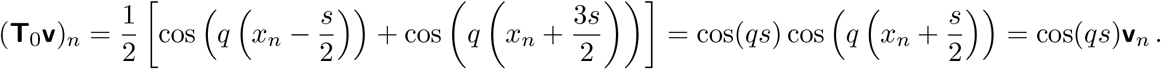
- Left boundary (n = 0).

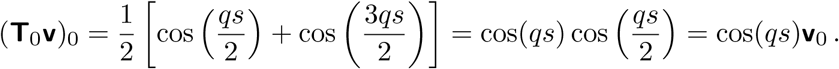
- Right boundary (n = N-1).

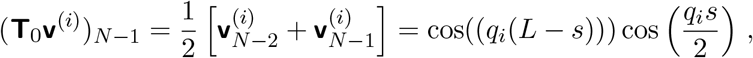 Note that cos((*q*_*i*_(*L*−*s*))) = cos(*πi*−*q*_*i*_*s*) = (−1)^*i*^ cos(*q*_*i*_*s*), hence 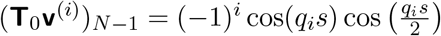. Now evaluating the RHS by firstly expanding 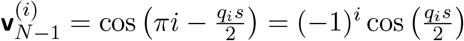. We hence have the equivalence.

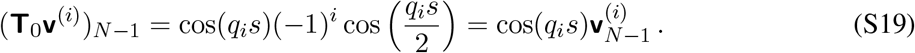

Note that the half-integer offset is the discrete Neumann condition, and is specifically generated by the reflecting boundary conditions.

Extrema of **v**^(*i*)^ sit at 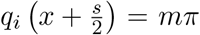, i.e., 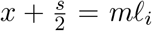 where *m* is arbitrary integer and 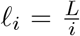. Hence the terminal states of option *i* (the maxima of **v**^(*i*)^) are spaced 2*l*_*i*_ apart, partitioning the track into segments of length 2*l*_*i*_. Starting uniformly within a basin, the executed displacement |Δ| ∼*U* (0, *l*_*i*_). The eigenoption ladder is hence harmonic.

With uniform selection amongst *K* options, the density of |Δ| can be computed as following.

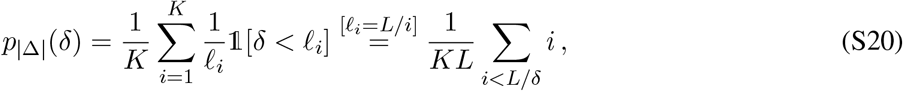

- For *δ > l*_min_, where *l*_min_ = *L/K*, the sum runs to *L/δ*.

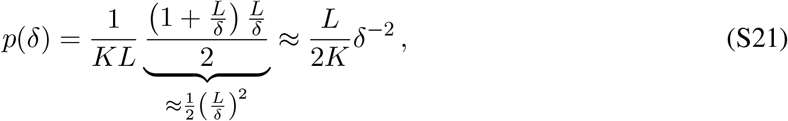

- For *δ < l*_min_, the sum runs over all *K* options.

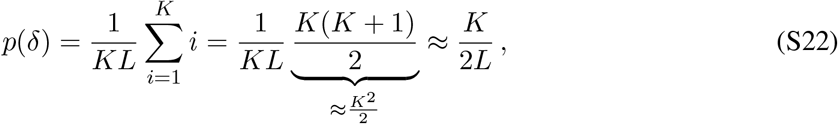

Note that *p*(*δ*) is a proper density function that integrates to 1.

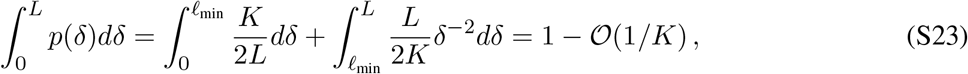

For symmetric spatial transitions on a line (Δ ∈ [−*L, L*]), we have that 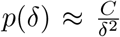 (Equation S21), where 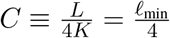.

It is also simple to check that 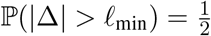, confirming that exactly 50% of all generated jump events are long-range transitions exceeding the option resolution *l*_min_.

In random-walk theory, heavy-tailed jump distributions satisfy the following condition.

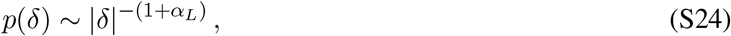

Matching the powers (Equation S21), we have that the Lévy index 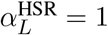, which corresponds to a Cauchy-flight Lévy process ^3^.

##### 2.3.1 Modulation of Lévy index through option selection

Instead of uniform selection of options, suppose we weight the probability of selecting the i-th option with probability *w*_*i*_ ∝ *i*^−*θ*^ for some *θ* ∈ (−1, 1).

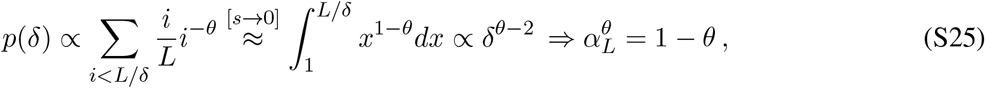

Hence *α*_*L*_ is a tunable knob, with three corresponding regimes.

- Uniform selection (*θ* = 0) ⇒ *α*_*L*_ = 1 (Cauchy flight).
- Upweighting coarse options (*θ >* 0) ⇒ *α*_*L*_ *<* 1 (more jumpy).
- Upweighting fine options (*θ <* 0) ⇒ *α*_*L*_ *>* 1 (less jumpy).

The Cauchy flight (*α*_*L*_ = 1) is a special regime. To see this, we firstly compute the mean displacement.

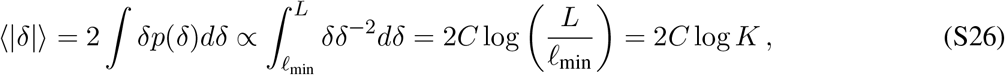

We then evaluate the contribution of ⟨*δ*⟩ within arbitrary spatial octave [*l*, 2*l*], ∀*l* ∈ (*l*_min_, *L/*2].

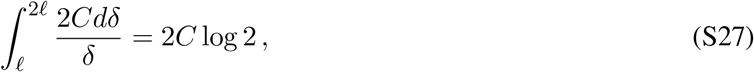

Hence, all spatial hierarchies (corresponding to all spatial octaves) contribute exactly 2*C* log 2 to the expected exploration distance, regardless of where the hierarchy sits. For *α*_*L*_ *<* 1, the mean is dominated by the coarse cutoff, and for *α*_*L*_ *>* 1 by the fine ones. Only at *α*_*L*_ = 1 the option ladder becomes scale-invariant, ensuring equal representation across all spatial scales without fine or coarse cutoff dominance.

##### 2.3.2 HSR is the Green’s function to the fractional Laplacian operator

Recall the resolvent formulation of the HSR (with respect to some high-level policy *µ*).

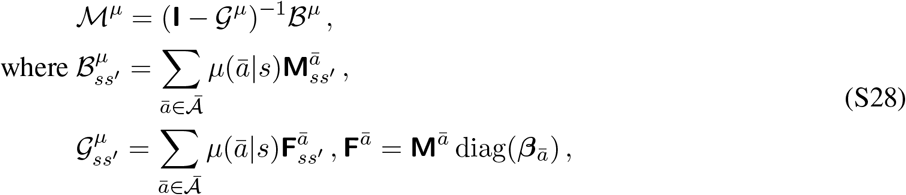

We again take the assumption that we are operating exclusively with options in our subsequent derivations. The *hierarchical continuation kernel* is equivalent to the discounted jump kernel (Equation S20). Specifically, for a jump of magnitude *δ*, the corresponding duration is *τ* = |*δ*| */s* lattice steps. We further assume, for simplicity, that the target lattice lies within the bulk of an infinite lattice, hence guaranteeing translation invariance.

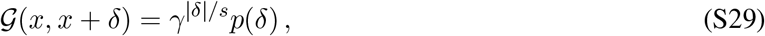

Leveraging the resolvent formulation of the HSR (Equation S28), we can express the Fourier transform of the HSR as following.

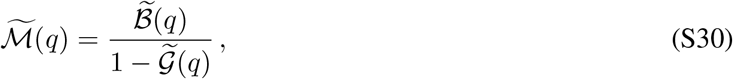

We focus on the denominator, which can be decomposed as following.

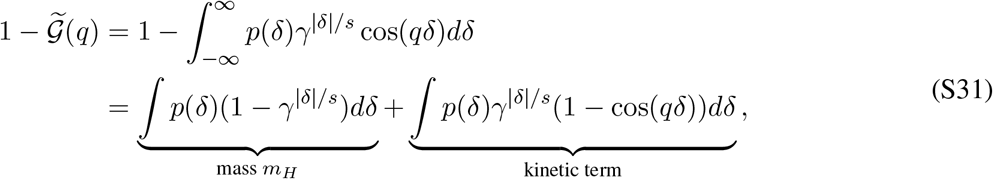

Note that the first equality holds since 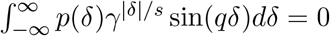, given that *p*(*δ*) is symmetric around 0.

For *γ* ∼ 1, *µ* ≪ 1, we leverage the following approximation of *γ*^*δ/s*^ to first order in *µ*(= 1 − *γ*)^1^.

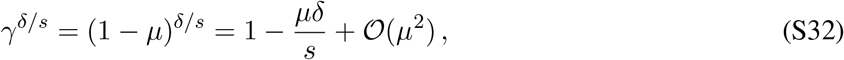

We then substitute this approximation into the mass term.

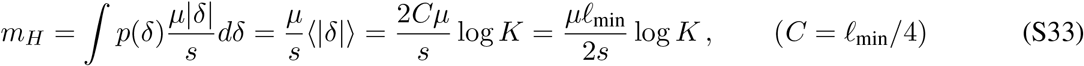

Given the derived *p*(*δ*) (Equation S21), we compute the kinetic term.

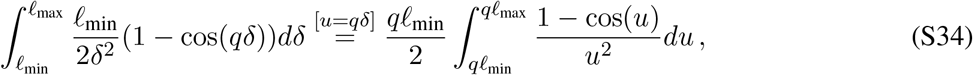

Assume 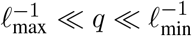, then the integral evaluates to 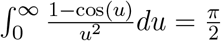. Hence the Fourier transform of the spatial propagator induced by the HSR takes the following expression.

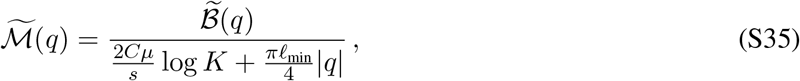

We then define 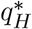 to be the “crossover” wavenumber, i.e., the critical spatial frequency at which the behaviour of the propagator transitions from kinetic-dominated to mass-dominated.

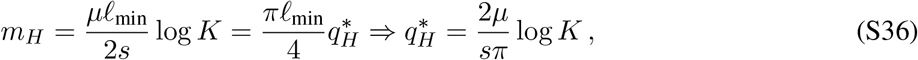

Hence, within the intermediate scaling regime, where 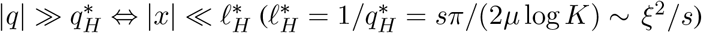, the kinetic term dominates.

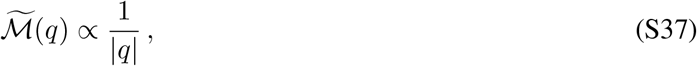

We can retrieve the HSR-induced spatial propagator through taking the inverse Fourier transform.

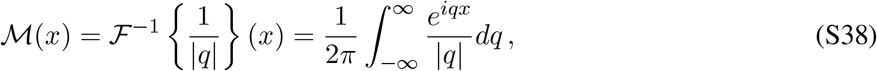

This integral cannot be directly evaluated, but its derivative can be analytically computed.

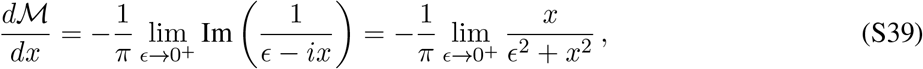

We hence have that 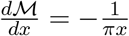, which we can integrate to derive the spatial propagator.

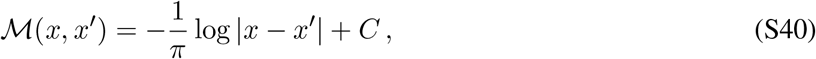

Hence the HSR yields logarithmic decay in real space, instead of exponential.

Note that so far we have only considered the intermediate scaling regime where 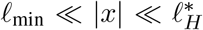. For the |*x*| ≤ *l*_min_ case, the transition kernel is locally flat since it is below the finest option scale. For the 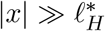 regime, because large spatial distances probe the small wavenumber limit 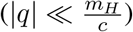, we can then expand the denominator as a geometric series around *q* = 0.

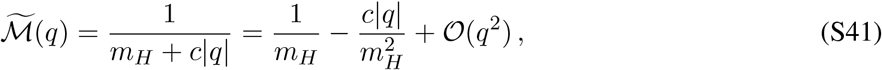

Taking the inverse Fourier transform, the first constant terms evaluates to a delta function, and the transition kernel is dominated by the linear term.

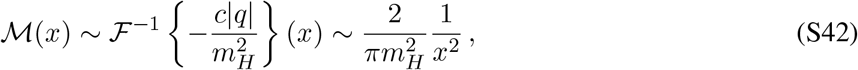

Again in contrast with the SR, the transition kernel is this small-wavenumber limit regime is polynomial in *q*^2^, instead of exponential decay.

Importantly, |*q*| is the Fourier transform of the fractional Laplacian operator of order 1, |∇| = (−Δ)^1*/*2 4^, and given Equation S37, we have that 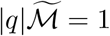. By taking the inverse Fourier transform, we have |∇|M = *δ*, and the HSR is the Green’s function of |∇|.

##### 2.3.3 Lévy index of the standard SR

The Fourier transform of the jump distribution corresponding to the symmetric single-step random-walk transition operator is simply its symbols.

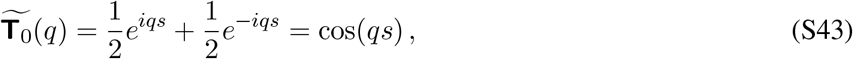

From random-walk theory, the fraction of spatial phase lost per step is governed by the kinetic term, 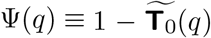, and 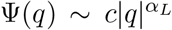 for any isotropic spatial transition, where *α*_*L*_ ∈ (0, 2] is the Lévy index. Leveraging the similar approximation scheme (Equation S9), we have that 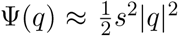, hence the Levy index for standard SR is *α*_*L*_ = 2.

#### 2.4 Summary comparison between SR and HSR

Below we summarise and contrast the key microscopic and spectral properties of both representations.

**Table 2:**
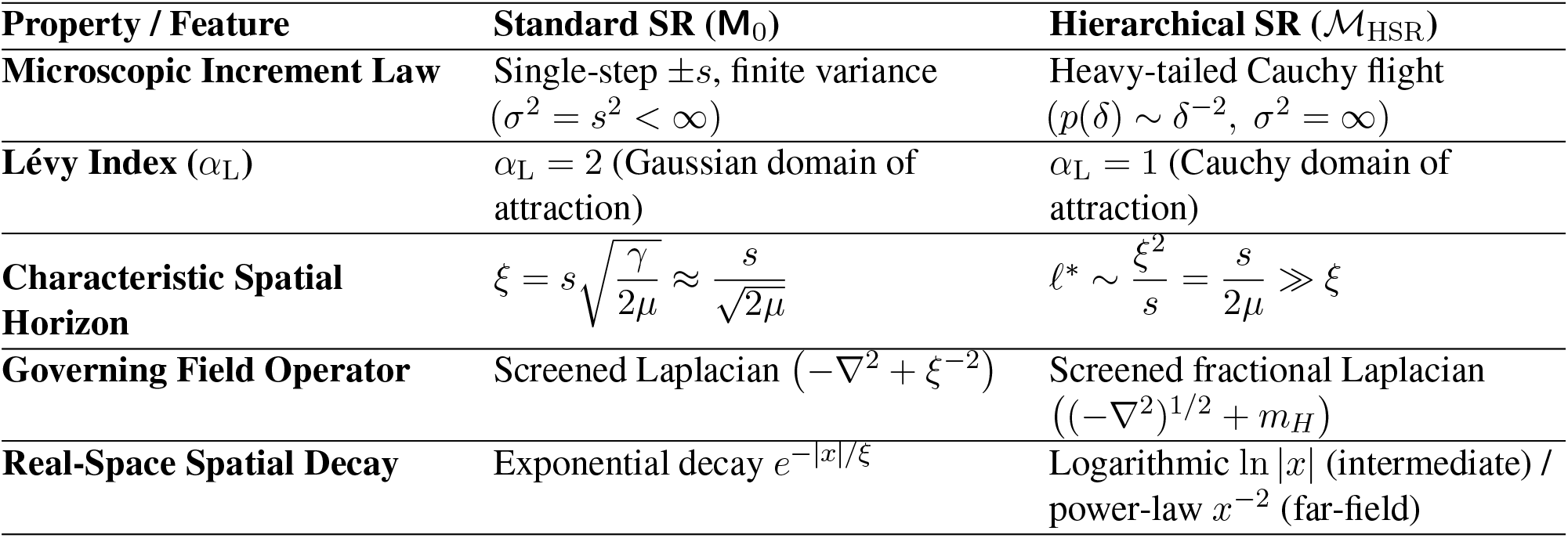
Key differences between SR and HSR.

| **Property / Feature** | **Standard SR (M<sub>0</sub>)** | **Hierarchical SR ($\mathcal{M}_{\mathrm{HSR}}$)** |
|---|---|---|
| **Microscopic Increment Law** | Single-step ±*s*, finite variance (*σ*<sup>2</sup> = *s*<sup>2</sup> < ∞) | Heavy-tailed Cauchy flight (*p*(*δ*) ∼ *δ*<sup>−2</sup>, *σ*<sup>2</sup> = ∞) |
| **Lévy Index (*α*<sub>L</sub>)** | *α*<sub>L</sub> = 2 (Gaussian domain of attraction) | *α*<sub>L</sub> = 1 (Cauchy domain of attraction) |
| **Characteristic Spatial Horizon** | $\xi = s\sqrt{\frac{\gamma}{2\mu}} \approx \frac{s}{\sqrt{2\mu}}$ | $\ell^* \sim \frac{\xi^2}{s} = \frac{s}{2\mu} \gg \xi$ |
| **Governing Field Operator** | Screened Laplacian (−∇<sup>2</sup> + *ξ*<sup>−2</sup>) | Screened fractional Laplacian ((−∇<sup>2</sup>)<sup>1/2</sup> + *m*<sub>*H*</sub>) |
| **Real-Space Spatial Decay** | Exponential decay *e*<sup>−\|*x*\|/*ξ*</sup> | Logarithmic ln \|*x*\| (intermediate) / power-law *x*<sup>−2</sup> (far-field) |

In summary, standard SR operates as a single-scale, local exponential smoothing filter that is fundamentally constrained by Gaussian diffusion dynamics (−∇^2^). By organising options according to the harmonic eigenvectors of the symmetric random-walk dynamics of the environment, the HSR transforms the predictive representation into a scale-free, non-local transport operator (|∇|). This structural shift resolves the single-parameter bottleneck of standard SR, and enables scale-invariant representation that spread spectral power across wider spectral bands.

### 3 Quasi-NMF decomposition

#### 3.1 Problem statement

As discussed in the Main text and Methods, the place cells are the shared overcomplete basis that could accurately reconstruct the hierarchical predictive map corresponding to arbitrary behavioural policy. Concretely, given a set of tasks, *G*, for any probability distribution, *p*(·)^2^, over *G*, we wish to solve the following optimisation problem.

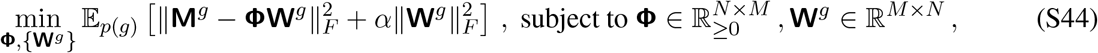

where *N* and *M* are the number of states and number of neurons, respectively, with *M* ≫ *N* . We take the derivative of L with respect to **W**^*g*^ and solve for **W**^*g*^.

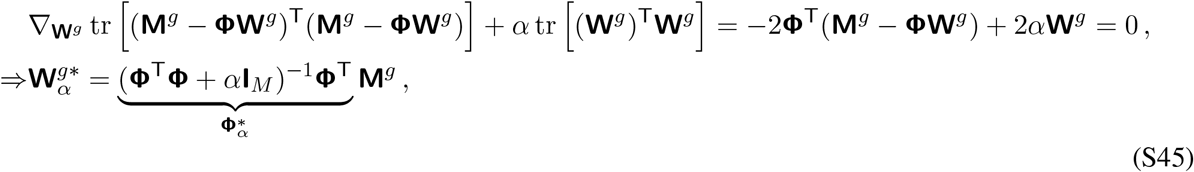

We see that the ridge regression penalty naturally deals with the degeneracy introduced by the overcomplete decomposition. Now inserting 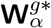 back into *L*.

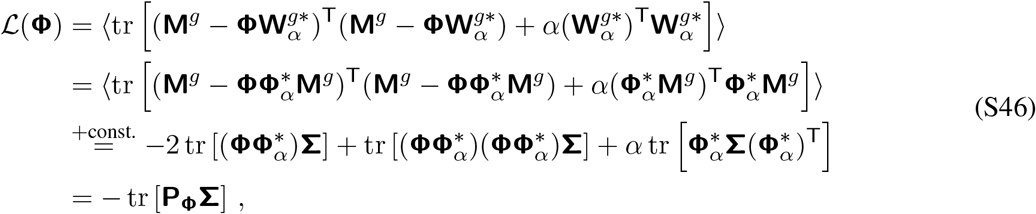

where **Σ** = E_*p*(*g*)_ [*M*^*g*^ (*M*^*g*^ )^T^] and **P**_**Φ**_ = **Φ** (**Φ**^T^**Φ** + *α***I**_*M*_)^−1^ **Φ**^T^. Note that the above derivations leveraged two properties of 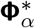:

- 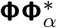 is symmetric: 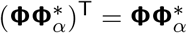;
- 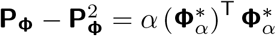;

#### 3.2 Gradient-based updates

For simplicity, we adopt the following notations.

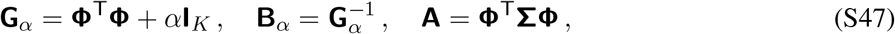

The loss function can then be re-expressed as following.

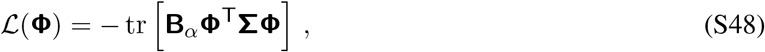

Taking the gradient with respect to **Φ**.

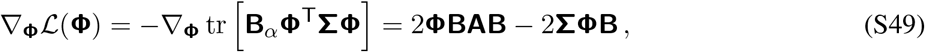

We further adopt the following notations.

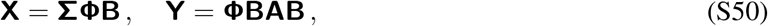

The gradient can hence be re-expressed as ∇_**Φ**_*L*(**Φ**) = 2(**Y** − **X**).

Due to the non-negativity constraint, we wish the gradient updates could preserve the non-negativity of **Φ** instead of applying further projections or hard thresholding. We take inspirations from the non-negative matrix factorisation (NMF)^5^ algorithm, and leverage the following trick that decomposes each matrix entry into their positive and negative parts.

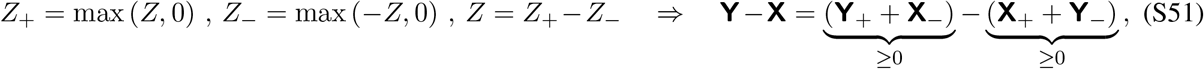

Hence the gradient becomes the difference between two non-negative matrices, ∇_**Φ**_L(**Φ**) = 2(**Y**_+_ + **X**_−_) − 2(**X**_+_ + **Y**_−_). Leveraging a dynamic learning rate, the following gradient updates always preserve the non-negativity of each matrix entry.

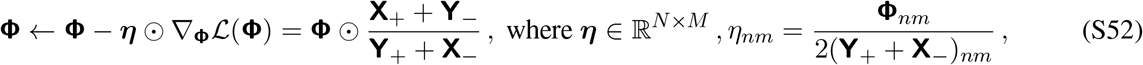

Note that in practice, we applied a sparsity (L1) penalty to the learned basis.

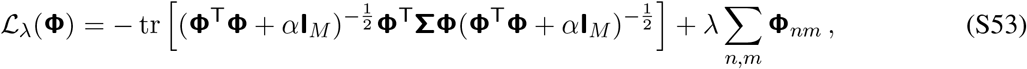

The gradient-based update rule can be easily modified to reflect the incorporation of the Lasso regularisation.

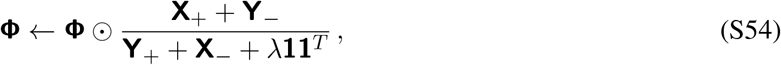

##### 3.2.1 Alternative parametrisation through explicit gain modulation

Given the derived gradients in the previous section such that the optimisation objective decreases monotonically whilst never violating the non-negativity constraint, we consider a different parameterisation with explicit gain modulation, **Φ** = **U**diag(**a**), with unit-norm columns ||**U**_:,*m*_ ||= 1, **U** ≥ 0, **a** ≥ 0. This gain-modulated parametrisation allows decoupling of the spatial topology of learned basis functions from their overall activation strength. Directly penalising an unnormalised basis matrix **Φ** via an L1-regularisation introduces undesirable coupling between an atom’s spatial extent and its regularisation cost, systematically penalising broad basis functions. We hence include an an aditional L1-penalty on **a** in the overall objective function (cf. Equation S48).

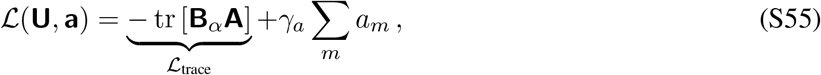

We can then derive the gradients wrt **U** and **a** by leveraging the derived gradients wrt **Φ** (Equation S49) and the definition Φ_*nm*_ = *U*_*nm*_*a*_*m*_, with an additional application of the chain rule.

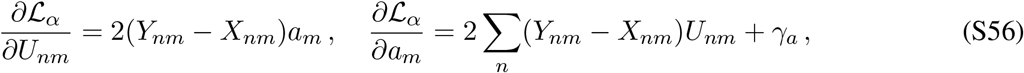

Given the above gradients, we can again leverage the NMF-style adaptive learning rate for ensuring the gradient-based updates never violate the non-negativity constraints (Equation S52).

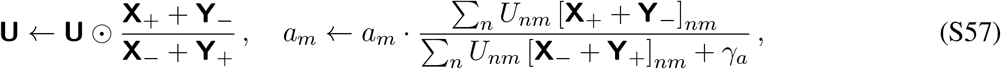

##### 3.2.2 Dealing with unstable training

In each iteration, the aforementioned optimisation procedures perform the following operations.

- **U***-update*: update **U** according to Equation S57. By construction, the trace loss decrease given **U**^′^, L_trace_(**Φ**^′^) ≤ *L*_trace_(**Φ**).
- *Renormalisation step*: divides each column **U**^′^_:,*m*_ by its norm *c*_*m*_ = ∥**U**_:,*m*_∥_2_ and absorbs *c*_*m*_ into *a*_*m*_.

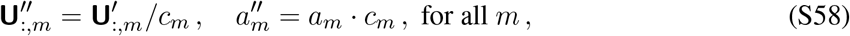 This leaves **Φ**^′′^ = **U**^′′^ diag(**a**^′′^) = **Φ**^′^ unchanged, hence the trace loss is unchanged and is still lower than the previous iteration. However, the L1-penalty on **a** changed.

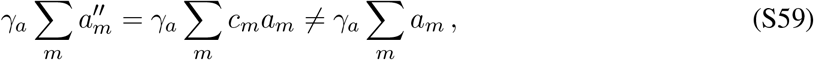

Hence if the U-updates causes columns to grow on average (⟨*c*_*m*_⟩ *>* 1), the total loss would go up.
- **a***-update*: update **a** according to Equation S57, decreases the total loss w.r.t. **a**, but the damage from the renormalisation step may outweigh it.

Hence the net per-iteration total loss is therefore not guaranteed to be monotonically decreasing.

To mitigate the instability in the optimisation process, we incorporate the unit-norm constraint 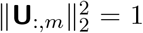 directly into the loss objective and associated gradients, via a per-column Lagrangian multiplier. In this case, the renormalisation step becomes a numerically stable measure and **a** is left untouched during the **U**-update.

The augmented loss function is shown as following.

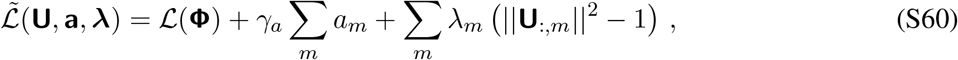

Taking the gradient of 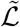 w.r.t. *U*_*nm*_ and set to 0, we have the following.

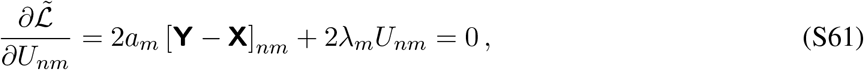

Multiply both sides of the above equation by *U*_*nm*_ and sum over *n*, and leverage the constraint that 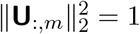, we have the following.

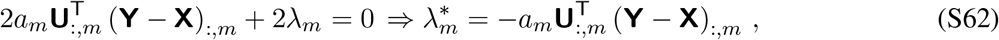

By defining 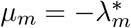, we can then compute the Lagrangian-augmented gradients.

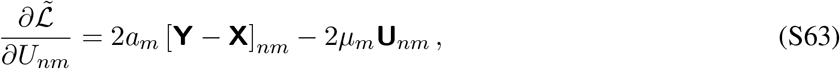

Note that this is the Riemannian gradient on the sphere: it removes the component of the Euclidean gradient that is parallel to **U**_:,*m*_, hence the update only changes the direction of **U**_:,*m*_, and not its scale ^6^.

To derive the NMF-style gradient-based updates, we again perform the positive-negative split of *µ*, defining the following quantities.

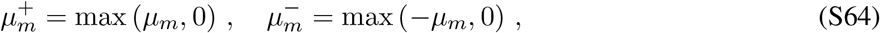

This hence decomposes the Lagrangian gradients as following.

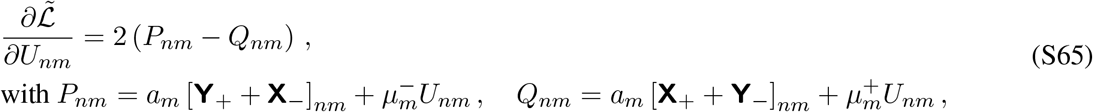

Hence the NMF-style gradient updates for **U** now becomes the following.

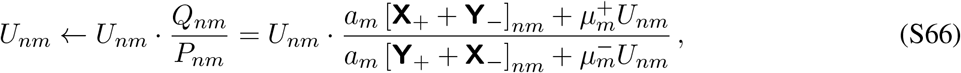

And after the **U**-update, we renormalise **U** for numerical stability only, whereas **a** is left unchanged.

##### 3.2.3 Relationship with symmetric NMF and heuristic initialisation scheme

Given the final form of the trace loss of the full optimisation objective (Equation S46), we can re-express the objective by introducing the following loosely “whitened” representation, 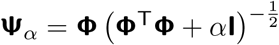 (note that 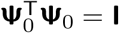).

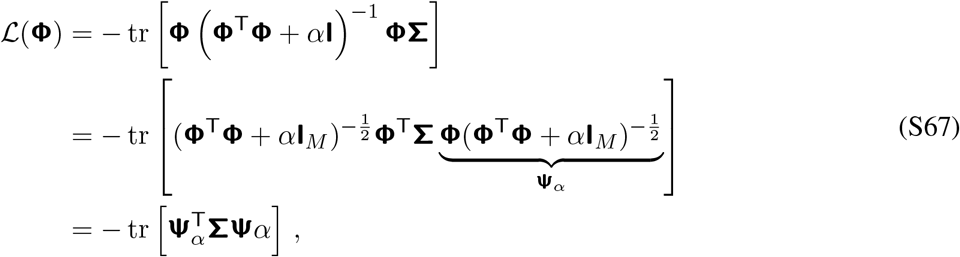

Compare with the symmetric NMF objective of **Σ** with factor matrix **Ψ**_*α*_ ^7,8^ as following.

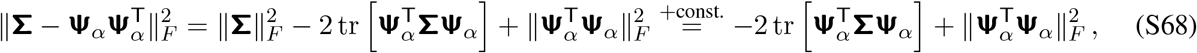

Note that taking the limit *α* → 0, we have that 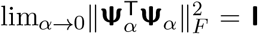, which is constant with respect to **Φ**. Hence the quasi-NMF optimisation objective contains exactly the data-alignment term of symmetric NMF of the second-moment matrix of the HSR. The difference lies in how factor norm, redundancy (Equation S60), and conditioning (see Section 3.5.2) are controlled.

Given the close analogy between symmetric NMF and the full quasi-NMF optimisation objective, we hence proposed a customised training procedure for improving the stability and efficiency of the overall decomposition, by heuristically initialising **Φ** as the factor matrix underlying symmetric NMF of **Σ** (given a similar iterative optimisation algorithm given adaptive learning rate scheduling; Section 3.2). Empirically, the initialisation scheme is more robust with respect to initialisation, and already provides a close-to-optimal solution of the full optimisation process (Extended Data Figure 9).

#### 3.3 Why decomposing the second-moment instead of raw HSR

Recall that the overall learning objective takes a quasi-NMF formulation.

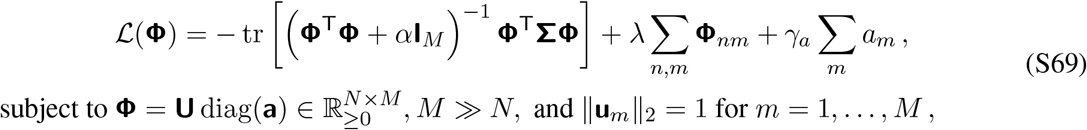

The input matrix **Σ** = *E*_*g*_[*M*^*g*^ (*M*^*g*^)^T^] is the expected second moment matrix of the SR/HSR, encoding the pairwise similarity between states in terms of their predictive trajectories, averaged over all policies. Below we list four reasons, each from a different perspective, for why decomposing this expected outer product is favourable.

- **Algebraic**. For each *g* ∈ *G*, the optimal value function takes the following form (assume vectorised notations in discrete state space).

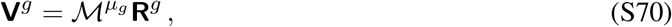

where *µ*_*g*_ is the optimal high-level policy for solving task *g* and **R**^*g*^ is the task-specific reward function. In addition to reconstruct the predictive map and represent hippocampal place cells, we expect the shared basis should also accurately encode the optimal value functions across tasks through linear reconstruction.

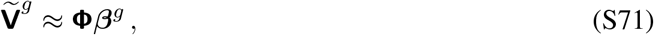

The optimisation objective for this value approximation problem (with L2-regularisation on the regression weights) is as following.

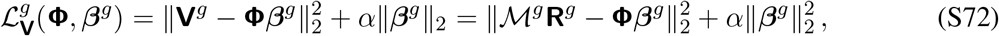

Taking the derivative with respect to ***β***^*g*^ and set to 0, we have the optimal weight vector for a fixed **Φ**.

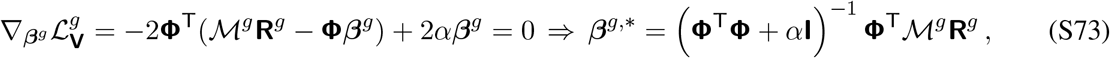

Substituting ***β***^*g*,*^, and we compute the regularised value approximation error.

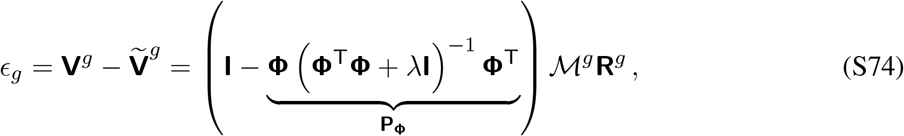

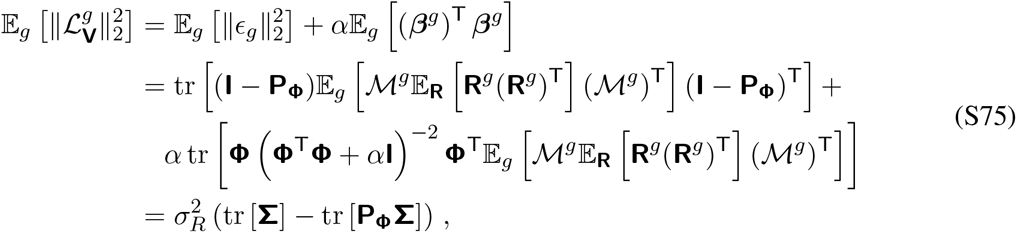

where we assumed **R**^*g*^ is isotropic across the task distribution, i.e., 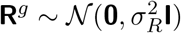, yielding 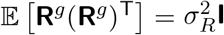. The last equality follows closely with our derivation of the simplification of the quasi-NMF objective (cf. Equation S46). Consequently, for any *α >* 0, the expected value approximation error simplifies to the ridge-regularised objective of the quasi-NMF optimisation (Equation S46).

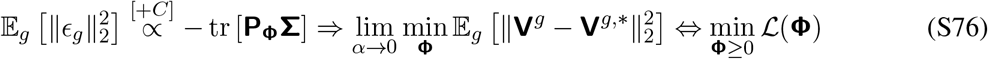

It is hence clear that minimising the quasi-NMF reconstruction loss with respect to **Σ** directly minimises the expected downstream value approximation error across all possible task reward configurations.
- **Statistical**. *M*^*g*^ represents a single realisation, and **Σ** is the two-point correlation function that describes how predictive occupancies co-vary across the full ensemble of task optimal policies. Therefore, extracting modes from *M*^*g*^ captures realisation-dependent noise and transient policy artefacts, whereas extracting modes from **Σ** yields structural normal modes that capture task-invariant geometry.
- **Spectral**. From Section 2, we know that 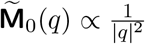 (Equation S10), and 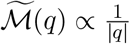 (Equation S37). Importantly, given the definition of **Σ**, its Fourier transform is the square of that of the raw predictive map.

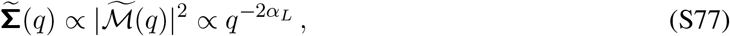 Hence the octave contrast (the log-ratio of energy contained between two adjacent spatial frequency bands separated by one octave) doubles for **Σ**.

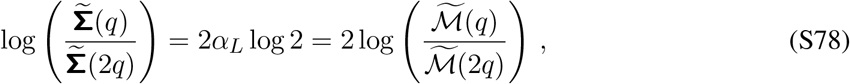

By doubling the octave contrast, **Σ** creates a steeper energy hierarchy between spatial octaves. This will push coarse modes further above fine modes in magnitude, and will sharpen the boundary between spatial scales, allowing sparsity penalties to cleanly prune scale-redundant atoms without distorting individual spatial profiles.
- **Symmetric NMF as soft-clustering**. By the definition of **Σ**, it is strictly symmetric and positive semi-definite. Hence, factorising **Σ** ≈ **ΨΨ**^*T*^ (Section 3.2.3) under non-negativity constraint is a continuous relaxation of kernel K-means clustering ^7^. In contrast, factorising *M* directly (which is asymmetric in general) would require an asymmetric factorisation, yielding distinct left and right singular vectors that break the geometric connection to spatial partitioning.

#### 3.4 Finite-sample reconstruction error bound on the quasi-NMF optimisation

In practice, the quasi-NMF optimisation is performed with respect to the finite-sample estimate of the second-moment of the HSR matrix (Equation S46), instead of the ground-truth second moment matrix. Here we derive the finite-sample error bound in using such empirical estimate.

Given *n* tasks, 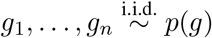, we estimate the second-moment matrix as following.

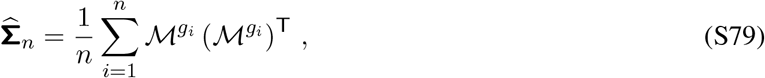

Recall the trace loss *L*^trace^ (**Φ**) = − tr [**P**_**Φ**_**Σ**], where **P** = **Φ** (**Φ**^T^ **Φ** + *α***I**)^−1^ **Φ**^*T*^ . We define the empirical loss function given the *n* tasks, 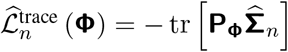. We further define the empirically and ground-truth optimal basis as 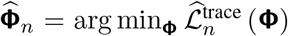 and **Φ**^*^ = arg min_**Φ**_ *L*^trace^ (**Φ**), respectively, over the constrained set *C*_*A*_ = {**Φ** = **U**diag (**a**) : **U** ≥ 0, ∥**u**_*k*_∥_2_ = 1, **a** ≥ 0, ∑_*k*_ *a*_*k*_ ≤ *A*} (see **A2** below). The finite-sample error of interest is as following:

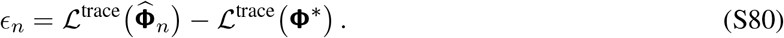

We need the following two key assumptions.

- **A1**. Tasks {*g*_1_, …, *g*_*n*_}are i.i.d. sampled from the task distribution *p*(*g*) (not necessarily uniform), and the task-dependent HSRs are bounded.

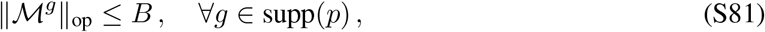

where ∥·∥_op_ is the operator (spectral) norm. Note that by definition, 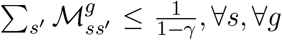, ∀*s*, ∀*g*. Consequently, 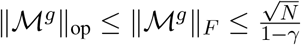, we can hence define 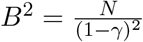 as an upper bound (conservative, not necessarily the tightest bound).
- **A2**. Under the gain-regularised parametrisation (Section 3.2.1), we assume the L1-norm of the gain modulations is bounded.

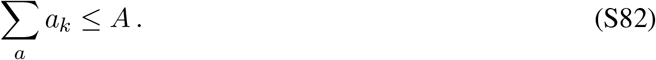 This has a direct consequence that the trace of the basis-dependent projection matrix is bounded (the bound is obvious by expressing **P**_**Φ**_ through the SVD of **Φ**).

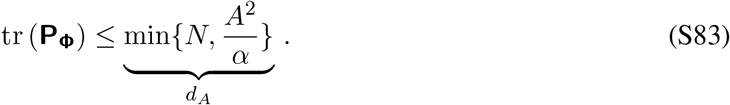

Note that *d*_*A*_ provides a measure of the effective reconstruction capacity of the regularised overcomplete dictionary, **Φ**.

To derive the finite-sample error bound, we will further leverage the following matrix-analogue of the scalar Bernstein inequality ^9^.

##### Theorem 3.1

(Matrix Bernstein inequality). *Let* **Y**_1_, …, **Y**_*n*_ *be i*.*i*.*d. random matrices of dimension d*_1_ × *d*_2_, *with* E [**Y**_*i*_] = **0**, *and* ∥**Y**_*i*_∥_*op*_ ≤ *L. Define the random summation*, 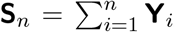, *and the corresponding matrix variance as following*.

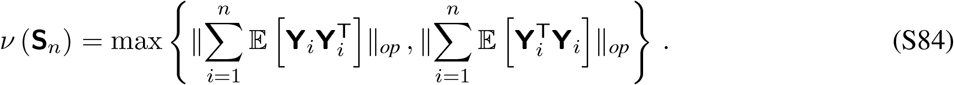

*Then for any t* ≥ 0, *we have the tail probability as following*.

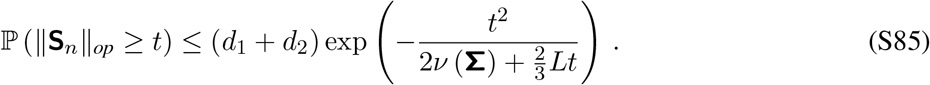

*Proof*. Please refer to Section 6.1 in Tropp, 2015^9^ for proof.

##### Theorem 3.2

(Finite-sample excess reconstruction error bound). *For any confidence level δ* ∈ (0, 1), *with probability at least* 1 − *δ over the sampled tasks*,

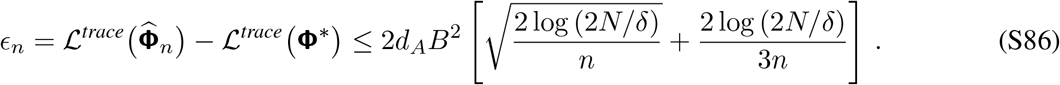

*Proof*. We firstly define 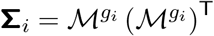. We immediately know that **Σ**_*i*_ ⪰ 0, and E [**Σ**_*i*_] = **Σ**. By **A1**, we have the following:

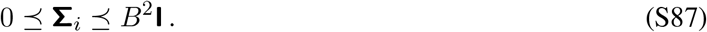

We then define **Z**_*i*_ = **Σ**_*i*_ − **Σ**, and **Z**_*i*_ are independent, zero-mean, and symmetric matrices, such that ∥**Z**_*i*_∥_op_ ≤ *B*^2^. To see why the last bound holds, we again leverage **A1**,

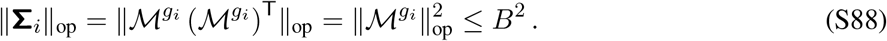

We hence have that 0 ⪯ **Σ**_*i*_ ⪯ *B*^2^**I** and 0 ⪯ **Σ** ⪯ *B*^2^**I**. Combining the two inequalities into the definition of **Z**_*i*_, we have the following inequality.

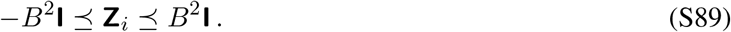

Since **Z**_*i*_ is symmetric, all of its eigenvalues must lie within [−*B*^2^, *B*^2^], we hence achieve an upper bound on the operator norm of **Z**_*i*_.

We further have that 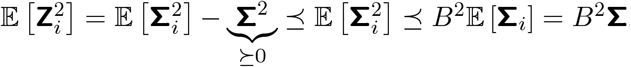, where the last inequality is given by 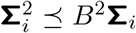. Given **Z**_*i*_ is zero-mean and symmetric for all *i*, we can derive an upper bound on the matrix variance of the summation 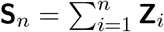.

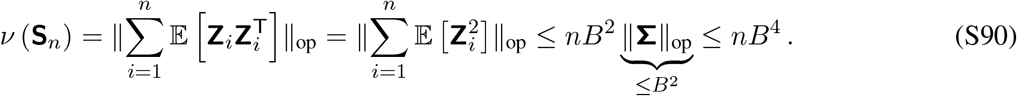

Substituting **Y**_*i*_ as **Z**_*i*_, *d*_1_ = *d*_2_ = *N, L* = *B*^2^, 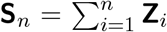, and *v* (**S**_*n*_) ≤ *nB*^4^ into the matrix Bernstein inequality (Equation S85),

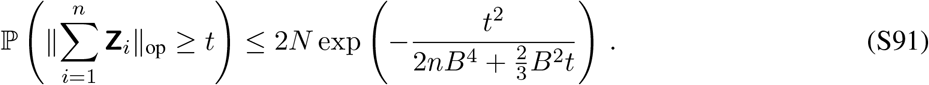

Note that the above inequality provides a bound on the summed, instead of the expected error. By the definition of 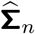 (Equation S79), we have that 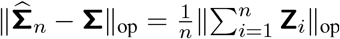. By further replacing *t* = *nϵ*, we have the following bound for the average error.

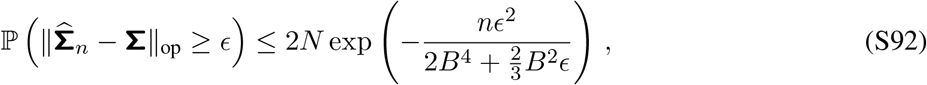

We now derive the threshold 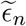 such that 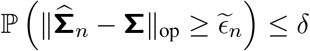 for a pre-specified confidence level *δ*.

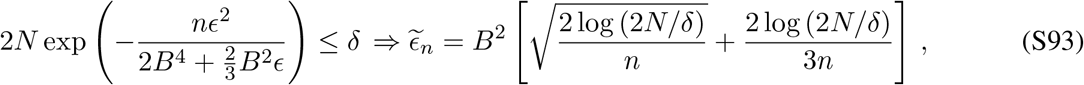

is a valid choice for 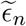 (the positive root of the quadratic function in *ϵ* is upper-bounded by this, leveraging the inequality 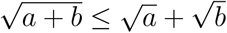.

Inserting the concentration bound on **Σ** back into the trace loss objective, we can derive the upper bound on the empirical trace loss, 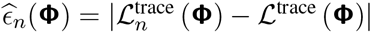.

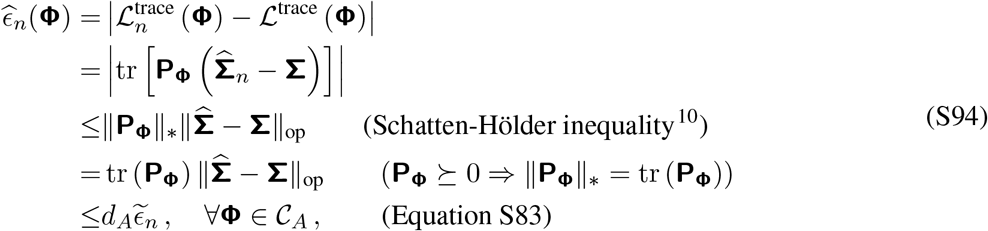

where ∥·∥_*_ is the nuclear norm (sum of all singular values), and 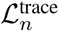 is the trace loss with respect to the empirical estimate of the second-moment matrix given *n* tasks.

Finally, we re-express the finite-sample error bound (Equation S86) as following.

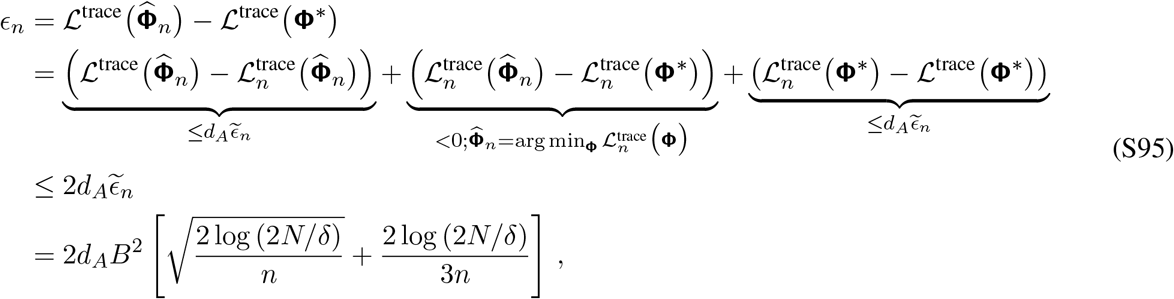

which decays on the order of 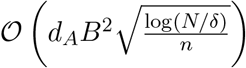 with the number of tasks, *n*.

#### 3.5 The quasi-NMF objective induces non-unimodal representation

We examine the properties of the non-negative factors, **Φ**, through its own singular value decomposition (SVD), **Φ** = **ΨTX**^T^, with **Ψ** ∈ ℝ^*N* ×*N*^ orthogonal, **T** = diag (*τ*_1_, …, *τ*_*N*_ ), and **X** ∈ ℝ^*M* ×*N*^ with orthonormal columns. We also define **X**_⊥_ that complement **X** to an orthonormal basis of ℝ^*M*^ . Leveraging the SVD and after some algebra, we can show that the trace loss in the overall objective can be re-expressed as following.

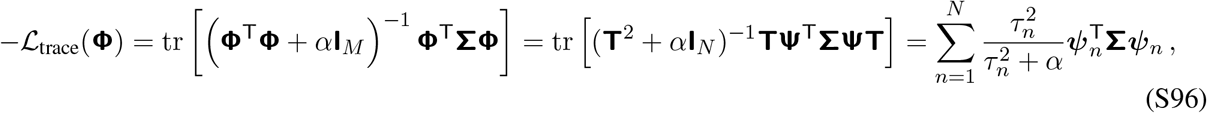

where *ψ*_*n*_ is the n-th column of **Ψ**. We hence see that the frame (space spanned by the non-negative factors) contributes to the trace loss only through its own left singular vectors and singular values. Individual atoms (basis) enter only via how they build their singular vectors and values. We further denote the weights.

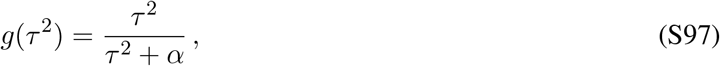

We note that 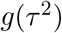 saturates at 1, and pushing *τ* ^2^ ≫ *α* yields minimal gainings. Moreover, at *α* = 0, *g* ≡ 1, and the trace loss becomes tr [**P**_range**Φ**_**Σ**] = tr **Σ**, where **P**_range**Φ**_ = **Φ** (**Φ**^T^ **Φ**)^−1^ **Φ**^T^ is the orthogonal projection matrix onto the column space of **Φ**, and the equality holds since we assume **Φ** is a spanning frame. Since tr **Σ** is a fixed scalar constant determined entirely by the data, the loss then has zero dependence on the actual shapes of structure of the columns in **Φ**, and every spanning frame **Φ** achieves the exact same maximum trace objective, which makes the loss landscape completely flat, hence the optimisation problem becomes degenerate. Therefore, the ridge regression penalty initially introduced over task-dependent loading weights (Equation S44) is not an ad-hoc fix for the degeneracy introduced by the overcomplete decomposition, but pushes the optimisation towards non-trivial solutions.

We then suppose the frame can re-orient its singular vectors freely, and choose to orient ***ψ***_*j*_ to be aligned to **Σ**’s eigenvectors, with corresponding eigenvalues 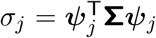. Under these assumptions, the optimisation problem becomes the following.

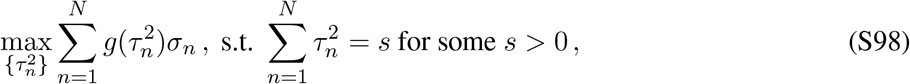

Given the Lagrange multiplier, *v >* 0, the loss function is as following.

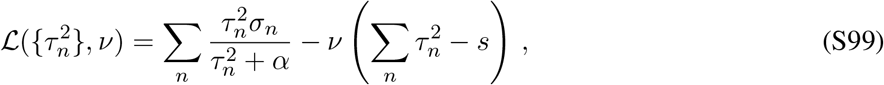

Maximising the loss function gives us the following optimal solution.

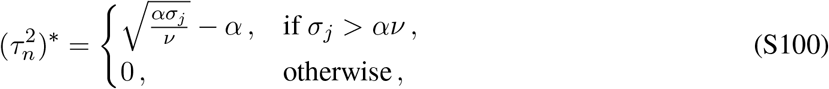

Hence the n-th principal direction survives if and only if *σ*_*n*_ *> σ*_*c*_ = *αv*, and when above threshold, 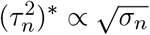. The latter point makes it clear that the frame *half-whitens*: energy are allocated as the square root of the spectrum, instead of matched-filtering (∝ *σ*) or whitening (∝ 1*/σ*).

Leveraging the infinite-lattice assumption, for a locally translational invariant frame in the bulk, ***ψ***_*n*_’s are Fourier modes, and the squared singular values become the spectral power of **Φ** at wavenumber *q*_*n*_ (or equivalently, the n-th eigenvalues of the frame autocorrelation operator **ΦΦ**^T^).

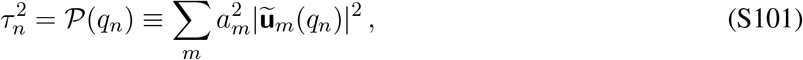

where we have leveraged the gain-modulation decomposition (**Φ** = **U** diag(**a**); Section 3.2.1). Furthermore, in the bulk states, the predictive similarity matrix **Σ** is spatially homogeneous and translation-invariant, and its eigenvalues are identical to its power spectrum, 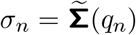. We hence have that 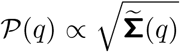 whenever 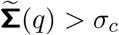. ∝

For the HSR, 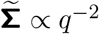 (Equation S35; S77), hence *P* ∝ *q*^−1^. Since 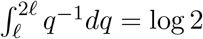 is independent of where the octave sits, the frame carries equal energy per octave. The preservation of spectral power across spatial hierarchy is not automatically inherited from the HSR (its Lévy index is *α*_*L*_ = 1), and only in combination with the half-whitening under the quasi-NMF optimisation forces the emergence of such scale-invariance.

In contrast, for the standard SR, the power spectrum 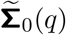 is dominated by the mass term for frequencies smaller than the mass-dependent frequency cutoff (*q < q*_*c*_ ≈ 1*/ξ*), making the spectrum flat in this case. Over high frequencies (*q > q*_*c*_), the *q*^−2^ kinetic term dominates, yielding a steeper decay, 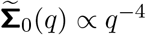. Then apply the half-whitening rule, *P*(*q*) is constant over low frequencies and scales with *q*^−2^ over high frequencies. The energy decay in standard SR falls off rapidly in frequency space, dropping below *σ*_*c*_ almost immediately. As a result, the standard SR cannot build a multi-scale hierarchy (a “ladder” spanning multiple spatial octaves). The associated frame would then allocate most of its capacity to a single, contiguous low-frequency band *q* ∈ [0, 1*/ξ*], leading to basis functions represented by single-scale blobs.

##### 3.5.1 Earnings of atoms

We now turn our attention to individual atoms (non-negative factors; **u**_*m*_ ≥ 0). We firstly list several elementary properties of non-negative vectors, which will be useful in subsequent analysis.

- **Orthogonality = disjointness**. For **u, v** ≥ 0, ⟨**u, v**⟩ ≥ 0, with equality if and only if **u**(*x*)**v**(*x*) = 0 for all *x*, i.e., **u** and **v** have disjoint supports.
- **Effective support**. For **u** ≥ 0 with ∥**u**∥_2_ = 1, we define 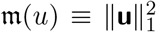. Applying Cauchy-Schwarz inequality, we have that 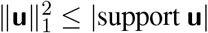, with equality if and only if **u** is flat on its support. We hence define *m* as the effective number of sites/fields occupied.
- **No cancellation**. Suppose **V** = ∑_*m*_ **w**_*m*_***φ***_*m*_ ≥ 0, with **w**_*m*_ ≥ 0 and ***φ*** ≥ 0, hence support(***φ***_*k*_) ⊂ support(**V**).

As a consequence of the third property, NMF leads to parts-based representation ^5^ since destructive interference is mathematically forbidden. However, we note that non-negativity alone does not guarantee localised representation, since it does not prevent broad functions from being added.

Define the predictive energy associated with atom *m* (Equation S69) as the contribution to the trace loss from the normalised m-th atom.

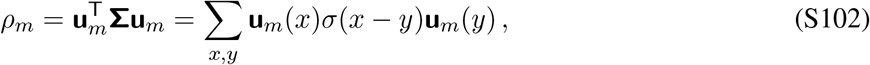

where 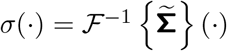 denotes the real-space spatial correlation function. Note that we have leveraged the in-bulk translation invariance of the predictive similarity kernel *σ*(*x* − *y*) ≈ **Σ**_*xy*_.

We further define the following useful quantities: *n*_*m*_ ∈ ℕ as the number of distinct spatial bumps/fields for the m- th atom; *w*_*mi*_ as the spatial width of the i-th field of the m-th atom^3^; *W*_*m*_ as the envelope width (total spatial span across all scattered lobes, from the leftmost to the rightmost fields) of the m-th atom; *m*_*m*_ ≡ *m*(**u**_*m*_) = *n*_*m*_*w*_*m*_ as the effective support size of the m-th atom; *η*_*m*_ ≡ *m/W*_*m*_ as the duty cycle, encoding the fraction of the envelope occupied by active lobes.

We separate the treatment of the predictive energy into two regimes.

- **Regime A** 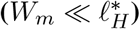. In this case, the spatial span of the atom fits inside a correlation length, then we can approximate *σ*(*x y*) = *σ*(0) across the atom’s spatial extent. Hence the energy associated with these atom is as following.

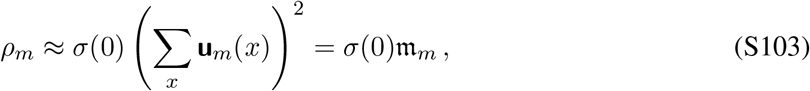 Under this regime, an atom’s earning depends strictly on its effective mass *m*_*m*_, and is completely blind to how its support is arranged: one interval of *m*_*m*_ sites, or *n* lobes of *m*_*m*_*/n*_*m*_ sites contributes identical energy.
- **Regime B** 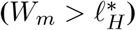. In this case, *σ*(*x* − *y*) decays, we hence need to consider the overlap between different lobes within the same atom. Suppose 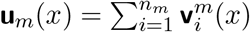, where each 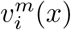 denotes a lobe with a narrow spatial width *w*_*m*_, and 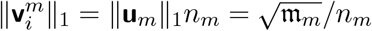. The predictive energy can hence be decomposed as following.

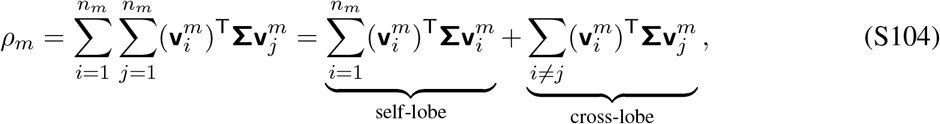
  – **Self-lobe**. Assume each individual field is narrow 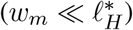, we can re-apply the energy expression derived in regime A.

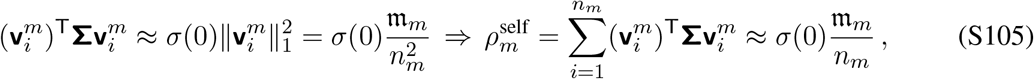
  – **Cross-lobe**. We apply first-order approximation to the HSR transition kernel 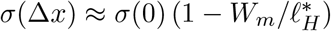 for small separations Δ*x*, and substitute it into the cross-lobe summation.

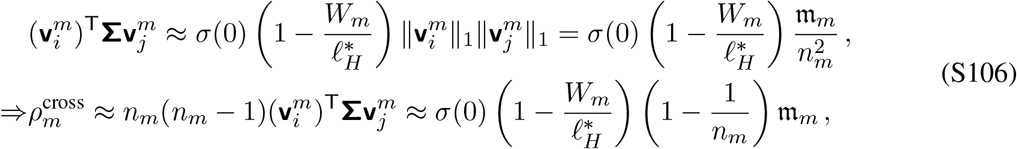

Combining the two terms together, we have the following error correction ratio for 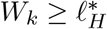.

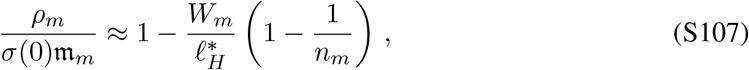

We can observe from the error correction term that scattering an atom’s mass over an envelope *W*_*m*_ is free while 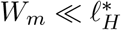, and become increasing prohibitive as 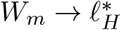.

##### 3.5.2 Costs of atoms

From the quasi-NMF loss objective (Equation S69), each atom pays exactly proportional *a*_*k*_.

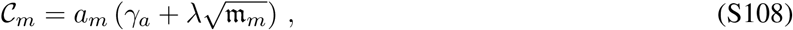

The earnings of each atom is *ρ*_*m*_ ∝ *m*_*m*_, whereas the cost is 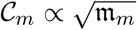 hence the net contribution is increasing in *m*_*m*_, and atoms grow their support unless something else stops them.

Instead of an explicit penalty, it is the *Gram-conditioning* that stops this growth. Let **G** = **U**^T^**U** be the Gram matrix **G** ∈ *R*^*M*×*M*^, which is positive semi-definite, its rank ≤ *N*, unit-diagonal (∥**u**_*m*_∥_2_ =1), and entrywise non-negative. Its eigenvalues *λ*_*m*_ satisfy ∑_*m*_ *λ*_*m*_ = *M*, with at most *N* non-zero eigenvalues. Given the SVD-derive trace-loss objective (Equation S96), for *g*(*τ*^2^) to saturate, 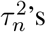’s need to be spread out rather being concentrated. Note that by the definition of 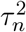 (eigenvalues of **ΦΦ**^T^ = **U** diag(**a**^2^)**U**), eigenvalues of **G** is closely related to 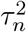 (via re-ordering, 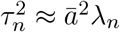, for *n* = 1, …, *N*, and *λ*_*n*_ = 0 for *n* = *N* + 1, …, *M*, where 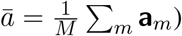, hence the concentration of 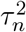’s is similarly reflected in the concentration of *λ*_*m*_. We note that the concentration of *λ*_*m*_’s can be evaluated through 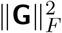.

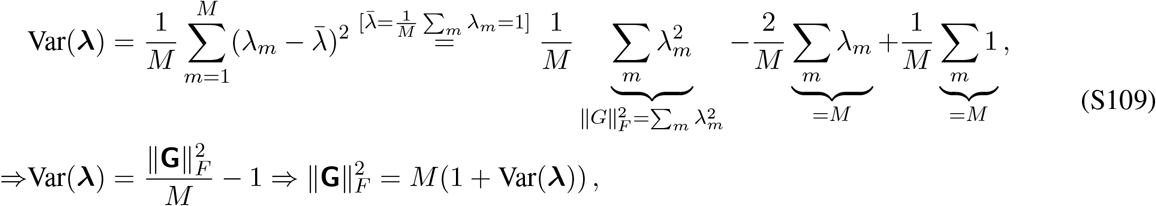

We hence evaluate 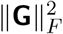.

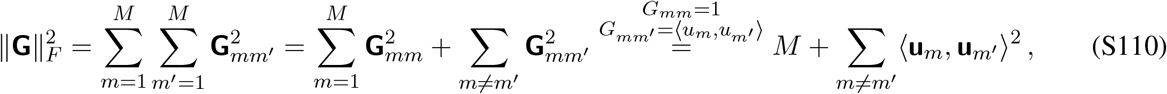

We hence see that

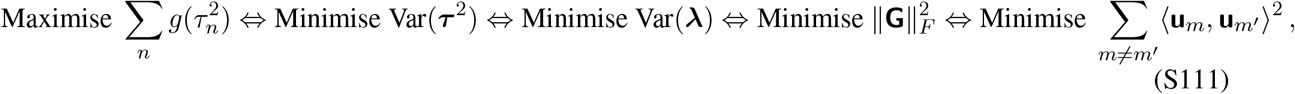

Hence from a spectral perspective, within a band of similar *σ*_*n*_, at a fixed budget 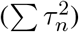, the concavity of *g*(·) favours spreading 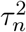 evenly, leading to a flat spectrum for the Gram matrix within that band and small overlap 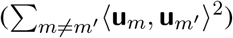.

From a spatial perspective, minimise spectral variation is algebraically equivalent to minimise 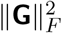, which enforces minimal spatial overlap between atoms.

##### 3.5.3 How much spatial overlap is unavoidable?

To determine the minimum total overlap enforced by the overcomplete decomposition (*M* ≫ *N* ), we sum all entries of **G**.

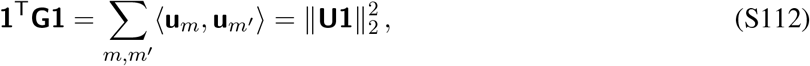

Define the atom summation vector as 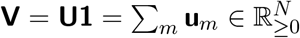. We apply the Cauchy-Schwartz inequality.

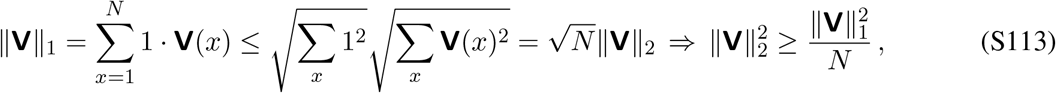

Recall 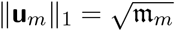, we have that

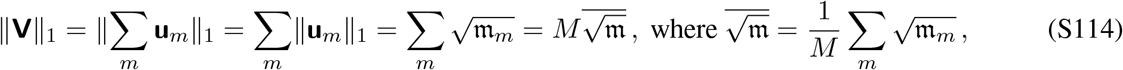

We hence substitute this expression back into the inequality.

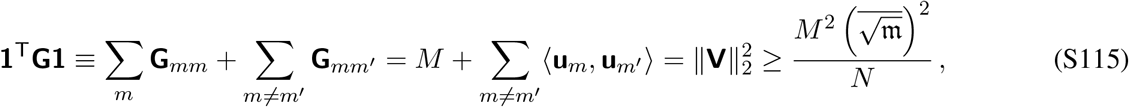

Thus, we can compute the lower bound on the mean overlap between any two atoms.

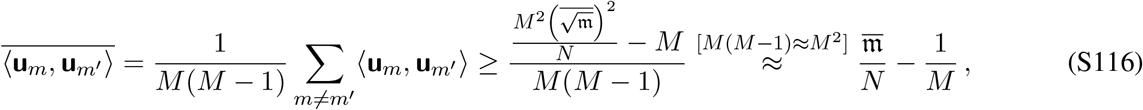

where 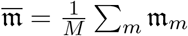. We observe that the total overlap is bounded below by the support budget and cannot be evaded.

However, we note that maximising *g*(*τ*^2^) penalises squares (Equation S111). Suppose we fix the total overlap assigned to the m-th atom across its partners 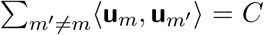 for some *C >* 0. Assume that there are *p* active overlapping pairs, such that 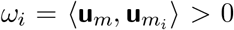 for *i* = 1, …, *p*, with 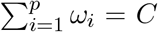. Then by the Cauchy-Schwarz inequality, we have that

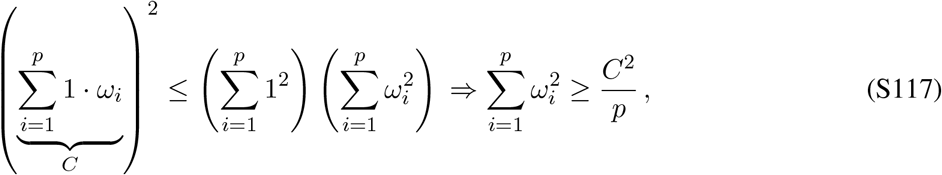

We note that the lower bound on the squared total overlap is decreasing in *p*.

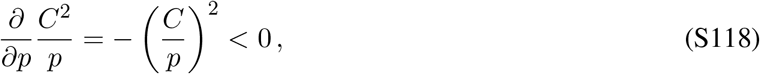

We hence have established that distributing a fixed budget on total overlap, *C*, across a large number of partners *p* (many small overlaps) yields a strictly smaller penalty than concentrating *C* onto a small number of partners (few large overlaps).

##### 3.5.4 Scattered beats compact

Why would the quasi-NMF optimisation produce multi-lobed factors instead of compact single-bump factors? To see this, we consider two types of atoms **u**_*C*_ and **u**_*S*_ with identical effective support mass 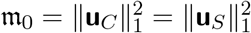.

- **Compact atom u**_*C*_, with envelope width *W*_*C*_ = *m*_0_, hence its duty cycle is *η*_*C*_ = *m*_*C*_*/W*_*C*_ = 1.
- **Scattered atom u**_*S*_, with *n*_*S*_ separate, interleaved lobes, each of width *w*_*S*_ = *m*_0_*/n*_*S*_, spread across an envelope *W*_*S*_ = *m*_0_*/η*_*S*_, with *η*_*S*_ *<* 1.

Assume 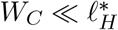 and 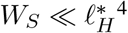 ^4^, then the earnings of both atoms are roughly similar.

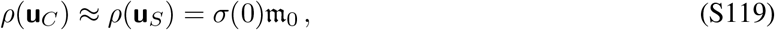

Moreover, since they have identical support mass, the associated L1-sparsity penalties are also identical (Equation S108).

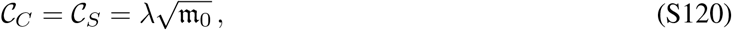

Since both the earnings and sparsity penalties are identical, the optimisation choice between compact and scattered representations is decided entirely by the conditioning penalty 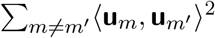 (Equation S111).

Suppose *R* atoms compete for covering a spatial track of length Λ.

- **Compact atoms**. In a contiguous region of length Λ, at most Λ*/m*_0_ compact atoms of width *m*_0_ can sit side-by-side without overlapping. Hence two atoms intersect with probability ∼ *m*_0_*/*Λ, and each atom overlaps with roughly 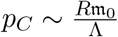 other atoms. Over the intersection region, their inner product is 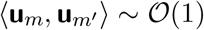, hence the sum of square overlaps is

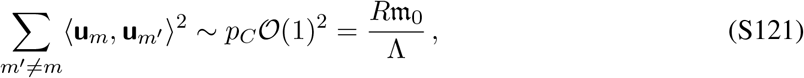
- **Scattered atoms**. A scattered atom has an extended envelope *W* = *m*_0_*/η*, hence spans a larger spatial domain and encounters more partner atoms.

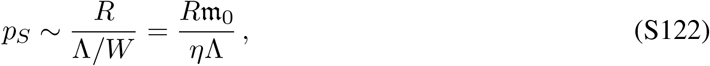 Since the lobes are scattered with duty cycle *η*, two independent scattered atoms sharing an envelope intersect only on a fraction *η* of their total mass. The pairwise inner product drops to 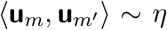, hence the sum of square overlaps now becomes

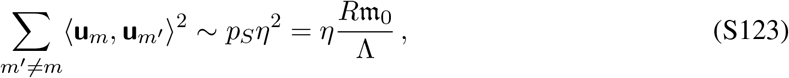

We have now established that scattering mass into *n* lobes with duty cycle *η* reduces the conditioning penalty directly by a factor of *η*, at zero cost to its contribution to the earnings or sparsity penalty.

We note that the total linear overlap is nevertheless unavoidable.

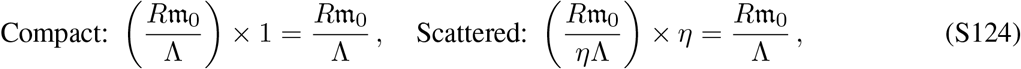

However, the objective penalises the sum of square overlaps (Equation S111), increasing the partner count *p* by 1*/η* (Equation S118) and reducing individual overlaps by *η*, scattering achieves a 1*/η* reduction in the squared overlap sum.

The maximum spatial envelope *W*_max_ an atom can occupy is constrained by two independent physical limits: (a) the environment length *L*, and (b) the spatial correlation horizon 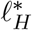 (beyond 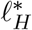, the cross-lobe incoherence causes predictive energy earnings *ρ* to drop rapidly; Equation S107). We hence take the tightest upper bound.

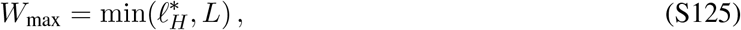

The octave span ℜ ≡ *W/w* measures the encoded dynamic range, i.e., the ratio between total envelope and the finest lobe resolution *l*_min_. Given *K* eigenoptions, we have that *l*_min_ = *L/K*. We hence have

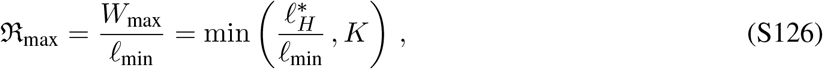

Two competing forces sets the content of scattering. As overcompleteness *M* increases, spatial collisions between atoms become severe. To minimise the sum of square overlaps, the optimisation would drive the duty cycle down *η* → 0, and effectively drives the envelope up. Without such redundancy (*M* ≈ *N* ), there is no conditioning pressure, hence the scattering is no longer desirable. On the other hand, the redundancy alone would drive *W* → ∞, but as 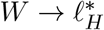, the cross-lobe penalty (Equation S107) destroys the atom’s earnings, making scattering beyond 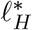 unaffordable.

Taking inspirations from continuous signal processing, a wavepacket is characterised by its spatial extent Δ*x* ∼ *W*, and its spectral bandwidth 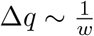. This gives us a novel interpretation of the octave span ℜ, as the phase-space volume occupied by the atom.

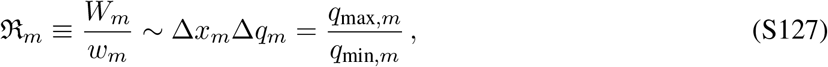

where 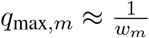 and 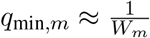 are the high- and low-frequency cutoffs, respectively.

As its name suggests, the octave span also defines the number of spatial octaves contained within the frequency band [*q*_min,*m*_, *q*_max,*m*_].

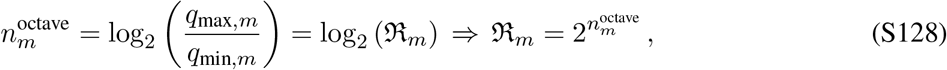

#### 3.6 Single-scale feature collapse of standard SR

The collapse of the frame for standard SR into a single spatial-scale representation is the result of a “pincer movement” that squeezes an atom’s inner feature size *w*_*m*_ and global envelope size *W*_*m*_ into the exact same value (*ξ*).

- **Lower bound on the lobe width**, *w*_*m*_. From Section 3.5, we established that the aggregate frame power spectrum 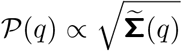. For standard SR, we have that 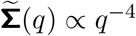 for wavenumbers *q >* 1*/ξ*. The quartic spectral scaling falls off extremely fast, hence 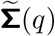 drops below the survival threshold *σ*_*c*_ almost immediately past *q* = 1*/ξ* (Equation S100). This hard frequency cutoff (*q*_max_ ≲ 1*/ξ*) means that the dictionary cannot encode any spatial resolution finer than 1*/q*_max_ ≈ *ξ*. Hence the minimum width of a single lobe *w*_*k*_ is bounded below by the correlation lengthscale.

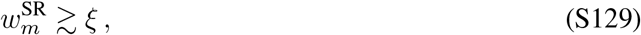
- **Upper Bound on the envelope size**, *W*_*m*_. Given the Fourier transform of the standard continuous SR transfer kernel (Equation S18), the second-moment matrix 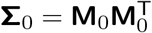 squares this transfer function in frequency space

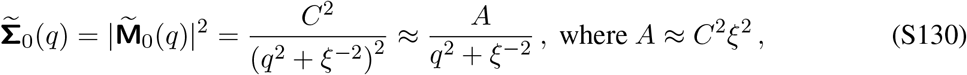

where *A* defines the effective amplitude scale of the spectrum. To re-express 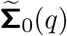 in the generic Lorentzian form, 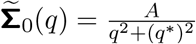, where *q*^*^ is the cutoff wavenumber where the spectrum transitions from a flat plateau (mass-dominated) to a power-law decay (kinetic-dominated). To retrieve the express of *q*^*^, we examine the zero-lag correlation.

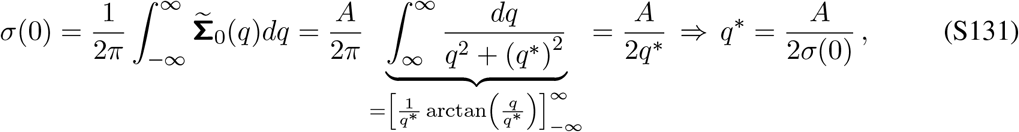

We hence evaluate the zero-lag correlation *σ*(0) given the exact expression of 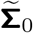 (Equation S130) for computing *q*^*^ for standard SR.

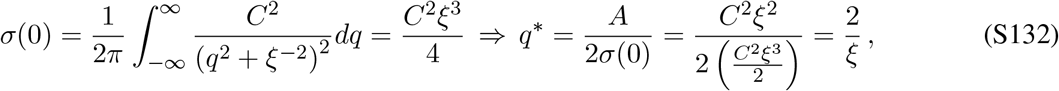

Hence we have that 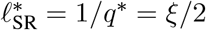. Therefore, an atom’s spatial envelope is bounded above by 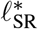 (due to diminishing earnings; Section 3.5.1).

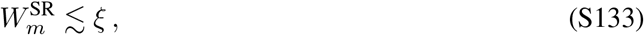

Combining the lower bound on the microscopic resolution with the upper bound spatial extend traps both parameters at *ξ*.

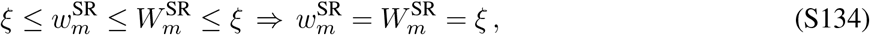

Thus, for standard SR, the octave span 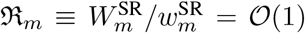 for every atom *m*. Consequently, every surviving atom is forced to exhibit a single, compact, Gaussian-like place field of width *ξ*, without any scattering (duty cycle *η* = 1).

Performing the quasi-NMF optimisation on the standard SR leads to several additional unfavourable consequences. Firstly, under overcomplete decomposition (*M* ≫ *N* ), overlaps between surviving atoms are massive, 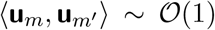, hence the Frobenius norm of the Gram matrix (Section 3.5.2), **G** = **U**^T^**U**, explodes: 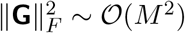. The spiking of 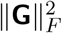 indicates the significant concentration of eigenvalues: a few dominant eigenvalues *λ*_*j*_ spike, while the vast majority collapse to zero. The single largest eigenvalue hence already saturates the trace objective filter *g*(*τ*^2^) (Equation S97), whilst the remaining atoms sit in the saturated regime of *g*(*τ*^2^), contributing marginally to the trace objective whilst inducing additional sparsity penalties. As a result, despite allocating an overcomplete dictionary *M* ≫ *N*, the effective rank of the learned representation collapses to rank(**G**) ∼ *L/ξ* ≪ *M* (effective number of atoms required for tiling up the track).

#### 3.7 Irregular scattering of intra-cell field locations

There is fundamental degeneracy the solution space of the quasi-NMF optimisation with respect to the lobe position. Within a single atom *m*:

- The predictive energy *ρ*_*m*_ ≈ *σ*(0)*m*_*m*_ depends only on its covering mass.
- The sparsity penalty 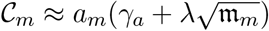 also depends strictly on *m*_*m*_.
- The Gram-conditioning (Equation S111) depends on lobe position only through overlapping with *other* atoms: mutual repulsion between lobes of different atoms, with no constraints for absolute lobe positions within each cell.

Hence, in order to enforce structure (e.g., periodicity) in the spatial arrangement of the intra-cell lobes, an explicit intra-option elastic/harmonic potential term (e.g., *v*(Δ*x*_*m*_) ∝ ∑_*i*_ (Δ*x*_*m,i*_ − Δ*x*_0_)^2^ for Δ*x*_0_ *>* 0, where Δ*x*_*m,i*_ denotes the spacing between the i-th and (i+1)th lobes in atom *m*) needs to be incorporated into the overall objective. No such term exists in the current quasi-NMF optimisation scheme.

Define Ω_periodic_ as the subspace of the space of optimal solutions such that every individual atom has perfectly intra-cell periodic lobe spacings. We further define Ω_disjoint_ as the subspace where lobes interleave such that total overlapping objective 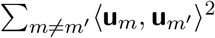 is minimised. Then Ω_periodic_ is of Lebesgue measure zero (lower intrinsic dimension), whereas Ω_disjoint_ contains a vast continuous manifold within the solution space. Hence, the probability that the optimisation process finds a periodic solution is almost surely 0, and we do not expect generic runs to find them.

### 4 Field count distribution

Given the spatial tuning curve (firing rate as a function of spatial location; modelled by **a**_*m*_**u**_*m*_), *f* (*x*), a firing rate threshold, *θ*_*c*_, a discrete “firing field” or a lobe corresponds to a supra-threshold excursion set, an interval [*x*_1_, *x*_2_] where *f* (*x*) ≥ *θ*_*c*_, ∀*x* ∈ [*x*_1_, *x*_2_]. We approximate the number of firing fields as the number of upward level-crossings, *N* ^up^(*θ*_*c*_), where *f* (*x*) crosses *θ*_*c*_ with positive derivative *f* ^′^(*x*) *>* 0. Hence, the expected number of firing fields, *λ*_*m*_ equals the expected number of upward crossings.

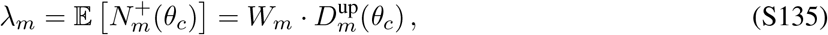

where 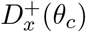 is the spatial density of upward crossings of *θ*_*c*_. As it turns out, 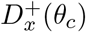 can be analytically computed for a general random function *f* (·) sampled from a Gaussian process with stationary kernel function ^11,12,13^ (see also Mainali et al. 2025^14^).

#### Theorem 4.1

(Kac-Rice Formula). *For a random function f* (·) ∼ *GP* (0, *k*(·, ·)) *with stationary kernel function, k*(*x, y*) = *k*(*x* − *y*), *variance s*_0_ ≡ *E*[ *f* (*x*)^2^], *and derivative variance* 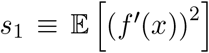. *The expected number of upcrossings* 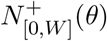 *of level θ within an interval* [0, *W* ], *given by* 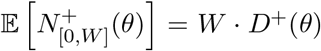, *where the mean upcrossing density per unit length D*^+^(*θ*) *can be expressed analytically as following*.

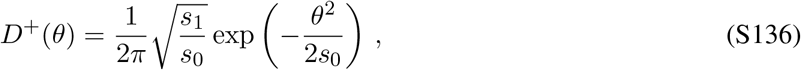

*Proof*. At an upcrossing at *x*_0_, we have *f* (*x*_0_) = *θ* and *f* ^′^(*x*_0_) *>* 0. We then apply first-order Taylor expansion for *x* around the crossing point *x*_0_.

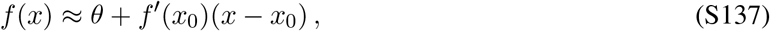

We leverage the composition rule for Dirac’s Delta function *δ*(·), i.e., for any continuously differentiable function g(x) with simple roots *x*_*i*_ (where *g*(*x*_*i*_) = 0 and *g*^′^(*x*_*i*_)/= 0),

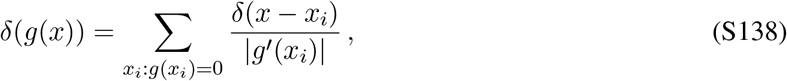

Let *g*(*x*) = *f* (*x*) − *θ*, we have that

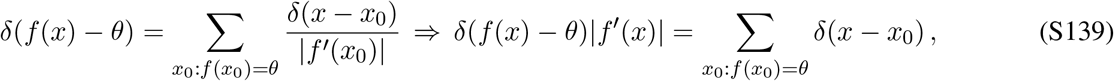

Note that the number of crossings of *θ* equals to integrating the RHS of the above equation over *x* ∈ [0, *W* ].

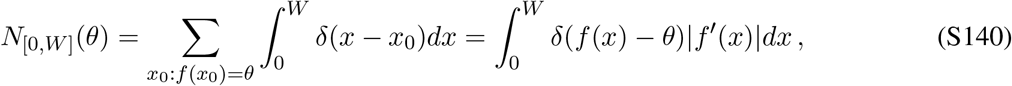

In order to isolate the upcrossings 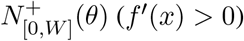, replace |*f* ^′^(*x*)| with max (0, *f* ^′^(*x*)) in the integrand.

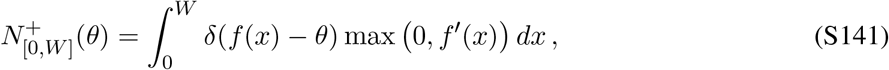

We then take the expectation, and apply Fubini’s theorem to interchange the expectation and spatial integral.

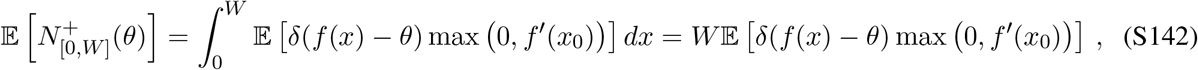

The second equality holds given the stationarity of *f* (*x*), hence the integrand does not depend on the specific spatial location of *x* and is constant over [0, *W* ]. The upcrossing density then becomes

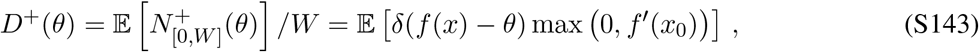

Let *p*(*y, v*) be the joint probability density function of the random vector [*f* (*x*), *f* ^′^(*x*)], where *y* = *f* (*x*) and *v* = *f* ^′^(*x*). We can then express the upcrossing density as the following double integral.

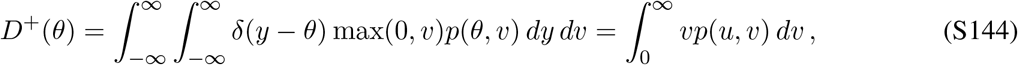

We then note that *f* (*x*) and *f* ^′^(*x*) are independent variables given the stationarity of the random process.

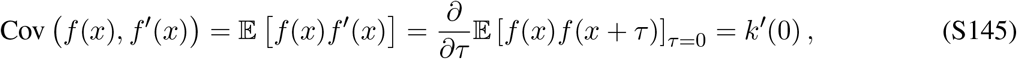

Given the stationarity of *k*(·), we know that *k*(·) is symmetric, hence proving the vanishing of its derivative at 0, and hence the independence between *f* (*x*) and *f* ^′^(*x*).

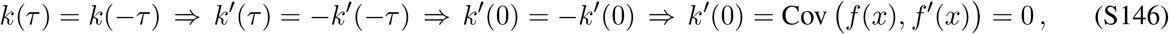

The joint density decomposes into the product of their respective Gaussian densities.

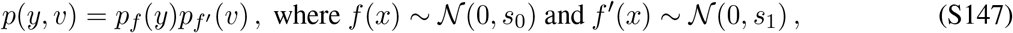

with variance *s*_0_ ≡ *E* [*f* (*x*)^2^], and derivative variance *s*_1_ ≡ *E* [(*f* ^′^(*x*))^2^]. We can hence evaluate the integral to compute the value of the upcrossing ratio.

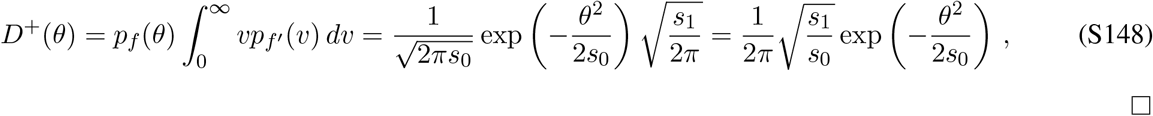

To evaluate *D*^+^(*θ*), we need to evaluate *s*_0_ = *E* [*f* (*x*)*f* (*x*)] = *k*(0) and *s*_1_ = *E* [*f* ^′^(*x*)*f* ^′^(*x*)]. We note that there is a deeper connection between both *s*_0_ and *s*_1_, and the power spectrum of the frame. Define *m*_*n*_ as the n-th spectral moment of the power spectrum of the kernel *k*(*τ* ).

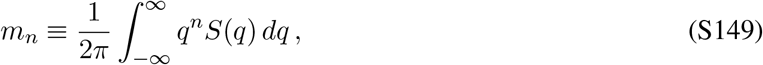

where 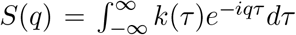is the power spectral density of *k*(*τ* ). We have the following equivalence between the (derivative) variance and spectral moments.

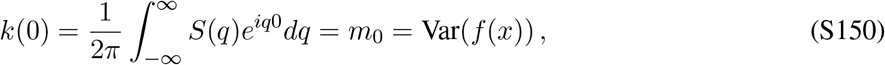

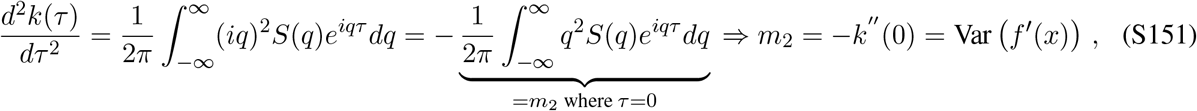

In Section 3.5, we showed that the frame’s power spectrum is half-whitening.

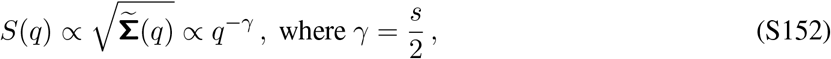

where *s* is the predictive similarity kernel’s spectral slope (*s* = 2 for HSR and *s* = 4 for standard SR; Equations S77 S10).

- **HSR**. *s* = 2, *γ* = 1, *S*(*q*) ∝ *q*^−1^ (Equation S37). The 0-th spectral moment (recall that 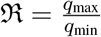; Equation S127).

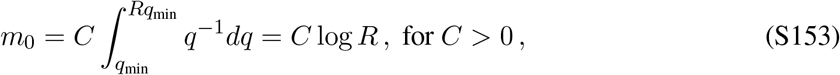

The second spectral moment (for multi-scale atoms).

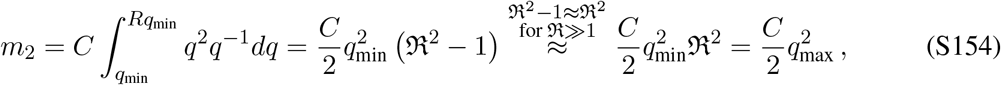

Substituting these into the Kac-Rice formula (Equation S136) for atom *m*.

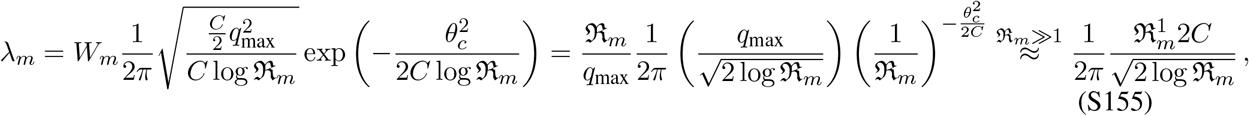

Ignoring the 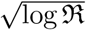-scaling relative to the power-law scaling, the expected field count obeys power-law scaling with respect to the octave span ℜ.

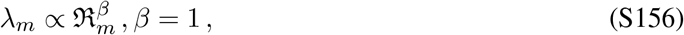
- **General spectral slope** *s*. For analysing what happens for general *s*, we ignore the exponential term in the Kac-Rice formula (Equation S136), since *θ* is constant, and 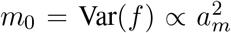, implying the exponential term 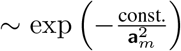. We hence only focus on the 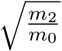 term, and we begin by noting its equivalence to the second moment under the power spectral density. Define 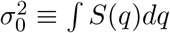 and noting that 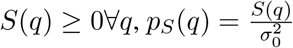 properly defines the spectral density function.

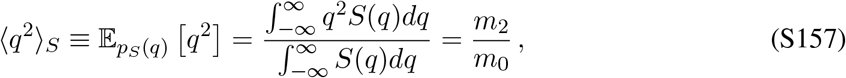 Hence for general *S*(*q*) ∝ *q*^−*s*^, the expectation can be evaluated analytically.

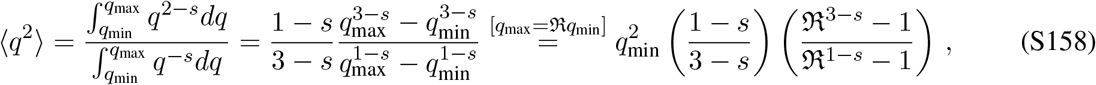

Since *W*_*m*_ ∼ 1*/q*_min_, we have

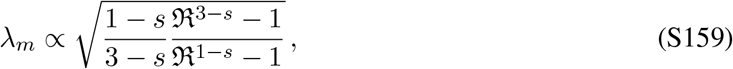
  - *s* < 1. As ℜ_*m*_ ≫ 1 for multi-scale atoms, 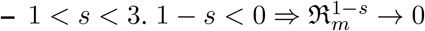.
  - 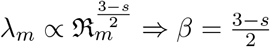. Hence 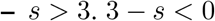.
  - *s* > 3.3 – *s* < 0 and 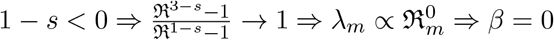
  In summary, we have that

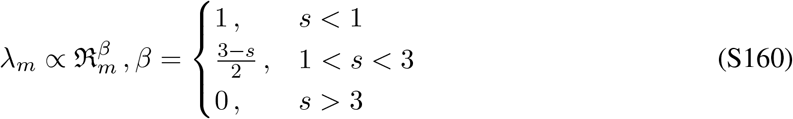

The expected field count log *λ*_*m*_ = *β* log ℜ_*m*_ + const.. For HSR, *β* = 0.5 (*s* = 2) (cf. Equation S156), whereas for standard SR, *β* = 0(*s* = 4 *>* 3), which provides an alternative interpretation of why standard SR does not yield multi-scale spatial representations despite similar quasi-NMF optimisation.

Moreover, we can draw the connection between the magnitude of duty cycles and admissible octave spans within single cells. Specifically, we note that 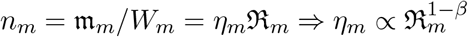. For HSR, this means 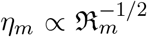 at *s* = 2. This hence predicts that the duty cycle falls as octave span grows, which is directly measurable from the tuning curves.

#### 4.1 Single-cell field counts are conditionally Poisson

For a single atom *m* with mean upcrossing expectation *λ*_*m*_ *E* ≡ [*n*_*m*_], we wish to establish that the realised count of supra-threshold place fields *n*_*m*_ on a given track is a discrete random variable following a Poisson distribution. The Poisson point process behaviour within a single cell arises from two complementary mechanisms operating at different spatial scales: Volkonskii-Rozanov asymptotic independence at large spatial lags ^15,16^, and Rényi thinning of frustrated packing at small spatial lags ^17,18^.

- **Long-range spatial independence**. Let *f* (*x*) be the continuous GP tuning field of atom *m*. The place field locations correspond to connected components of the excursion set above the threshold *θ*.

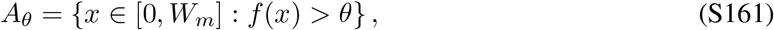 According to the Volkonskii-Rozanov limit theorem ^15,16^, for a continuous and stationary Gaussian process *f* (*x*), as the threshold increases (*θ* → ∞), the point process formed by its upcrossings converges weakly to a spatial Poisson process if the normalised autocorrelation function 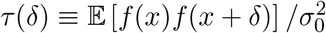 satisfies the Berman’s condition:

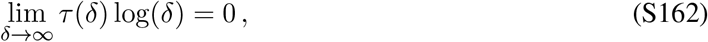

Under the HSR transition kernel, *τ*_HSR_(*δ*) ∼ *δ*^−2^ (Equation S42) for large *δ*.

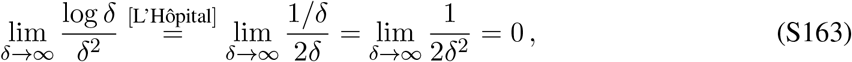

The algebraic decay satisfies Berman’s condition, excursion events (place fields) occurring at spatially separated regions of the track become asymptotically independent, guaranteeing Poisson statistics across macro-scale distances ^16^.
- **The local repulsion**. From Section 3.7, we see that the inter-cell repulsion conditioning (minimizing 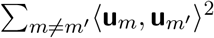; Equation S111) generates a frustrated packing configuration (with respect to the overcomplete decomposition) of candidate lobe positions. This creates an effective (“hard-core” ^18^) spatial exclusion zone of radius *r*_repulsion_ ∼ *w*_*m*_ around each candidate field centre.

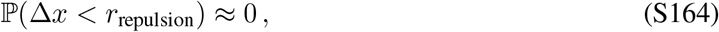

where Δ*x* denotes the spatial gap between adjacent field centres. The minimum gap constraint induces weak spatial regularities, hence locally, the point process is strictly under-dispersed (Var [*n*] *< E* [*n*]). Let X be the spatial point process on ℝ exhibiting the hard-core spatial repulsion with exclusion radius *r*_repulsion_. Suppose each candidate field peak is independently retained with probability *p* = ℙ(*f* (*x*) *> θ*), and deleted with probability 1 − *p*. As the *θ* increases, the detection probability drops (*p* ≪ 1). The average distance between surviving fields expands.

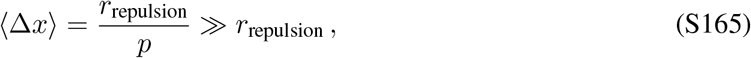

We then leverage the following theorem ^17^

#### Theorem 4.2

(Rényi, 1956). *Let X be a stationary point process on* ℝ *with finite intensity λ*_0_ *>* 0 *and i*.*i*.*d. inter-event gaps T*_*i*_ *(a renewal process). Suppose the process is subjected to independent Bernoulli thinning with retention probability p* ∈ (0, 1), *which retains each point with probability p and deleting it with probability* 1 − *p. If the spatial axis is rescaled by y* = *p* · *x, then as p* → 0, *the thinned and rescaled point process X*_*p*_(*y*) *converges in distribution to a homogeneous Poisson point process with intensity λ* = *λ*_0_.

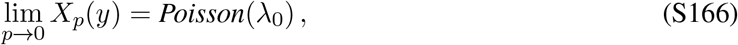

*Proof*. In the unthinned process *X*(*x*), let *T*_1_, *T*_2_, *T*_3_, … be the i.i.d. spatial gaps between consecutive candidate events, with mean gap *E*[*T* ] ≡ *µ* = 1*/λ*_0_. When each event is independently kept with probability *p*, the number of unthinned lobes *G* skipped before the first retained lobe follows a geometric distribution.

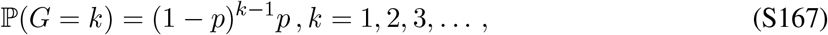 The spatial gap *S*_*G*_ between two retained events on the original coordinate *x* is the sum of *G* independent candidate gap variables.

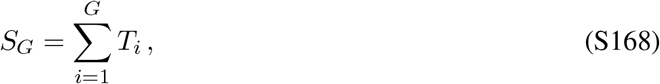

Applying the spatial rescaling, 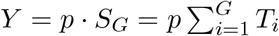, and compute the Laplace transform *ϕ*_*Y*_ (*s*) ≡ *E*[*e*^−*sY*^].

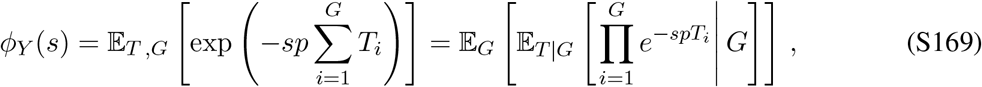

Since *T*_*i*_’s are i.i.d. with Laplace transform *ϕ*_*T*_ (*s*) ≡ E *e*^−*sT*^, we can express *ϕ*_*Y*_ (*s*) in terms of *ϕ*_*T*_ (*s*).

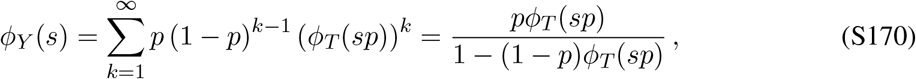

Applying first-order Taylor expansion of *ϕ*_*T*_ (*sp*) around *p* = 0, and using 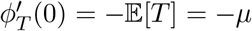.

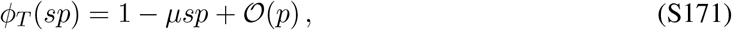

Substituting this Taylor expansion back into *ϕ*_*Y*_ (*s*) and take the limit *p* → 0.

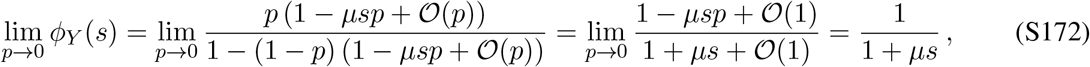

which is exactly the Laplace transform of an exponential distribution with rate parameter *λ* = 1*/µ*. Such exponential inter-event interval uniquely defines a homogeneous Poisson point process, hence random thinning converts any finite-mean interval distribution into a memoryless exponential-interval distribution. Hence by Rényi’s theorem, as *θ* → ∞ and *p* → 0, the thinned point process *X*_*p*_(*y*) converges in distribution to a homogeneous Poisson process, with intensity *λ*_*m*_.

In conclusion, at both the global and local scale, the stochastic event of upcrossings that define the place fields converge towards Poisson independence, hence conditioning on its mean rate *λ*_*m*_, the field count *n*_*m*_ within any individual cell *m* is asymptotically Poisson distributed.

#### 4.2 Expected number of fields is Gamma-distributed across cells

Having established the intra-cell field distribution is conditionally Poisson, we now address the second part of the experimental findings ^19,20^: across cells, *λ* follows a Gamma distribution. Below we provide two complementary approaches that arrive at the same distributional form.

- **Multiplicative cascade**. We established in Section 2.3.1 that the HSR power spectrum distributes frame energy equally across arbitrary spatial octaves. Hence, when an overcomplete dictionary learns a multiscale representation, each atom *m* with octave span ℜ_*m*_ requires making a series of independent inclusion choices across log_2_ ℜ_max_ available spatial octaves. We can formulate this inclusion process as following.

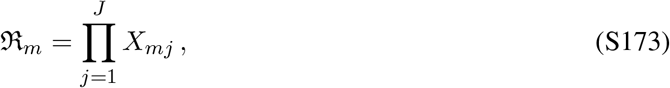

where *X*_*mj*_ *>* 1 represents the expansion factor contributed by including the j-th octave band. Taking the natural logarithm of ℜ_*m*_, log ℜ_*m*_ = ∑_*j*_ log *X*_*mj*_, then by the central limit theorem applied to sums of independent log variables, in the infinite limit of number of available octaves, log ℜ_*m*_ converges to a Gaussian distribution.

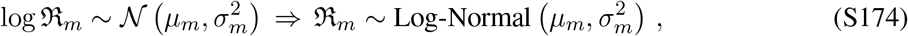

We previously established that the expected field count *λ*_*m*_ scales with the atom’s octave span, 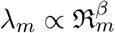 (Equation S160). Hence log *λ*_*m*_ also follows a Gaussian distribution, and the raw counts *λ*_*m*_ follows a log-normal distribution.

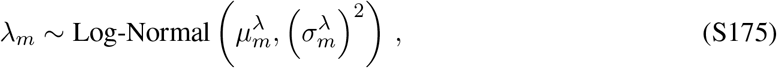

Note that a log-normal distribution and a Gamma distribution with matched first and second moments are virtually indistinguishable across moderate variance regimes 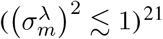.

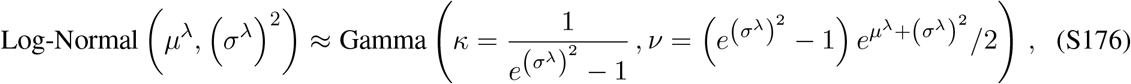

The equal spread of power across multiple spatial hierarchies naturally manifests into the heavy-tailed rate distributions across the dictionary.
- **Maximum entropy principle**. The maximum entropy principle drives the emergence of Gamma-distributed *p*(*λ*) without assuming a specific ladder-generating process. In addition to maximising the entropy, there are several (implicit) constraints on *p*(*λ*).
  - To represent a one-dimensional lattice with *N* sites without leaving unrepresented “blind spots”, the *M* atoms must completely tile the two-dimensional phase-space (*x, q*). Hence the sum of phase-space volumes (Equation S127) across all atoms must satisfy a lower bound proportional to the lattice size *N*.

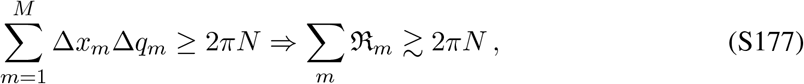 We previously established the expected number of fields is directly proportional to its spatial octave span (Equation S160). The mean rate is hence proportional to the sum of octave span, and to the sum of phase-space volumes across all neurons. Therefore, the mean rate is finite and should be constrained to be some non-zero value that is proportional to the environment size.
  - Under the HSR, we previously established that its power spectrum 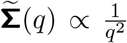 attributes equal energy to every spatial octave. An atom *m* spanning multiple octaves builds its octave span ℜ_*m*_ = *q*_max,*m*_*/q*_min,*m*_, and given the relationship between field count rate and the octave span (Equation S160) in the log-space, we have that log *λ*_*m*_ = *β* log ℜ_*m*_ + const.. Thus, requiring the dictionary to be scale-invariant means that the expected value of log ℜ_*m*_, and therefore log *λ*_*m*_, is a fixed and finite constant across the population.

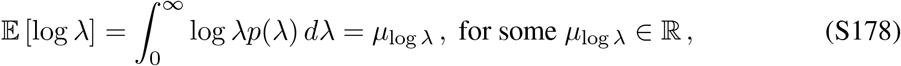
  Combining the above two physical constraints with an additional normalisation constraint ∫ *p*(*λ*) = 1, the free-form optimisation problem is as following.

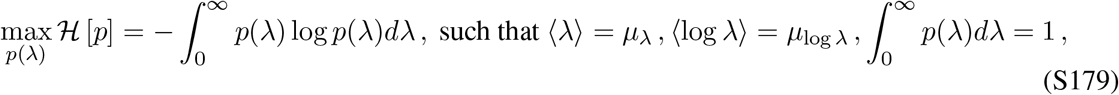

The functional Lagrangian objective, with Lagrange multipliers *α*_0_, *α*_1_, *α*_2_ is as following.

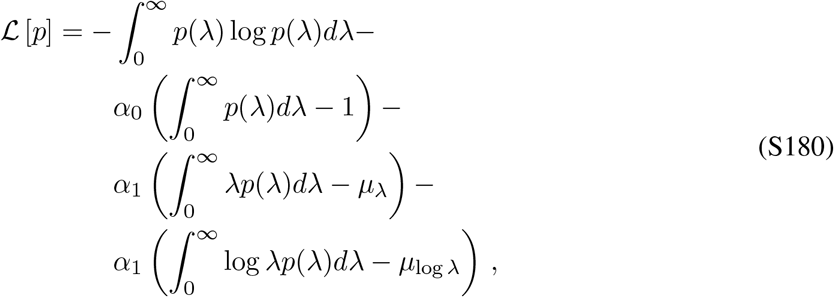

Taking the variational derivative with respect to *p*(*λ*) and set to zero.

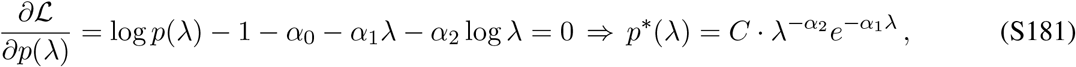

Matching the terms with a Gamma distribution, we see that *p*^*^(*λ*) follows a Gamma distribution.

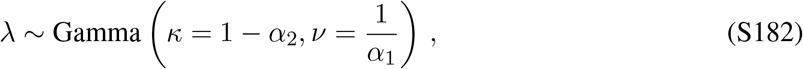

A direct consequence of the max-entropy-derived functional form of *p*(*λ*) is that when the model operates over only one single octave (single-scale code; e.g., standard SR), the log-constraint ⟨log *λ*⟩ becomes completely redundant with the mean constraint ⟨*λ*⟩. In this case, taking the limit *κ* → ∞ and *v* → 0 whilst holding the mean *µ* = *κv* = ⟨*λ*⟩ fixed, the Gamma distribution collapses into a Dirac delta function, forcing *λ*_*m*_ to be constant for all cells.

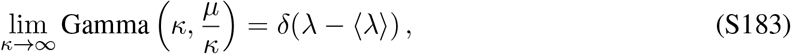

#### 4.3 A partial spectral estimate of Gamma-distributed propensity

By definition, **Σ** is symmetric and positive semi-definite, hence **Σ** admits an orthonormal eigendecomposition.

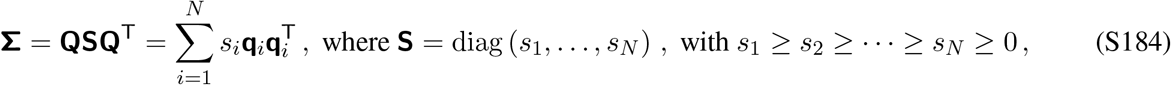

where **q**_*i*_’s are orthonormal spatial eigenbasis.

For atom *m*, the predictive energy given the above eigendecomposition takes the following quadratic form.

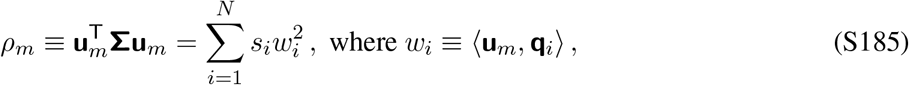

To compute the population ensemble statistics over predictive energies across generic unit vectors **u**_*m*_, we model **u**_*m*_ as random vectors uniformly distributed over the unit hypersphere *S*^*N* −1^ ⊂ ℝ^*N*^. We wish to note that this is not a physically valid assumption for the “place cells” discovered through the quasi-NMF decomposition. Instead, it serves purely as an isotropic null model, a mathematical trick adopted from random matrix theory used to analytically evaluate the moments over *ρ* in closed form using spherical tensor moments. Mathematically, these random vectors can be treated as normalised Gaussian spatial white noise.

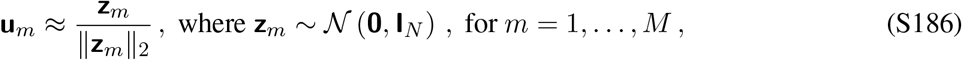

In high dimensions (*N* ≫ 1), this means that at every spatial coordinate **u**_*m*_(*x*) is independently drawn from a zero-mean Gaussian distribution with variance 1*/N*, **u**_*m*_(*x*) ∼ *N* (0, 1*/N* ).

We can then compute moments of predictive energy *ρ*. We start with the mean energy.

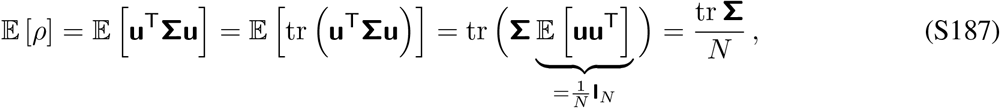

To compute its variance, we first evaluate the second moment *E* [*ρ*^2^].

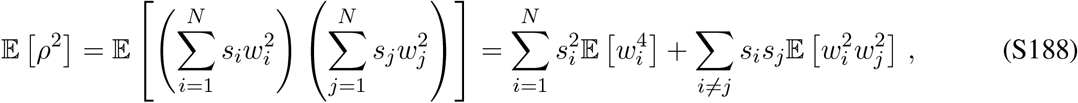

Since **w** = (*w*_1_, …, *w*_*N*_ )^T^ is a projection of eigenvectors onto random vectors over *S*^*N* −1^, **w** itself is a random vector that is uniformly distributed over *S*^*N* −1^. We can hence analytically evaluate its fourth-order coordinate moments.

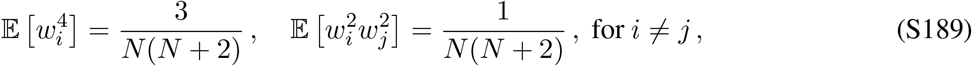

We first note that the (tr **Σ**)^2^ decomposes into diagonal and off-diagonal terms: 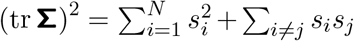. We then substitute these back in the definition of the second moment,

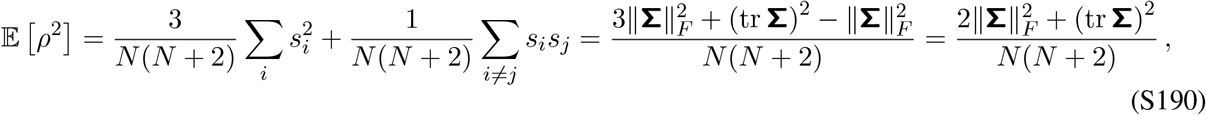

We can then compute the variance.

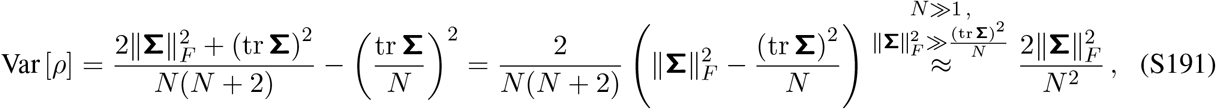

To approximate the distribution of the atom-specific energy *ρ* with a Gamma distribution 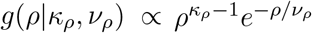, we match the first two moments.

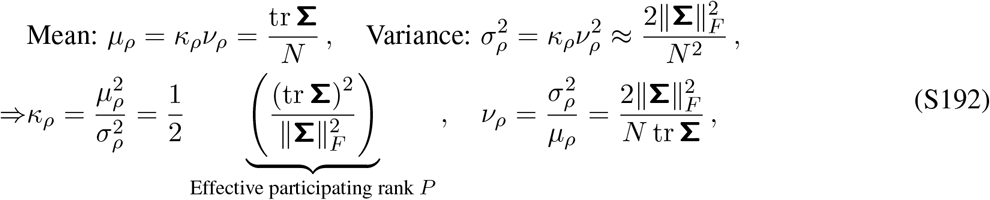

Note that the Gamma-distributed predictive energy is related to the field count through a shared equivalence to the atom’s support mass *m*_*m*_, which is the summarised in the diagram below.

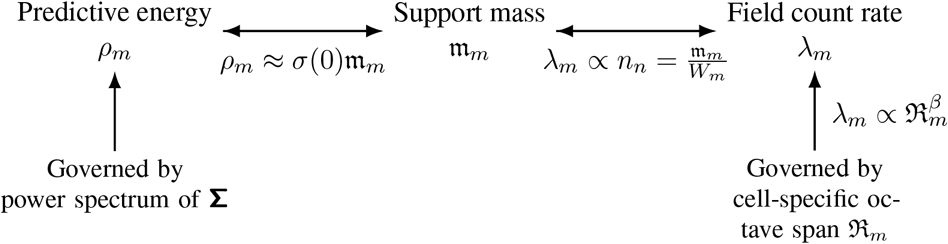

### 5 Multi-scale representations and spatial decoding accuracy

#### 5.1 Local precision

For conditionally independent Poisson spikings with tuning curves from *M* neurons, 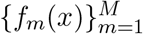, and temporal bin size *τ*, the Fisher information at each specific spatial location is as following.

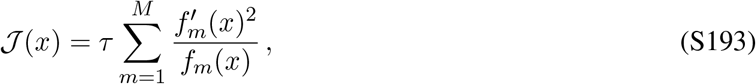

The local decoding error around the ground-truth location is governed by the Cramér-Rao lower bound, with its variance bounded below by the inverse of the population Fisher information *J* (*x*). For a spatial field/lobe of width *w*_*m*_, the spatial gradient of the firing scales 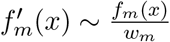, where we have assumed Gaussian firing fields. Hence, the population-level Fisher information can be re-expressed as following.

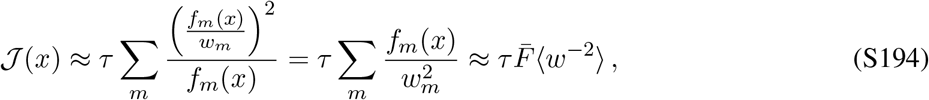

where 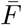 and ⟨*w*^−2^⟩ denote the peak firing rate and the average of 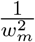 across all place fields and all neurons, respectively. Due to the inverse quadratic scaling, ⟨*w*^−2^⟩ is dominated almost entirely by the narrowest fields across all neurons (*w*_min_). Hence, the local decoding precision scales as

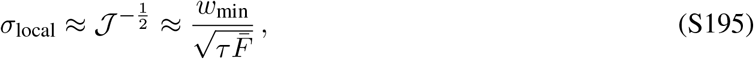

In short, the local resolution of spatial decoding is set linearly by the narrowest field width.

#### 5.2 Global aliasing

Local Fisher Information measures decoding error only over a spatial resolution of a single firing field, and it cannot account for *global aliasing* effect, where the decoder confuses true position *x* with a distant candidate position *x*^′^ that happens to share co-active neurons.

We define the cell-specific active fraction, *h*_*m*_, as the fraction of the track length *L* over which neuron *m* is “ON” (covered by active firing fields).

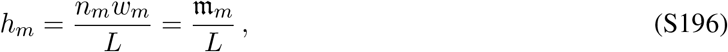

At location *x*, let *S*_*x*_ = {*m* : *x* ∈ fields_*m*_} be the set of active neurons. A candidate position *x*^′^ survives as a false alias if *x*^′^ falls inside a field of every cell *m* ∈ *S*_*x*_. Assume mutual independence across neurons, the survival probability is 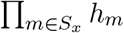, and the expected log-survival rate of a candidate false location is the sum of cell-specific negative entropies.

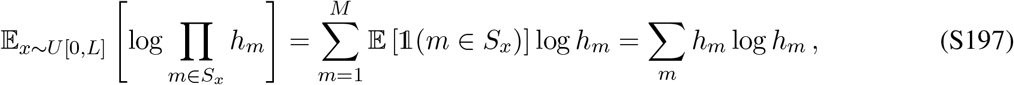

Hence the expected number of aliases is

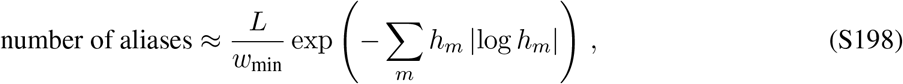

The per-cell contribution −*h* log *h* is maximised at 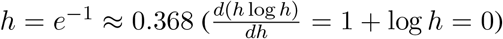. Cells with *h*_*m*_ ≪ *e*^−1^ fires too sparsely to contribute to the disambiguation sum, whereas cells with *h*_*m*_ ≫ *e*^−1^ fire almost everywhere, hence providing little spatial specificity. In contrast, cells with *h*_*m*_ ∼ *e*^−1^. maximise their spatial coding entropy and provide optimal global aliasing suppression per neuron.

We then investigate implications from the single- and multi-scale representations on spatial decoding separately.

- **Single scale**. Consider a single-scale frame consists of *M* neurons, where each neuron *m* has the exact same field width *w* and a single firing field *n*_*m*_ = 1. Hence *h*_*m*_ = *w/L* for all *m*. We additionally define the redundancy ratio as the ratio of total number of cells to the number of resolution bin.

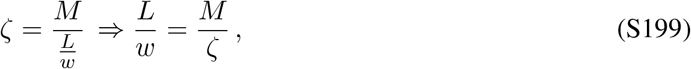 We then evaluate the expected log-survival rate (Equation S197) across the *M* single-scale neurons.

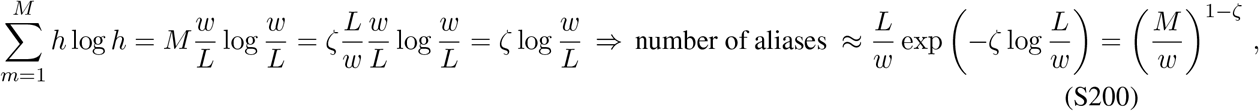

At critical sampling 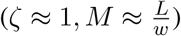, the decoder is plagued by ≈ 1 persistent global aliases somewhere along the track. In order to reduce aliasing, we can increase the redundancy ratio (*ζ* ≫ 1), such that the aliasing suppression improves only as a power law of cell count *M* .
- **Multi-scale**. Consider a multi-scale code combining fine-scale fields of widths *w*_min_ with a pool of *M*_coarse_ coarse fields whose spatial coverage is tuned to near entropy-maximising optimum (*h*_coarse_ = *e*^−1^ ≈ 0.368). We now split the computation of the expected log-survival rate into separate fine and coarse scales.

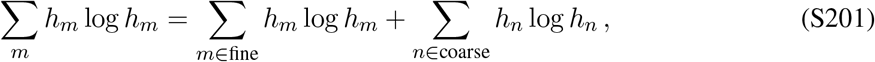 Assume *ζ*_fine_ = 1 (⇒ *M*_fine_ = *L/w*_min_), we hence have

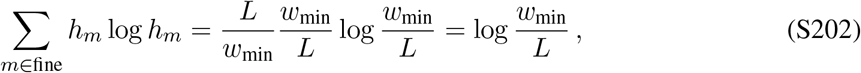

Each coarse-scale neuron contribute near *h*_*n*_ ≈ *e*^−1^ to the entropy sum.

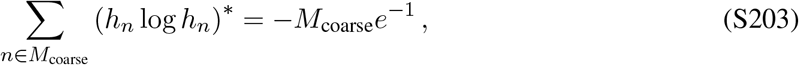

Thus, the expected number of aliases is as following.

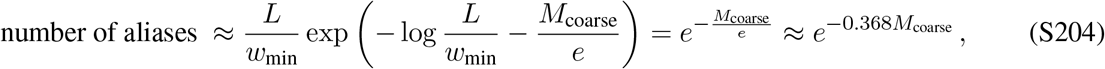

We hence have shown that the single-scale code admits polynomial suppression in global aliasing, whereas multi-scale code yields exponential suppression. Thus, in terms of coding efficiency, the multi-scale code requires significantly less number of neurons to reach the same level of global aliasing. For example, for *M* = 1000, to reach negligible aliasing probability of *ϵ* ∼ 10^−6^, the single-scale would require 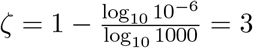, hence requiring a three-fold increase in the entire neural population. In contrast, the multi-scale coding would only require −log 10^−6^· *e* ≈ 38 corase neurons. Conversely, under the same number of neurons, the multi-scale coding would lead to significantly lower probability for global aliasing.

### 6 HSR and hyperbolic geometry

The spatial atoms discovered through the quasi-NMF decomposition of the HSR are naturally described by translations and changes of spatial scale. We show that the orbit generated by these two transformations is diffeomorphic to the orientation-preserving affine group and that the canonical affine-invariant Riemannian metric on this group is precisely the Poincaré metric that defines the hyperbolic geometry. We note that our argument makes no assumption of the Gaussianity or uni-modality about the shape of the atoms.

Let *u*_0_ : ℝ → ℝ_≥0_ be the spatial function prescribed by a non-negative, localised reference atom with finite, non-zero spatial variance. A translation *b* ∈ ℝ and dilation *c* ∈ ℝ_*>*0_ act on *u*_0_ according to

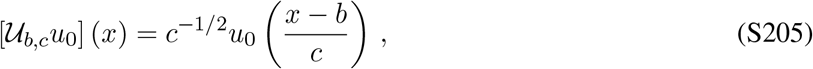

Note that the factor *c*^−1*/*2^ ensures that the *L*^2^ norm is preserved: 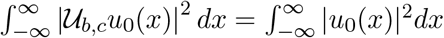. Thus translations and dilations alter the location and scale of an atom without changing its normalised shape or gain.

To define the location and scale of a non-negative atom **u**, we first construct the corresponding normalised spatial firing profile, 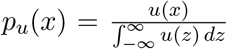 . The corresponding spatial centre and root-mean-square scale are 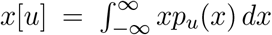 and 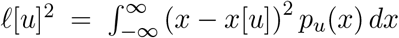, respectively. On the discrete lattice, the corresponding definitions are as following.

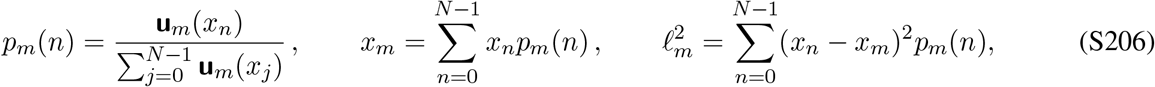

where *x*_*n*_ = *ns* is the physical coordinate of the n-th lattice state.

Under the affine transformation (Equation S205), we have the following transformed normalised spatial profile and corresponding translated-and-scaled coordinates.

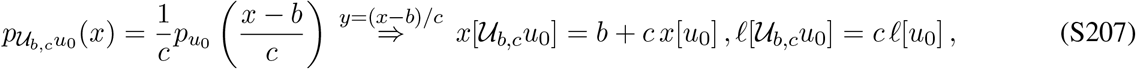

Without loss of generality, choose the reference atom to have *x*[*u*_0_] = 0 and *l*[*u*_0_] = 1. The transformed atom then has coordinates *x*[ *U*_*b,c*_*u*_0_] = *b* and *l*[*U*_*b,c*_*u*_0_] = *c*. The group parameters are therefore exactly the spatial position and scale of the transformed atom.

The set of orientation-preserving translations and dilations is the affine group

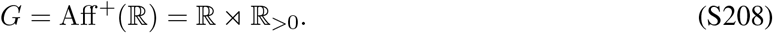

Composition of two transformations gives 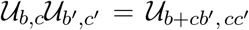, we can hence define the group action as following.

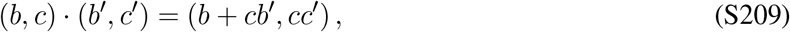

The identity is (0, 1) and the inverse 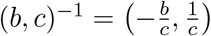, for all (*b, c*) ∈ *G* (since *c >* 0).

Consider the orbit of the reference atom, 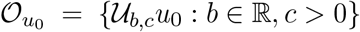, and its stabiliser, Stab(*u*_0_) = {(*b, c*) *G* :*U*_*b,c*_*u*_0_ = *u*_0_ }. By the Orbit-Stabiliser theorem for Lie groups, we have that 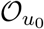 is isomorphic to the quotient space *G/* Stab(*u*_0_).

For a localised atom with finite, non-zero spatial variance, its stabiliser group is trivial. To see this, suppose _*b,c*_*u*_0_ = *u*_0_. Then the affine transformation (Equation S207; S209) requires *l*[*u*_0_] = *c l*[*u*_0_]. Since *l*[*u*_0_] *>* 0, this implies *c* = 1. Equality of the centres then requires *x*[*u*_0_] = *b* + *x*[*u*_0_], which implies *b* = 0. Therefore, the stabiliser group only contains the identity element, and 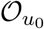 is isomorphic as a G-manifold to *G*.

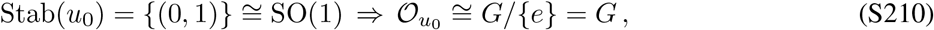

Indeed, the map Ψ : *G*→ 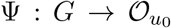, where Ψ(*b, c*) = *U*_*b,c*_*u*_0_, is a smooth bijection whose inverse is supplied by the centre and scale coordinates (Equation S207). The atom orbit is therefore diffeomorphic, not merely homeomorphic, to the affine-group manifold.

We next compute the Riemannian metric on *G*. At the identity (0, 1), assign the Euclidean inner product as following.

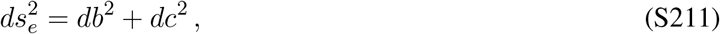

Left multiplication by (*b, c*) maps a group element (*b*^′^, *c*^′^) to *L*(*b,c*)(*b*^′^, *c*^′^) = (*b*+*cb*^′^, *cc*^′^), hence the corresponding differential maps a tangent vector (*db*^′^, *dc*^′^) at the identity to *dL*(*b,c*)(*db*^′^, *dc*^′^) = (*c db*^′^, *c dc*^′^). Requiring the tangent-vector norm to remain unchanged under this left action therefore entails 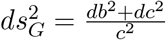 . The *ds*^2^ on *G* (Equation S211) is thus invariant under the translation-dilation group.

Identifying *b* with the horizontal coordinate *x* and *c* with the positive scale coordinate *l* that constitutes the parameter groups for the spatial atoms, *G*, we have the Riemannian metric over *G* as following.

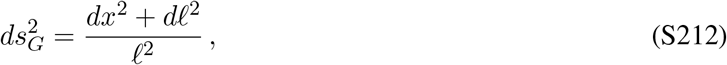

This is exactly the canonical Poincaré metric of the upper-half-plane ℍ ^222^.

The curvature of this metric can be evaluated directly. We define the logarithmic scale coordinate, *y* = log *l*. Since *dl* = *l dy*, the metric tensor (Equation S212) then becomes 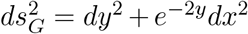. For a two-dimensional warped metric of the form *ds*^2^ = *dy*^2^ + *f* (*y*)^2^*dx*^2^, the Gaussian curvature is 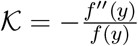 . Here *f* (*y*) = *e*^−*y*^ and *f* ^′′^(*y*) = *e*^−*y*^, hence *K*_*G*_ = −*e*^−*y*^*/e*^−*y*^ = −1. Therefore, 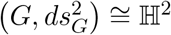 as Riemannian manifolds. Combining this isometry with the diffeomorphism between 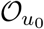 and *G* (Equation S210) gives

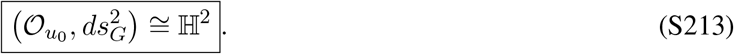

Thus, the position-scale orbit of any localised spatial atom is a hyperbolic manifold. The result depends only on the translation–dilation action and not on whether the atom is Gaussian, unimodal, compact, or multi-lobed.

For a general *d*-dimensional physical environment, the same construction uses **b** ∈ ℝ ^*d*^ and *c >* 0:

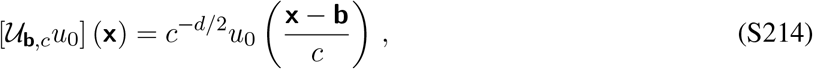

The corresponding invariant metric tensor is

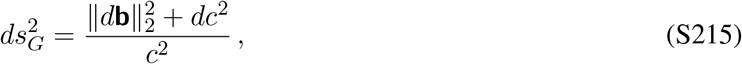

which is the metric of H^*d*+1^. Hence a one-dimensional physical environment generates a two-dimensional hyperbolic position–scale manifold, whereas a two-dimensional environment generates H^3^.

The conclusion is exact for atoms related by translations and isotropic dilations of a common reference shape. If learned atoms contain additional shape variation that cannot be represented by (*b, c*), then the full atom space is not a single copy of H^2^. Instead, it is a union of hyperbolic group orbits indexed by the remaining shape parameters. The accuracy of the single-orbit description can be quantified without assuming a particular parametric atom shape by measuring the minimal deviation.

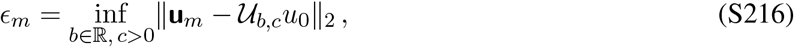

Small residuals *ϵ*_*m*_ would establish that the learned population lies close to a common hyperbolic position–scale manifold. On a finite lattice, the allowed position and scale ranges are bounded, so the realised representation occupies a truncated region of H^2^; reflecting boundaries and lattice discretisation introduce additional deviations near the edges.

### 7 Population-level spatial oscillations

#### 7.1 The eigenvalue-retention criterion fixes an environment-agnostic frequency cutoff scale

For the reflecting one-dimensional random walk, the spatial eigenmodes are standing waves with Neumann correction.

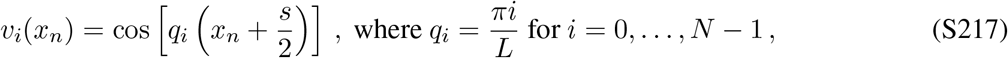

The corresponding eigenvalues of the random-walk SR are

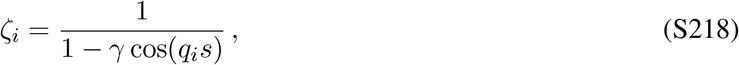

Note that we exclude the constant mode from option construction because it produces flat spatial pseudo-reward. Removing one mode changes the continuum expressions below only by a finite-size correction.

For *q*_*i*_*s* ≪ 1, expanding the denominator with Taylor expansion (Equation S10), we have the following approximation.

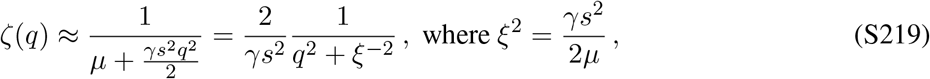

Let *θ*_*f*_ ∈ (0, 1) denote the threshold on cumulative variance explained for selecting the number of eigenoptions (in the simulations, *θ*_*f*_ = 0.95). Note that the highest frequency is the lattice Nyquist wavenumber, *π/s*. We then compute the *K*-th cumulative explained variance ratio as following (replacing the discrete sums by integrals).

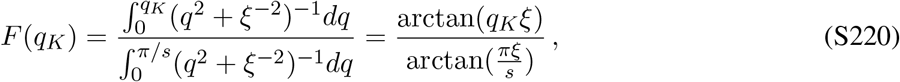

Solving *F* (*q*_*K*_) = *θ*_*f*_ we have that

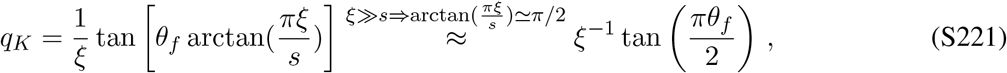

The characteristic scale of the finest retained option, and associated number of retained options can then be computed analytically as following.

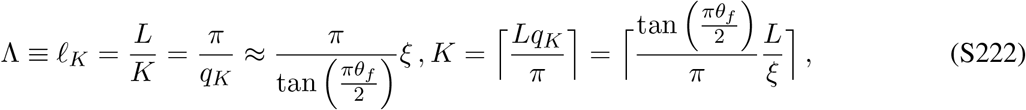

Thus the number of retained options grows approximately linearly with track length, whereas their finest physical scale remains fixed by the SR correlation length and is independent of the track length. Integer rounding of *K*, finite lattice spacing, exclusion of the constant mode, and deviations from the Lorentzian approximation produce finite-size corrections to Equation (S222), but do not change the cancellation of *L*.

#### 7.2 Eigenoptions impose population-level phase coherence

The independence of Λ from track length explains the available spatial scale but not, by itself, why that scale appears in summed population activity. The latter requires coherence across cells.

We define the summed population-level spatial tuning profile as following.

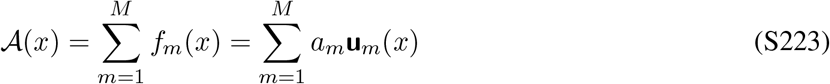

Its Fourier transform follows trivially as 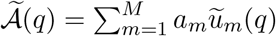, and its power spectrum decomposes exactly into within-cell and cross-cell terms.

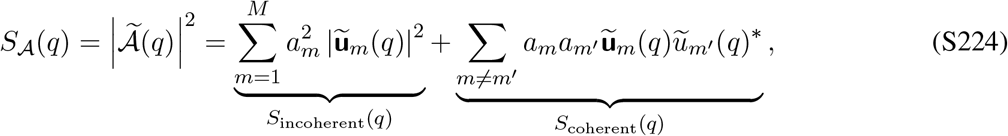

The first term is the incoherent frame power. It depends only on individual cells’ spectral amplitudes and is therefore retained when cells are shuffled independently. The second term depends on relative Fourier phases and vanishes in expectation under independent spatial shuffling.

Although the spatial organisation of lobes of any individual learned atom are irregular because of the frustrated quasi-NMF packing (Section 3.7), their eigenmode components remain referenced to the same boundary-pinned combs. Summation across cells therefore cancels cell-specific irregularities while preserving weak phase-locked modulation at the option scales.

To quantify this effect, we define 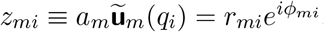, with *r*_*mi*_ = |*z* | and *ϕ* = arg(*z*_*mi*_ ), as the Fourier transform of the atom *m* at frequency *q*_*i*_. We further define the population-level phase-coherence factor at the i-th frequency scale as following.

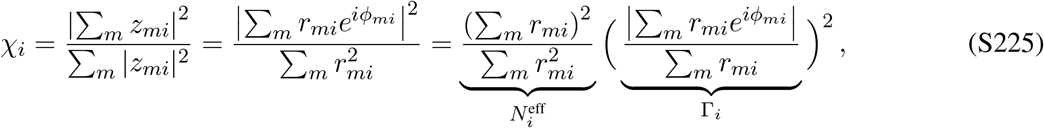

where 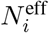 and Γ_*i*_ define the effective number of cells contributing and the weighted phase-locking value at frequency *q*_*i*_, respectively.

If all cells have equal amplitudes, then 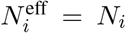 . Perfect phase alignment gives Γ_*i*_ = 1 and therefore 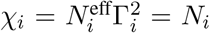 . By contrast, if the phases are independently and uniformly distributed, then 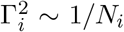 and E [*χ*_*i*_] = 1. Consequently, *χ*_*i*_ scales with cell number only when the population retains a non-vanishing degree of phase-locking between atoms. Simply sharing spatial frequencies is not sufficient. More generally, suppose the phases *ϕ*_*mi*_ are independent draws from a circular distribution with mean resultant vector length (1 - circular variance) 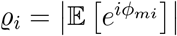, then 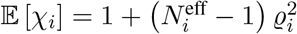 . Hence the independent-phase limit is *ϱ*_*i*_ = 0, whereas perfect phase-locking corresponds to *ϱ*_*i*_ = 1.

We next explain how the HSR can generate a non-zero mean resultant length *ϱ*_*i*_. Importantly, this coherence does not follow merely from cells sharing the same spatial frequency. It arises because all eigenoptions associated with a given mode inherit a common spatial phase from the reflecting boundaries. Recall that the i-th eigenmode of the reflecting random walk are standing waves (Equation S217), which means each extrema (terminal state) occurs at 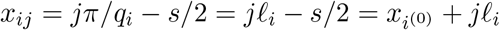, where *l*_*i*_ = *π/q*_*i*_ = *L/i* is the i-th characteristic lengthscale and *x*^(0)^ = *s/*2. Hence, all terminal locations have exactly the same phase: 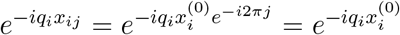.

This common phase is the source of population coherence, which is imposed by the shared optimal terminal states, and is therefore common to all cells that draw substantial predictive structure from option scale *i*.

To allow for irregular field location produced by the quasi-NMF optimisation (Section 3.7), suppose the actual centre of a place field is displaced from its nearest ideal terminal location (*j*) with random spatial jitter.

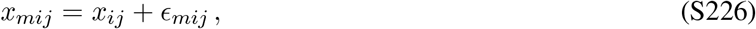

A general atom drawing substantial predictive structure on scale *i* can then be approximated locally as following.

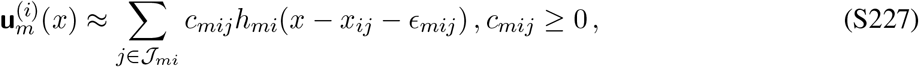

where *h*_*mi*_ is the local spatial response profile around each lobe, and ℐ_*mi*_ denotes the subset of terminal locations represented by atom *m*. Taking the Fourier transform at the *i*-th frequency scale gives

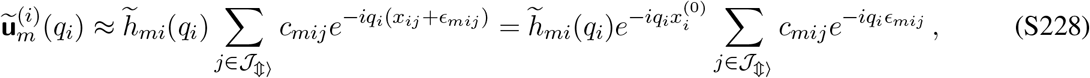

The factor 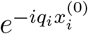 is common to all included cells, hence the cell-to-cell phase differences arise only from the spatial jitters *ϵ*_*mij*_ and from any asymmetry in the local lobe profiles.

Assume the residual jitters are i.i.d., the remaining phase concentration is governed by the characteristic function of the jitter distribution.

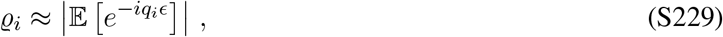

Hence *ϱ*_*i*_ *>* 0 whenever the lobe positions remain statistically concentrated around the shared terminal lattice. Conversely, if the residual offsets become uniform, and the concentration becomes 0, and population phase coherence vanishes.

We assume, for simplicity, the spatial jitter is Gaussian: 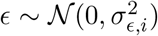, then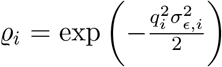 . Thus, the coherence is preserved when *q*_*i*_*σ*_*ϵ,i*_ ≪ 1, meaning that the spatial jitter is small relative to the corresponding terminal spacings. Fine spatial scales are more sensitive to a fixed absolute amount of spatial jitter since their corresponding *q*_*i*_ is larger.

Substituting the Gaussian-coherence into the expected coherence.

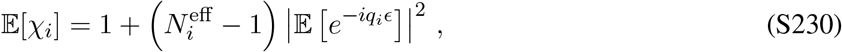

The HSR therefore generates *χ*_*i*_ *>* 1 through two ingredients: multiple cells must draw on the same option scale, giving 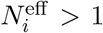, and their lobes must retain some alignment with the shared option-terminal lattice, giving *ϱ*_*i*_ *>* 0.

This mechanism also clarifies what is and is not guaranteed by the model. The reflecting boundaries and common eigenoption library provide a shared phase reference, but the quasi-NMF optimisation breaks perfect phase locking. The magnitude of *ϱ*_*i*_ depends on how strongly the learned atoms inherit the terminal locations relative to the irregularity introduced by the overcomplete packing. Phase coherence is therefore a quantitative prediction to be measured from the learned atoms rather than an exact consequence of half-whitening.

The scaling *χ*_*i*_ ∝*q*_*i*_ requires one additional assumption. Let 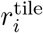denote the redundancy, or effective number of cells allocated per terminal interval, at scale *i*. The effective number of contributing cells is then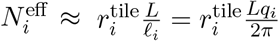. We assume that the redundancy 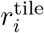and phase concentration *ϱ*_*i*_ are approximately constant across scales, then we have 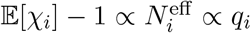. This scaling is therefore contingent on the overcomplete decomposition, rather than from the option frequencies alone.

Finally, substituting the HSR frame spectrum *P*_Φ_(*q*) ∝|*q* |^−1^ and expected coherence (Equation S230) into the power spectrum (Equation (S224)), we have

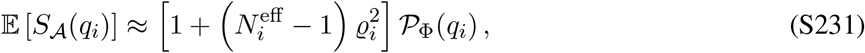

Under the standard assumption of 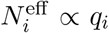 and *ϱ*_*i*_ is scale-independent, the coherent part is approximately constant across the option ladder:

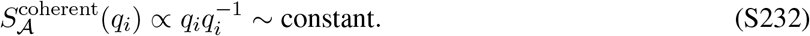

#### 7.3 The first non-zero autocorrelation peak is set by the finest retained comb

By the Wiener–Khinchin theorem, the population autocorrelation is the inverse Fourier transform of power spectrum density.

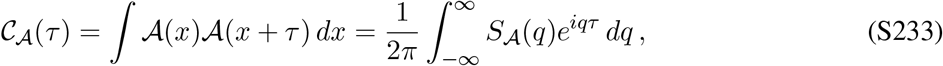

For a population assembled from boundary-locked option combs, this takes the following schematic form.

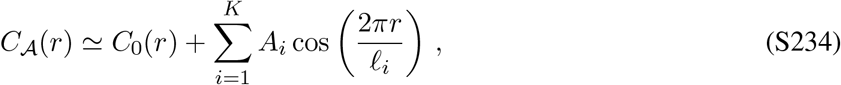

where *C*_0_(*r*) is a smooth background and *A*_*i*_ measures coherence at scale *i*.

The finest retained comb has the shortest period *l*_*K*_ = *L/K*. Provided its coherent amplitude is non-zero, it sets the scale of the first resolvable non-zero autocorrelation peak:

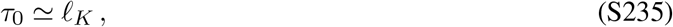

The equality is exact for an isolated ideal comb. In a superposition of scales, coarser components can shift the local maximum by an order-one amount, so the estimate of the first non-zero peak is a scaling prediction rather than an exact identity. Given *l*_*K*_ is independent of track length (Equation S222), we know that *τ*_0_ is also independent of track length.

#### 7.4 Why field shuffling removes the oscillation

Suppose each cell is independently translated by a random displacement *δ*_*m*_, and the associated Fourier transform goes through phase shift accordingly.

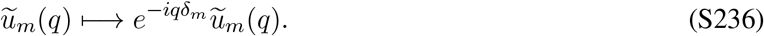

For independent uniform translations, we h ave that

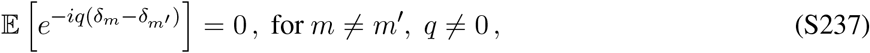

Consequently, the expected phase coherence under the multiplicative factor of the random shuffling becomes zero, and the power spectrum is solely dependent on the incoherent half-whitening power spectrum of the frame, which provides a random baseline due to the frustrated packing.

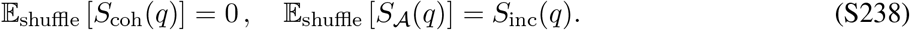

This explains why shuffling procedures that preserve each cell’s field count, field widths, and total activity nevertheless remove the population oscillation: these controls preserve marginal single-cell statistics but destroy cross-cell phases.

The same argument applies to equal-Poisson and gamma-Poisson resampling. Matching the count distribution determines how many fields each cell has, but independent placement sets the expected cross-cell term to zero. A population oscillation surviving beyond these controls therefore reflects organisation of field locations across cells, rather than the field-count distribution alone.

## Footnotes

1 We intentionally use the notation *μ* both for the complement of the discount factor and for the high-level policy. This overloading is harmless because the two meanings occupy distinct mathematical roles: *μ* denotes a scalar in the former case and a policy in the latter.

2 Without loss of generality, we assume uniform sampling from a finite task set, i.e., 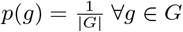.

3 For simplicity, we assume all lobes within the same atom have equal weights, *w*_*m*_.

4 The validity of the assumption is provided by the fact that 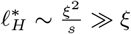, hence the HSR yields an enormous spatial horizon.

## References

[1] Tolman, E. C. Cognitive maps in rats and men. Psychological review 55, 189 (1948).

[2] O’Keefe, J. & Nadel, L. The hippocampus as a cognitive map (Oxford university press, 1978).

[3] Stachenfeld, K. L., Botvinick, M. M. & Gershman, S. J. The hippocampus as a predictive map. Nature neuroscience 20, 1643–1653 (2017).

[4] Dayan, P. Improving generalization for temporal difference learning: The successor representation. Neural computation 5, 613–624 (1993).

[5] O’Keefe, J. & Dostrovsky, J. The hippocampus as a spatial map: preliminary evidence from unit activity in the freely-moving rat. Brain research (1971).

[6] Mehta, M. R., Quirk, M. C. & Wilson, M. A. Experience-dependent asymmetric shape of hippocampal receptive fields. Neuron 25, 707–715 (2000).

[7] Schapiro, A. C., Turk-Browne, N. B., Norman, K. A. & Botvinick, M. M. Statistical learning of temporal community structure in the hippocampus. Hippocampus 26, 3–8 (2016).

[8] Lambert, M. S., Quy, R. J., Smith, R. H. & Cowan, D. P. The effect of habitat management on home-range size and survival of rural norway rat populations. Journal of Applied Ecology 45, 1753–1761 (2008).

[9] Tsoar, A. et al. Large-scale navigational map in a mammal. Proceedings of the National Academy of Sciences 108, E718–E724 (2011).

[10] Fenton, A. A. et al. Unmasking the ca1 ensemble place code by exposures to small and large environments: more place cells and multiple, irregularly arranged, and expanded place fields in the larger space. The Journal of Neuroscience 28, 11250–11262 (2008).

[11] Rich, P. D.Liaw, H.-P. & Lee, A. K. Large environments reveal the statistical structure governing hippocampal representations. Science 345, 814–817 (2014).

[12] Eliav, T. et al. Multiscale representation of very large environments in the hippocampus of flying bats. Science 372, eabg4020 (2021).

[13] Harland, B., Contreras, M., Souder, M. & Fellous, J.-M. Dorsal ca1 hippocampal place cells form a multi-scale representation of megaspace. Current Biology 31, 2178–2190 (2021).

[14] Maimon, S. R. et al. Sparse-to-dense coding transformation between hippocampal areas ca3 and ca1. Nature 1–10 (2026).

[15] Lee, J. S., Briguglio, J. J., Cohen, J. D., Romani, S. & Lee, A. K. The statistical structure of the hippocampal code for space as a function of time, context, and value. Cell 183, 620–635 (2020).

[16] Mainali, N., da Silveira, R. A. & Burak, Y. Universal statistics of hippocampal place fields across species and dimensionalities. Neuron 113, 1110–1120 (2025).

[17] Zhang, H., Rich, P. D., Lee, A. K. & Sharpee, T. O. Hippocampal spatial representations exhibit a hyperbolic geometry that expands with experience. Nature Neuroscience 26, 131–139 (2023).

[18] Mathis, A., Herz, A. V. & Stemmler, M. Optimal population codes for space: grid cells outperform place cells. Neural computation 24, 2280–2317 (2012).

[19] Sutton, R. S., Precup, D. & Singh, S. Between mdps and semi-mdps: A framework for temporal abstraction in reinforcement learning. Artificial intelligence 112, 181–211 (1999).

[20] Botvinick, M. M., Niv, Y. & Barto, A. G. Hierarchically organized behavior and its neural foundations: A reinforcement learning perspective. cognition 113, 262–280 (2009).

[21] Balaguer, J., Spiers, H., Hassabis, D. & Summerfield, C. Neural mechanisms of hierarchical planning in a virtual subway network. Neuron 90, 893–903 (2016).

[22] AbdelRahman, N. Y. et al. Composing trajectories for rapid inference of navigational goals. Neuron (2026).

[23] Yu, C. & Lengyel, M. Hierarchical successor representation for robust transfer. In Forty-third International Conference on Machine Learning (2026).

[24] Watkins, C. J. & Dayan, P. Q-learning. Machine learning 8, 279–292 (1992).

[25] Sutton, R. S., Barto, A. G. et al. Reinforcement learning: An introduction, vol. 1 (MIT press Cambridge, 1998).

[26] Gershman, S. J. The successor representation: its computational logic and neural substrates. Journal of Neuroscience 38, 7193–7200 (2018).

[27] Machado, M. C. et al. Eigenoption discovery through the deep successor representation. In International Conference on Learning Representations (2018).

[28] Drai, D., Kafkafi, N., Benjamini, Y., Elmer, G. & Golani, I. Rats and mice share common ethologically relevant parameters of exploratory behavior. Behavioural brain research 125, 133–140 (2001).

[29] Schapiro, A. C., Rogers, T. T., Cordova, N. I., Turk-Browne, N. B. & Botvinick, M. M. Neural representations of events arise from temporal community structure. Nature neuroscience 16, 486–492 (2013).

[30] Thompson, L. T. & Best, P. Place cells and silent cells in the hippocampus of freely-behaving rats. The Journal of neuroscience 9, 2382–2390 (1989).

[31] Ziv, Y. et al. Long-term dynamics of ca1 hippocampal place codes. Nature neuroscience 16, 264–266 (2013).

[32] Giusti, C., Pastalkova, E., Curto, C. & Itskov, V. Clique topology reveals intrinsic geometric structure in neural correlations. Proceedings of the National Academy of Sciences 112, 13455–13460 (2015).

[33] Ratcliffe, J. G. Foundations of hyperbolic manifolds (Springer, 2006).

[34] Momennejad, I. & Howard, M. W. Predicting the future with multi-scale successor representations. BioRxiv 449470 (2018).

[35] Brunec, I. K. & Momennejad, I. Predictive representations in hippocampal and prefrontal hierarchies. The Journal of Neuroscience 42, 299–312 (2022).

[36] Kjelstrup, K. B. et al. Finite scale of spatial representation in the hippocampus. Science 321, 140–143 (2008).

[37] Piray, P. & Daw, N. D. Linear reinforcement learning in planning, grid fields, and cognitive control. Nature communications 12, 4942 (2021).

[38] Stensola, H. et al. The entorhinal grid map is discretized. Nature 492, 72–78 (2012).

[39] El-Gaby, M. et al. A cellular basis for mapping behavioural structure. Nature 636, 671–680 (2024).

[40] Ribas-Fernandes, J. J. et al. A neural signature of hierarchical reinforcement learning. Neuron 71, 370–379 (2011).

[41] Fang, C., Aronov, D., Abbott, L. & Mackevicius, E. L. Neural learning rules for generating flexible predictions and computing the successor representation. elife 12, e80680 (2023).

[42] George, T. M., de Cothi, W., Stachenfeld, K. L. & Barry, C. Rapid learning of predictive maps with stdp and theta phase precession. Elife 12, e80663 (2023).

[43] Pfeiffer, B. E. & Foster, D. J. Autoassociative dynamics in the generation of sequences of hippocampal place cells. Science 349, 180–183 (2015).

[44] Widloski, J. & Foster, D. J. Replay without sharp wave ripples in a spatial memory task. Nature Communications 16, 10287 (2025).

[45] McNamee, D. C., Stachenfeld, K. L., Botvinick, M. M. & Gershman, S. J. Flexible modulation of sequence generation in the entorhinal–hippocampal system. Nature neuroscience 24, 851–862 (2021).

[46] Ji, Z. et al. Dynamical modulation of hippocampal replay through firing rate adaptation. Nature Communications (2026).

[47] McNamee, D., Wolpert, D. M. & Lengyel, M. Efficient state-space modularization for planning: theory, behavioral and neural signatures. Advances in neural information processing systems 29 (2016).

[48] Olshausen, B. A. & Field, D. J. Emergence of simple-cell receptive field properties by learning a sparse code for natural images. Nature 381, 607–609 (1996).

[49] Pehlevan, C. & Chklovskii, D. B. A hebbian/anti-hebbian network derived from online non-negative matrix factorization can cluster and discover sparse features. In 2014 48th Asilomar Conference on Signals, Systems and Computers, 769–775 (IEEE, 2014).

[50] Qin, S. et al. Coordinated drift of receptive fields in hebbian/anti-hebbian network models during noisy representation learning. Nature Neuroscience 26, 339–349 (2023).

[51] McNaughton, B. L., Barnes, C. A. & O’Keefe, J. The contributions of position, direction, and velocity to single unit activity in the hippocampus of freely-moving rats. Experimental brain research 52, 41–49 (1983).

[52] Markus, E. J. et al. Interactions between location and task affect the spatial and directional firing of hippocampal neurons. The Journal of neuroscience 15, 7079–7094 (1995).

[53] Hollup, S. A., Molden, S., Donnett, J. G.Moser, M.-B. & Moser, E. I. Accumulation of hippocampal place fields at the goal location in an annular watermaze task. The Journal of Neuroscience 21, 1635–1644 (2001).

[54] Dupret, D., O’neill, J., Pleydell-Bouverie, B. & Csicsvari, J. The reorganization and reactivation of hippocampal maps predict spatial memory performance. Nature neuroscience 13, 995–1002 (2010).

[55] Yaghoubi, M. et al. Predictive coding of reward in the hippocampus. Nature 651, 414–420 (2026).

[56] Maaten, L. v. d. & Hinton, G. Visualizing data using t-sne. Journal of machine learning research 9, 2579–2605 (2008).

[57] Ding, C., He, X. & Simon, H. D. On the equivalence of nonnegative matrix factorization and spectral clustering. In Proceedings of the 2005 SIAM international conference on data mining, 606–610 (SIAM, 2005).

[58] Lee, D. D. & Seung, H. S. Learning the parts of objects by non-negative matrix factorization. nature 401, 788–791 (1999).

[59] Absil, P.-A., Mahony, R. & Sepulchre, R. Optimization algorithms on matrix manifolds (Princeton University Press, 2008).

[60] Zhang, K., Ginzburg, I., McNaughton, B. L. & Sejnowski, T. J. Interpreting neuronal population activity by reconstruction: unified framework with application to hippocampal place cells. Journal of neurophysiology 79, 1017–1044 (1998).

[61] Brown, E. N., Frank, L. M., Tang, D., Quirk, M. C. & Wilson, M. A. A statistical paradigm for neural spike train decoding applied to position prediction from ensemble firing patterns of rat hippocampal place cells. The Journal of Neuroscience 18, 7411–7425 (1998).

[62] Freeman, L. C. A set of measures of centrality based on betweenness. Sociometry 35–41 (1977).

[63] Kruskal, J. B. Multidimensional scaling by optimizing goodness of fit to a nonmetric hypothesis. Psychometrika 29, 1–27 (1964).

[64] Rich, D., Liaw, H. & Lee, A. K. Extracellular CA1 activity from rats during exploration of a novel 48m track. CRCNS.org (2022). URL 10.6080/K0BV7DT2.Dataset.

## References

[1] Dayan, P. Improving generalization for temporal difference learning: The successor representation. Neural computation 5, 613–624 (1993).

[2] Chaikin, P. M., Lubensky, T. C. & Witten, T. A. Principles of condensed matter physics, vol. 10 (Cambridge university press Cambridge, 1995).

[3] Samoradnitsky, G. & Taqqu, M. S. Stable non-Gaussian random processes: stochastic models with infinite variance (Routledge, 2017).

[4] Di Nezza, E., Palatucci, G. & Valdinoci, E. Hitchhiker’s guide to the fractional sobolev spaces. Bulletin des sciences mathématiques 136, 521–573 (2012).

[5] Lee, D. D. & Seung, H. S. Learning the parts of objects by non-negative matrix factorization. nature 401, 788–791 (1999).

[6] Absil, P.-A., Mahony, R. & Sepulchre, R. Optimization algorithms on matrix manifolds (Princeton University Press, 2008).

[7] Ding, C., He, X. & Simon, H. D. On the equivalence of nonnegative matrix factorization and spectral clustering. In Proceedings of the 2005 SIAM international conference on data mining, 606–610 (SIAM, 2005).

[8] Pehlevan, C. & Chklovskii, D. B. A hebbian/anti-hebbian network derived from online non-negative matrix factorization can cluster and discover sparse features. In 2014 48th Asilomar Conference on Signals, Systems and Computers, 769–775 (IEEE, 2014).

[9] Tropp, J. A. An introduction to matrix concentration inequalities. Foundations and trends® in machine learning 8, 1–230 (2015).

[10] Ball, K., Carlen, E. A. & Lieb, E. H. Sharp uniform convexity and smoothness inequalities for trace norms. Inventiones mathematicae 115, 463–482 (1994).

[11] Kac, M. On the average number of real roots of a random algebraic equation. Bulletin of the American Mathematical Society 49, 314–320 (1943).

[12] Rice, S. O. Mathematical analysis of random noise. The Bell System Technical Journal 23, 282–332 (1944).

[13] Azaïs, J.-M. & Wschebor, M. Level sets and extrema of random processes and fields (John Wiley & Sons, 2009).

[14] Mainali, N., da Silveira, R. A. & Burak, Y. Universal statistics of hippocampal place fields across species and dimensionalities. Neuron 113, 1110–1120 (2025).

[15] Volkonskii, V. & Rozanov, Y. A. Some limit theorems for random functions. i. Theory of Probability & Its Applications 4, 178–197 (1959).

[16] Berman, S. M. Asymptotic independence of the numbers of high and low level crossings of stationary gaussian processes. The Annals of Mathematical Statistics 42, 927–945 (1971).

[17] Rényi, A. A poisson-folyamat egy jellemzése= a characterization of poisson processes. A Magyar Tudományos Akadémia Matematikai Kutató Intézetének Közleményei 1, 519–527 (1956).

[18] Daley, D. J. & Vere-Jones, D. An introduction to the theory of point processes: volume I: elementary theory and methods (Springer, 2003).

[19] Rich, P. D.Liaw, H.-P. & Lee, A. K. Large environments reveal the statistical structure governing hippocampal representations. Science 345, 814–817 (2014).

[20] Lee, J. S., Briguglio, J. J., Cohen, J. D., Romani, S. & Lee, A. K. The statistical structure of the hippocampal code for space as a function of time, context, and value. Cell 183, 620–635 (2020).

[21] Firth, D. Multiplicative errors: log-normal or gamma? Journal of the Royal Statistical Society Series B: Statistical Methodology 50, 266–268 (1988).

[22] Ratcliffe, J. G. Foundations of hyperbolic manifolds (Springer, 2006).

